# High microbial diversity in the rhizosphere and soil improve carrot (*Daucus carota* L.) postharvest storability

**DOI:** 10.64898/2025.12.02.691807

**Authors:** Sannakajsa Velmala, Taina Pennanen, Minna Haapalainen, Petteri Karisto, Satu Latvala, Minna Pirhonen, Terhi Suojala-Ahlfors

**Author notes:** **One-sentence Summary** Higher bacterial diversity, particularly in organically managed soils, alongside greater overall microbial diversity, was associated with improved postharvest storability of carrots.

## Abstract

Postharvest diseases can cause significant losses in carrot (*Daucus carota*) yield during storage. In this study, the effects of soil quality and microbial diversity in the field bulk soil and rhizosphere on the development of postharvest diseases were investigated. Microbial genera in bulk soil and rhizosphere samples were identified by Illumina Miseq amplicon sequencing of bacterial 16S and fungal ITS regions. Disease symptoms in the stored carrots were monitored after six months of cold storage, and the main pathogens were identified. The field sites with mineral soils, which had a higher pH, had a higher bacterial and fungal diversity than organic soils. The organically managed fields had a higher bacterial diversity than the conventionally managed fields. Community structure of both bacteria and fungi was largely driven by field site including its main chemical characteristics, with rhizosphere and bulk soil samples from the same site displaying comparable assemblages. Higher soil microbial diversity was associated with higher postharvest storability in the stored carrots. The weather during the growing season also had an effect on the observed fungal biodiversity, which was higher at sites that had lower temperatures and less rain prior to harvest.

## 1 Introduction

Soil health encompasses the soil’s physical, chemical, and biological condition, and determines its capacity to function as a vital living system and to provide ecosystem services (Directorate-General for Environment, 2023; Larkin, 2015). The importance of healthy soil in mitigating greenhouse gases and facilitating food production cannot be overstated. On a global scale, a considerable proportion of agricultural soils are degraded (Lal 2015), which has resulted in diminished productivity due to the weakening of soil ecosystem services (Pereira et al., 2018). This suggests that the early signs of decreased soil health, even before measurable changes in soil physical properties can be observed, may be a factor predictive of an increased incidence of diseases, especially in cases of intensive cropping.

The rhizosphere, defined as the interaction zone between plant roots and soil, including all its microbial diversity, is of critical importance for plants, as it governs their water and nutrient uptake, which is required for growth and development. The rhizosphere is the zone of direct root contact for beneficial microorganisms, including plant growth-promoting bacteria and symbiotic mycorrhizal fungi (Raaijmakers et al., 2009). In addition to beneficial microbes, the rhizosphere microbiome can also contain pathogens (Mendes et al., 2013; Mesny et al., 2023). The structure and functionality of the rhizosphere microbial populations are also shaped by the plant species and genotype and by the soil type (Berg & Smalla, 2009). It has been suggested that plants actively recruit beneficial microorganisms to improve their resistance against various stress factors (Mesny et al., 2023). In disease suppressive soils, the antimicrobial compounds produced by rhizosphere microorganisms suppress pathogen populations in the soil (Raaijmakers et al., 2009). Moreover, the beneficial microorganisms can enhance the plant defence system against pathogens. Recent studies have demonstrated that even in soils which are chemically and structurally relatively similar, the composition of soil microbiota can have a significant impact on crop health and yield (Ali et al., 2023).

Plants interact with a plethora of different microorganisms that colonize all the plant surfaces (Mesny et al., 2023). Bacterial species that have previously been linked to the suppression of disease are found in several phyla, including Pseudomonadota (synonym Proteobacteria, new phylum names by Oren et al. (2021)), Bacillota (syn. Firmicutes), and Actinomycetota (syn. Actinobacteria) (Kinkel et al., 2012; Mendes et al., 2011). Moreover, both biotic and abiotic environmental factors induce changes in the prevalence of different microbial species that may suppress or promote plant diseases. Vegetable decay during storage is frequently attributable to the interaction of decreased biodiversity and augmented proportions of pathogenic species (Kusstatscher et al., 2020).

Carrot (*Daucus carota* subsp. *sativa*) is a globally important field vegetable, with a seventh-highest production volume (Que et al., 2019). In the Northern Europe, where the cultivation of field vegetables is confined to a single annual growing season, most of the carrot yield is typically stored prior to marketing. However, during the storage period, which may even last for six months, a significant proportion of the yield is lost, mostly due to deterioration by pathogenic microbes. As demonstrated in the research conducted in Finland by Latvala et al. (2024) the postharvest storage losses of carrot due to decay can be approximately 20% even in well-controlled storage conditions. An earlier study in Norway by Hermansen et al. (2012) gave an estimate of losses between 20% and 50%, bringing considerable economic losses along.

Many fungal genera, including plant pathogens, such as *Rhizoctonia* and *Fusarium*, are predominant members of the core mycobiome of carrot (Abdelrazek et al., 2020). Both the wounds caused by harvesting, and the water loss during extended postharvest storage render the carrot root more vulnerable to pathogens. In Finland, the predominant pathogenic fungi responsible for postharvest decay of carrots are *Mycocentrospora acerina*, causing liquorice rot, *Botrytis cinerea*, causing grey mould, and *Fusarium* spp., which are often considered secondary pathogens (Latvala et al., 2024; Suojala, 1999). Furthermore, different bacterial strains of the genera *Pseudomonas*, *Pectobacterium* and *Erwinia* have also been shown to cause decay of carrots during storage (Kahala et al., 2012). In sugar beets (*Beta vulgaris* L.), the core microbiome was found to differ between healthy and diseased plants, with high proportions of antagonistic bacteria providing resistance to deleterious shifts in the fungal microbiome (Kusstatscher et al., 2019). Thus, the plant microbiome was proposed to be the key element in biological control of postharvest pathogens and storability of root vegetables (Kusstatscher et al., 2020).

Despite the extensive research on plant-associated microbial communities, their role in carrot health during storage remains unclear. The present study investigates the influence of soil and rhizosphere microbes on the storability and postharvest diseases of carrot. We hypothesised that 1) both beneficial and potentially pathogenic bacteria and fungi are found in the rhizosphere and bulk soil surrounding the carrot roots, and 2) a high microbial diversity suppresses postharvest diseases, and 3) a good storability is associated with high abundance of beneficial microbial groups. A study was conducted in which samples were collected from 26 carrot fields in Finland. The samples were analysed to assess soil characteristics and bacterial and fungal diversity, using sequencing of the 16S and ITS2 barcode regions. The storage losses of carrots grown in these fields were monitored over half a year of cold storage. Furthermore, research was conducted on whether the initial indications of deteriorating soil health were predominantly associated with biological factors as opposed to physical or chemical characteristics.

## 2 Material and methods

### 2.1 Carrot and soil sampling

Carrot and soil samples were taken between mid-September to early October in 2020 and 2021 from 14 and 12 field sites, respectively (Table 1; Supplementary Figure S1; A to Z). All the field sites sampled in 2021 were different from the sites in 2020. In both years three of the field sites were organically managed. Sampling was performed on six replicate areas per field. The distance between separate sampling areas was 30-100 m, depending on the size of the field. The sampling area consisted of a single carrot row, five meters in length. The fields utilised in this study are authentic commercial carrot production fields, reflecting authentic soil types and management practices. This approach guarantees that the results represent genuine production conditions; however, it does not allow for completely balanced design.

**Table 1.**
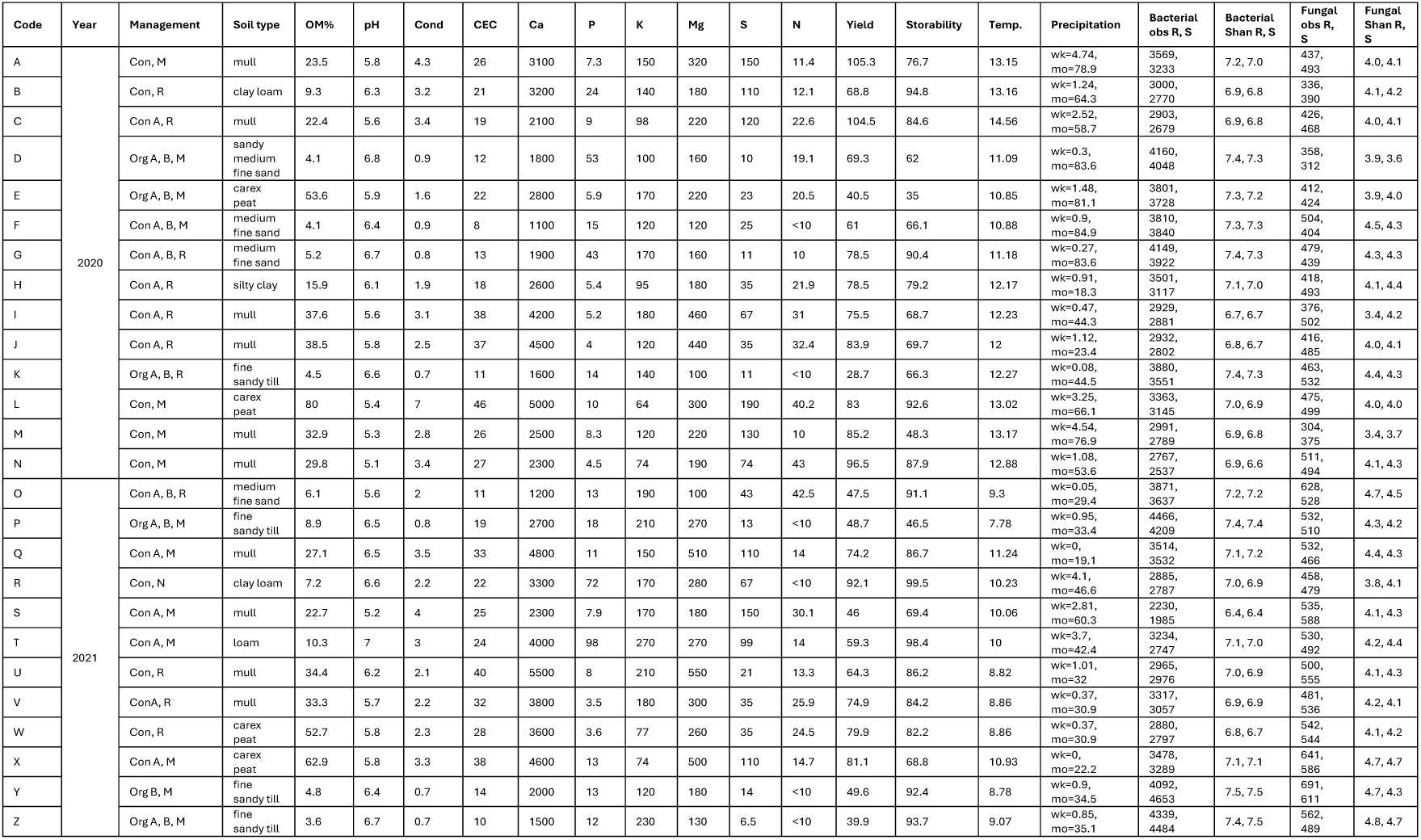
Site characteristics across 26 carrot fields, including management (farm type Con/Org, carrot in history (A), green manure used (B), variety (Maestro M, Romance R, Newhall N)), soil properties (electrical conductivity (10 mScm^-1^), cation exchange capacity (cmol/kg), nutrient concentrations (mg/l)), weather (mean temperature in the month before sampling; mean rainfall in the week and month before harvest), and microbial diversity (richness and Shannon index) in rhizosphere and bulk soil.

Soil samples for DNA analysis of rhizosphere and bulk soil were collected first. Rhizosphere soil was sampled from ten sampling points per area, by lifting two carrots at 50 cm distance from each other. Rhizosphere soil was gently brushed from these 20 carrots to a zip-lock bag with a clean disposable toothbrush. The same toothbrush was used for all carrots within the same sampling area, and the soil detached from the carrots and the toothbrush was frozen at -20 °C within 24 h. Altogether, six replicate rhizosphere samples were taken per field. In total 156 rhizosphere samples were collected.

Bulk soil was sampled by taking six replicate soil cores (distance 1 m, diameter 25 mm, depth 0-10 cm) from each sampling area in clean zip-lock bags, mixed carefully, and divided into two subsamples: one for analysis of soil properties and the other for DNA analysis. The latter sample set was frozen at -20 °C within 24 h. Altogether six replicate bulk soil samples were taken per field. The soil borer was washed with soap and tap water between the fields and wiped clean with ethanol between the replicate sampling sites.

Additional soil samples (depth 0-25 cm, 400 ml) were taken for soil nutrient analysis. Soil analysis was performed on a composite sample, which was formed by mixing the six subsamples from all six sampling areas of the same field.

After soil sampling, all the carrots within the sampling area were collected to a plastic container, and the carrot tops were taken off. Altogether, six replicate 8-kg carrot samples were taken per field and packed into woven polypropylene bags, separately for each sampling area. Bags were transferred to the cold store and placed in 1 m^3^ plastic containers for the storage experiment, at 0-1 °C and 90% relative humidity, in darkness.

### 2.2 Carrot yield and storage

Carrot yield was calculated individually for each sampling area by dividing the total weight of carrot roots by the area sampled (Table 1).

The storage loss of carrots was analyzed in mid-March after approximately 5.5-months storage time. The cold-stored carrots, six 8-kg samples per field, were divided into healthy and diseased carrots based on visual symptoms. The carrot postharvest storability (as percentage), was calculated as the ratio of the weight of healthy carrots to the total weight of the lot, multiplied by 100 (Table 1). In March, the mean and median storability percentages were recorded as 78% and 82%, respectively. For downstream analyses that use categorical explanatory variables (i.e., differential abundance analysis), the samples were divided into two groups: high (>80%) and low (≤80%) storability.

### 2.3 Soil analysis

Soil analysis and measurement of cation exchange capacity (CEC) was performed in the laboratory of Eurofins Finland. The analysis included the sensory analysis of soil type, measurements of electrical conductivity (ISO 11265: 1994, modified), pH in water (ISO 10390:2005, modified), macro nutrients determined by the acid ammonium acetate extraction (Vuorinen & Mäkitie, 1955), soluble nitrogen (SFS-EN 13654-1:2002), and organic matter (OM%) (SFS-EN 15935:2012). Soil with an OM content of 20% or more was considered to be organic, while soil with an OM content of less than 20% was considered to be mineral.

### 2.4 Climate data

For assessing the daily mean temperature and the rain sum of the growing season we used 1*1 km grate-type data from the Finnish Meteorological Institute (Table 1). To simplify climate parameters, we calculated their mutual correlations. Mean temperature of week, fortnight and month prior sampling were positively correlated (r>0.8, *P*<0.05) (Supplementary Figure S2), and thus *monthly mean temperature* was chosen to be used as the representative descriptive parameter. Also, the total sum of rain during the growing season had high negative correlation (r= -0.8, *P*<0.05) with monthly mean temperature (Supplementary Figure S2). Additional parameters included the average rainfall during the week before harvest (*preharvest week mean of rainfall*) and the average rainfall during the month before harvest (*preharvest month mean of rainfall*).

### 2.5 DNA extraction and sequencing

Frozen rhizosphere and bulk soil samples were melted and carefully mixed, and a 250 mg subsample of soil was taken with a sterile spatula for DNA extraction with DNeasy Power soil Pro (Qiagen). Lysis buffer, 800 µl, was added on the samples, followed by short mixing by vortex genie 2 (Scientific industries) and 10 min incubation at 65 °C. Samples were disrupted after incubation with TissueLyzer II (Qiagen) at 25 Hz, 2 x 5 min and then shaken horizontally in a bead tube holder (Macherey-Nagel) for 15 min. Finally, the samples were centrifuged for 2 min 15 000 x g, and 600 µl of supernatant was transferred to DNA extraction by QIAcube Connect, following manufacturer’s instructions (Qiagen). DNA was eluted to 100 µl elution buffer. In total 312 rhizosphere and soil DNA samples were extracted and stored -20 ° C.

Amplification of DNA and amplicon sequencing (Illumina MiSeq V3, paired end 2 × 301 bp, 2x10bp dual index, MCS 2.5.0.5/2.6.2.1 and RTA 1.18.54.0) were performed at the Institute of Genomics, University of Tartu, Estonia. For amplification of the bacterial 16S V4 region, primers 515 and 806 (Caporaso et al., 2012) were used, and for fungal internal transcribed spacer region ITS2, primers gITS7 (Ihrmark et al., 2012) and ITS4 (White et al., 1990) were used.

### 2.6 Bioinformatics

Bioinformatic analyses were done with PipeCraft 2.0 (Anslan et al., 2017), pipecraft2-manual.readthedocs.io/en/stable, with OTU workflow that utilises cutadapt 3.5 (Martin, 2011), fqgrep 0.4.4 (Das, 2011), and vsearch 2.18.0 (Rognes et al., 2016) software, with default parameters with following exceptions: re-orient and cut primers (2 mismatches), reads were merged, and quality filtered with minimum overlap 15 and length 150 bp, max differences 99, and Maxee: 1, MaxN: 0. De novo filtering was used for chimeras with threshold of 0.97. In addition, fungal ITS2 region was extracted from reads with ITSx v1.1.3 (Bengtsson-Palme et al., 2013) and mothur v1.46.1 (Schloss et al., 2009), and then sequence reads were clustered and OTU table created with threshold 0.97.

Annotation of OTU sequences was done with BLAST v2.11.0+ (Camacho et al., 2009) against databases SILVA SSU 138 (Quast et al., 2012) for bacteria and UNITE (sh_general_release_dynamic_10.05.2021.fasta) (Abarenkov et al., 2021) for fungi with the parameters: word size = 7; e = 0.001; reward = 1; penalty = −1; gap open = 1; gap extend = 2. During secondary filtering, OTUs without any blast result, identity less than 70% to bacteria or fungi, and e-value higher than 1E-24 were removed. Bacteria with the same annotation were consolidated. The most abundant bacterial OTUs in the dataset originated from the crop plant itself: carrot chloroplast and mitochondria with 115 706 (1.7%) and 30 101 (0.4%) reads respectively. Also 126 other mitochondria originating OTUs were found, in total 35 928 reads (0.5%). The plant mitochondrial and chloroplast OTUs were excluded from further analysis. For fungi, OTUs with the same *Species hypothesis code* (Kõljalg et al., 2013) were consolidated. Lastly, OTUs with less than 5 reads were removed. This resulted in 14 947 bacterial and 3947 fungal OTUs. The median library size per sample was 22 150 reads for bacterial 16S region (Supplementary Figure S3) and 16 775 reads for fungal ITS region (Supplementary Figure S4). As the smallest library size differed from the median less than 3-fold, no samples were removed from the analysis as outliers.

Alpha-diversity indices were calculated from raw OTU data, which was then normalized for further analyses using the geometric mean of pairwise ratios (GMPR) method (Chen et al., 2018) which deals with zero-inflated community data and preserves differences in relative abundances of taxa.

### 2.7 Data analysis

Data was analysed with R 4.4.1. (R Core Team, 2024) phyloseq library (McMurdie & Holmes, 2013), and visualized with ggplot2 (Wickham, 2016) and. Bacterial OTUs are reported on both phylum and family levels, and fungal OTUs are reported on both phylum and genus levels. This is because the barcoding regions and the available databases do not separate lower taxonomical levels well (O’Donnell et al., 2022).

Pearson correlation coefficients between the measured parameters were calculated with cor function, library (corrplot (Wei & Simko, 2024)), using a significance level of 0.05 for all the statistical tests. Factor analysis of mixed data (FAMD) was done with libraries FactoMineR (Lê et al., 2008), vcd and factoextra (Kassambara & Mundt, 2020), to study associations between both quantitative and qualitative variables using principal component analysis for quantitative variables, and multiple correspondence analysis for qualitative variables (Supplementary Figures S7, S8). A linear model was used to study the relationship of use of green manure and storability.

Alpha-diversity indexes (e.g., richness, Shannon, chao1 estimates) were calculated with microbiome package (Lahti & Shetty, 2019). We also calculated the richness from a rarefied with resubmission data, which resulted in lower number of OTUs but with the same rank order of fields (data not shown). Beta-diversity was investigated with non-metric multidimensional scaling (NMDS) with library vegan (Oksanen et al., 2020). Envfit function was used to fit environmental parameters to the ordination. Permutational multivariate analysis of variance using distance matrices was assessed with adonis2 function. Betadisper function was used to determine if there are significant differences in the dispersion of multivariate data among groups. Core taxa, microbiome library (Lahti & Shetty, 2019), represent OTUs that are characteristic of a sample type, here with 90% prevalence. This returns the taxa that exceed the given prevalence and detection thresholds.

To study predominance of certain OTUs in high and low storability carrots we cross-correlated the storability with GMPR normalized OTU data input matrices (library microbiome, function associate), using Spearman correlation.

Differential abundance analysis of microbial taxa between experimental groups was performed using the DEseq2 (Love et al., 2014) package in R. DESeq2 utilizes a negative binomial distribution model to detect differences in read counts, accounting for both the mean-variance relationship inherent to count data and overdispersion. The method internally performs normalization to correct for differences in sequencing depth and library composition between samples, thus eliminating the need for pre-normalization of raw counts.

The effects of farm management type, and the use of green manure, on microbial alpha diversity and richness were studied using a linear mixed model from the nlme package (Pinheiro et al., 2023), with the field site as a random factor. Analyses were performed separately for the rhizosphere and the bulk soil.

### 2.8 Data storage and availability

All the data will be made available upon publication. Raw sequences with metadata and attributes are available in NCBI GenBank BioProject PRJNA976070 with accessions SAMN35358255-SAMN35358630. Other data is stored in Zenodo **10.5281/zenodo.17212592.**

## 3 Results

### 3.1 Soil properties, other environmental factors, carrot storability and their correlations

Soil properties, management, cultivar, and location information for the studied carrot field sites are presented in Table 1 and Supplementary Figure S1. In five out of six organically managed fields carrot had been grown earlier and they all used green manure in rotation. In eight out of 20 conventionally managed fields carrot had not been grown before, and in four, carrots had been grown in the last three years (Table 1). Only three conventional field sites, all in which carrot had been grown before, used green manure in rotation. Soil type varied from fine sandy till and medium fine sand to fine clay-rich loam and finally to organic mull and peat (Table 1). Soil types were characterized by distinctive organic matter content (OM%), varying from 4 to as high as 80. Here OM% is used to describe the variation between soil types.

Carrot storability had weak negative association with OM% (*r*=-0.2, *P*<0.05, Supplementary Figure S2) and carrots from the three loamy soils, mineral soil type, showed the highest storability (Table 1). Soil phosphorus content had a weak positive correlation with storability (*r*=0.3., *P*<0.05). Other soil chemical characteristics were not associated with carrot storability (Supplementary Figure S2). Soil pH varied across fields from 5 to 7 (Table 1). Soil pH had a strong positive correlation with phosphorus content (*r*=0.6., *P*<0.05), and a strong negative correlation with conductivity (*r*=-0.6, *P*<0.05), nitrogen content (*r*=-0.7, *P*<0.05), and OM% (*r*=-0.6, *P*<0.05). *Monthly mean temperature* of growing season had a very strong negative correlation with the total sum of rain during the growing season (*r*=-0.8, *P*<0.05), and a positive correlation with the total carrot yield (*r*=0.6, *P*<0.05). Carrot storability had a weak negative association with mean rainfall prior to sampling (*r*=-0.3, *P*<0.05).

The associations of both quantitative and qualitative variables were further explored with factor analysis of mixed data (FAMD) that considers all the dependent variables together (Supplementary Figure S5). FAMD separated the sites according to soil type, management practises (farm type and use of green manure), carrot cultivation history, and carrot cultivar. The first dimension separated sites according to conductivity, soil type (mineral vs. organic), pH, sulphur content, CEC, OM%, farm type (conventional, organic), and carrot yield. Sites with organic soil type had higher conductivity, CEC, OM%, sulphur and nitrogen content and low pH (5.7). On the contrary, sites on mineral soil had higher pH (6.5, Table 1). The second dimension separated sites according to average rainfall during the preceding week before harvesting, phosphorus content, carrot cultivar and cultivation history.

Sites with high phosphorus, Newhall carrot variety and rainy weather clustered together. In the FAMD analysis carrot storability was colinear with the second dimension but had a low contribution to it. Thus, the storability of carrots is only weakly associated with the parameters contributing to the second dimension. Moreover, the use of green manure did not significantly associate with storability (*P*>0.17) even though the average storability was higher in sites where no green manure was used (Table 1).

### 3.2 Characteristics of microbial data in relation to field site, management, weather, and soil properties

#### 3.2.1 Microbial community composition, diversity and species richness

##### The main microbial phyla in rhizosphere and bulk soil

Based on the 16S sequence data, the most abundant bacterial phyla in the rhizosphere and bulk soil samples, respectively, were Pseudomonadota (26%, 24%), Actinomycetota (17%, 19%), and Acidobacteriota (13%, 15%) (Figure 1 a, (Oren et al., 2021, 2022)). Based on the ITS2 sequence data, the most abundant fungal phyla in both the rhizosphere and the bulk soil communities were Ascomycota (42%, 41%), Basidiomycota (28%, 24%), and Mortierellomycota (18%, 23%) (Figure 1 b).

**Figure 1.**
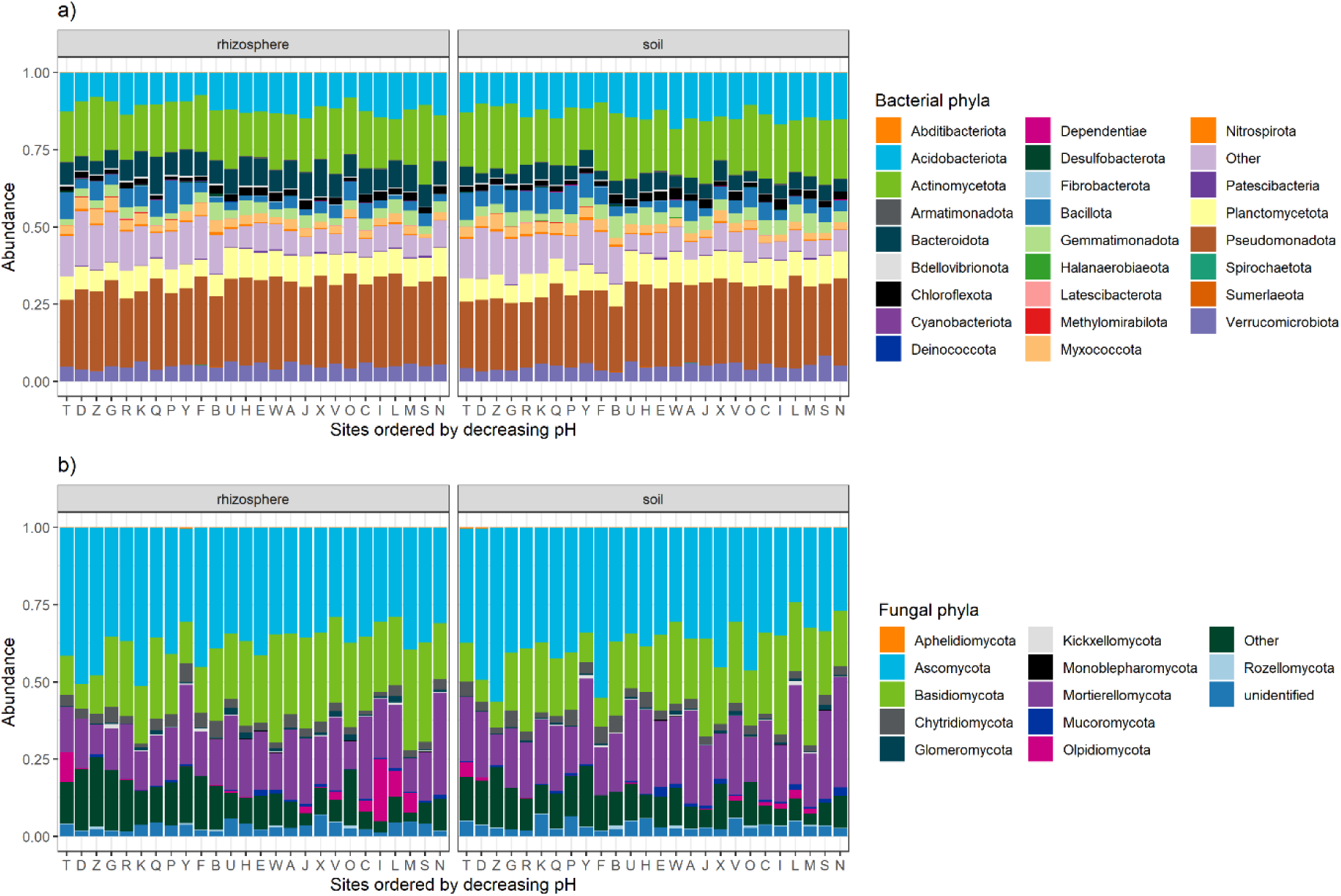
Relative abundance of main bacterial and fungal phyla. X-axis sites ordered by decreasing pH. The figure displays the dominant bacterial phyla across sample types and sites, with rare phyla grouped into the category “Other” for clarity. Colours represent different phyla as indicated in the legend. Data are presented as relative proportions. Pseudomonadota (syn. Proteobacteria), Actinomycetota (syn. Actinobacteria), Chloroflexota (syn. Chloroflexi), Bacillota (syn. Firmicutes), Cyanobacteriota (syn. Cyanobacteria) (Oren et al., 2021, 2022).

##### Microbial diversity in rhizosphere and bulk soil

All bacterial and fungal OTUs were present in both rhizosphere and bulk soil samples (14 947 bacterial and 3 947 fungal OTUs). The mean observed bacterial OTU richness in a sample was only marginally significantly higher in the rhizosphere than in the bulk soil (*P*=0.053), 3 424 and 3 279 OTUs respectively (Table 1, Supplementary Figure S7). The lowest bacterial OTU richness, 1 985, was found in the conventionally managed field S with acidic organic bulk soil (low pH, high nitrogen content, Table 1), and the highest, 4 653, in the organically managed field Y with mineral rhizosphere soil (high pH, low nutrients) (Table 1). The median observed fungal OTU richness was similar in rhizosphere and in bulk soil, 477 and 498 respectively (Supplementary Figure S7). The lowest fungal OTU richness was 312, found in organically managed mineral bulk soil (field D, high pH, low nutrients) and the highest, 691, was again in the organically managed mineral rhizosphere soil (field Y, high pH, low nutrients) (Table 1).

The average bacterial Shannon diversity was 7 in both rhizosphere and bulk soil samples (*P*=0.056); it was the lowest in conventionally managed acidic organic bulk soil (field S) and the highest in organically managed mineral bulk soils, fields Y and Z, with high pH and low nitrogen content (Table 1, Figure 2 a). Soil pH and bacterial alpha diversity correlated positively; The sites that had higher pH, on mineral soil, also showed higher bacterial diversity (*r*>0.5, *P*<0.05, Supplementary Figure S2). Both the bacterial diversity and the observed species richness correlated negatively with soil conductivity, sulphur content and CEC (*r*<-0.5, *P*<0.05, Supplementary Figure S2).

**Figure 2.**
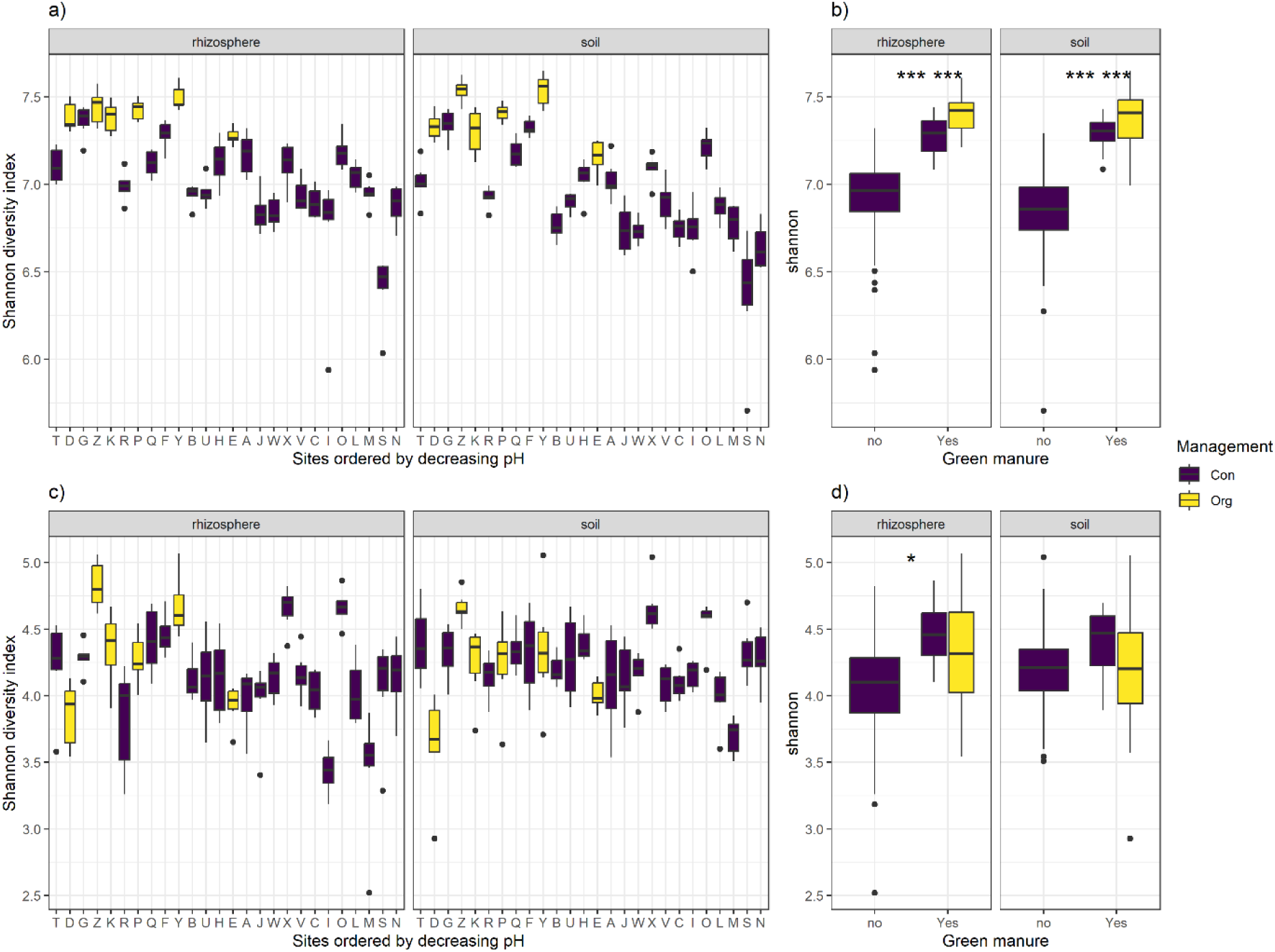
Shannon diversity indices of bacterial (a, b) and fungal (c, d) OTUs across different sample types. Boxplots represent the median and interquartile range. Diversity values (a, c) grouped by sites, ordered by decreasing soil pH. Diversity values (b, d) grouped by green manure use and management type. There were statistically significant differences between b) both management and green manure use types p<0.001 ***, and d) between sites that use green manure p<0.01 *.

Bacterial OTU richness and Shannon diversity was statistically significantly higher (*P*<0.001) in the organically managed sites than in the conventionally managed sites in both rhizosphere and bulk soil (Figure 2 b, Supplementary Figure 7). Likewise, the use of green manure was associated to higher (*P*<0.001) bacterial OTU richness and Shannon diversity in both rhizosphere and bulk soil (Figure 2 b).

The fungal Shannon diversity was on average 4.2 in both the rhizosphere and bulk soil samples (*P* >0.1); it was the lowest in conventionally managed acidic organic rhizosphere soil, field I (low pH, high nitrogen content) and the highest in organically managed mineral rhizosphere soil, field Z (high pH, low nutrients) (Table 1, Figure 2 c). There was no strong linear association between the soil physicochemical parameters and the fungal biodiversity. Fungal Shannon diversity and OTU richness correlated negatively (*r*<-0.3, *P*<0.05) with temperature and rainfall: the diversity was higher in sites that had lower temperatures and less rain preceding the harvest (Supplementary Figures S2, S5).

Management type did not affect the diversity (Figure 2d). However, the Shannon diversity in the rhizosphere was significantly higher (*P*<0.05) in sites that had use green manure in rotation (Figure 2d, Supplementary figure S8).

##### Core taxa and drivers of community composition in rhizosphere and bulk soil

The core bacterial and fungal OTUs in both the rhizosphere and the bulk soil were representative of the main phyla and were mainly shared (324 bacterial OTUs and 67 fungal OTUs) between both sample types (Figure 1, Supplementary Table 1). In addition, 72 bacterial OTUs were characteristic to only rhizosphere and 38 to bulk soil. There were 57 fungal core OTUs found in rhizosphere and 56 in bulk soil, from which 46 were shared between the sample types (Supplementary Table 1).

Both bacterial and fungal communities grouped according to the field, and the rhizosphere and bulk soil samples of each site shared similar microbial communities (Figures 3 a and 3 c). The fields with high OM% and low pH clustered tightly together, and so did the mineral field sites with a higher pH. Thus, the soil type and its main chemical characteristics pH, OM%, and phosphorus and nitrogen content appear to be the primary drivers of bacterial and fungal communities, respectively, along the first axis (Figures 3a-3d, Supplementary Table 2). Variation in *Preharvest week mean of rainfall*, soil CEC, conductivity, other macronutrients K, S as well as the use of green manure, especially for fungi, aligned with the 2^nd^ axis and significantly correlated with the ordination (Figures 3 a-3d, Supplementary Table 2). Furthermore, the organically managed sites clustered tightly together and shared similarities especially in their bacterial communities. Carrot storability, aligned with the 2^nd^ axis, had a weak but significant correlation with the ordination for both bacteria and fungi (Supplementary Table 2).

**Figure 3.**
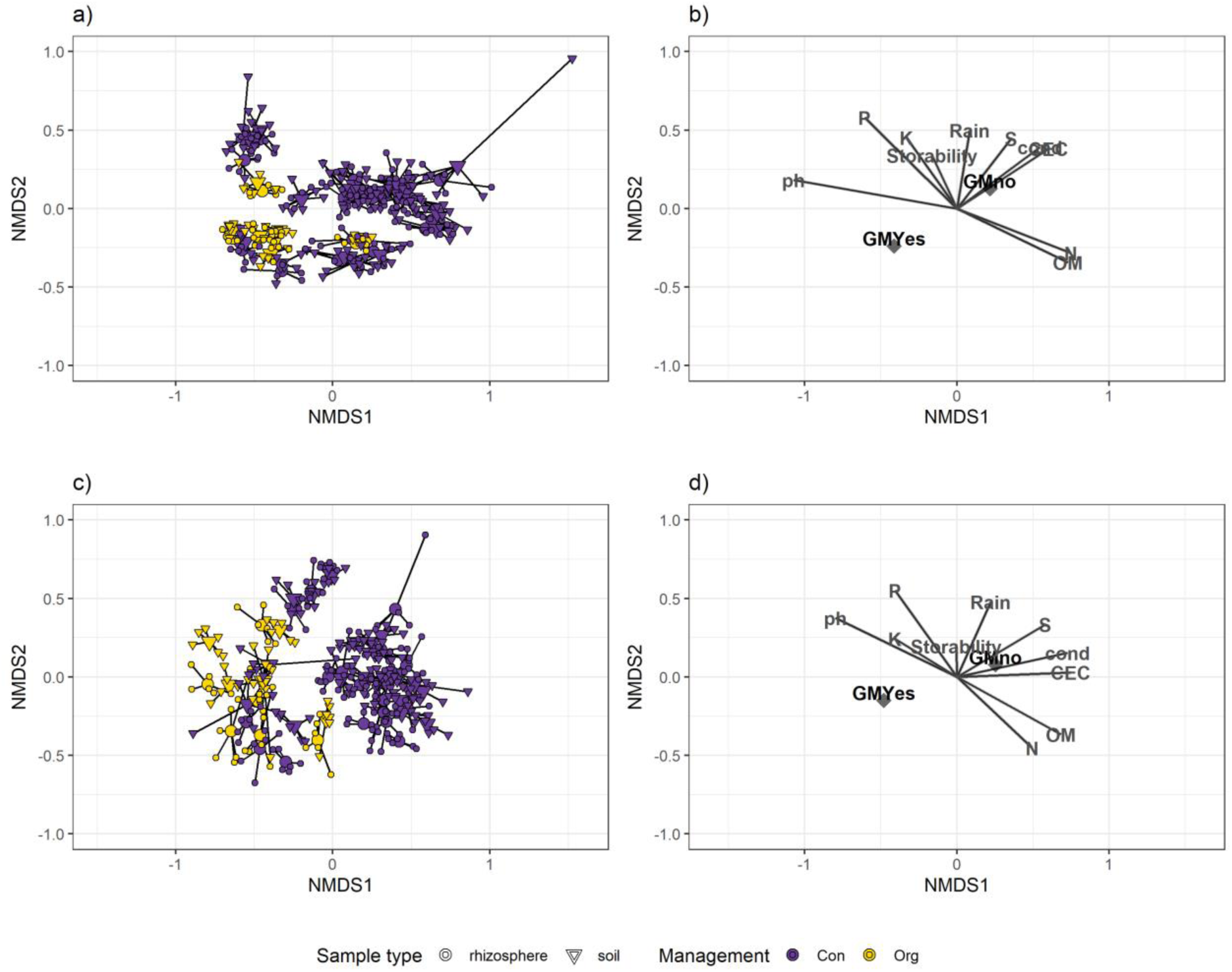
A 2D non-metric multidimensional scaling NMDS of a) bacterial and c) fungal community highlighting the sites, sample types (○ rhizosphere, Δ bulk soil) and management (conventional (purple) vs. organic (yellow)), and showing the soil characteristics (OM%, pH, P, K, S, N, CEC, rain, storability, green manure, conductivity) and carrot storability and fitted as vectors in the same ordination (b, d). Environmental factors unit length vectors are scaled by their correlation (p<0.05).

The permutational ANOVA supports the conclusions based on NMDS visualization (Table 2); The main factor explaining the variation in the bacterial and fungal communities was the field site (R^2^=0.61 and R^2^=0.59, *P* <0.001). By using the field as a grouping factor, we calculated the marginal effects of other parameters: OM% and carrot cultivar, and the use of green manure in rotation and all the other measured parameters together explained less than half of the variation in the bacterial and fungal communities (Table 2). However, there was a significant dispersion among the fields and thus the data may slightly overestimate the field effect.

**Table 2.** Results of PERMANOVA (adonis2) analysis on bacterial 16S rRNA and fungal ITS data using "field" as the grouping variable. The table presents the partitioning of multivariate variation in microbial community composition This analysis tests for significant differences in community structure among fields based on Bray-Curtis dissimilarities, with significance assessed via permutations. By margin, field as grouping variable. CCH carrot cultivation history.

|  | Bacteria |  |  | Fungi |  |  |
| --- | --- | --- | --- | --- | --- | --- |
|  | R<sup>2</sup> | F | Pr(>F) | R<sup>2</sup> | F | Pr(>F) |
| Sample type: Rhi - Bulk soil | 0.025 | 13.68 | 0.001 | 0.023 | 11.80 | 0.001 |
| P | 0.016 | 8.60 | 0.069 | 0.018 | 8.96 | 0.001 |
| N | 0.010 | 5.29 | 0.001 | 0.010 | 5.11 | 0.001 |
| pH | 0.014 | 7.87 | 0.075 | 0.012 | 6.13 | 0.001 |
| OM% | 0.026 | 14.44 | 0.001 | 0.024 | 11.95 | 0.001 |
| Carrot storability | 0.015 | 8.22 | 0.001 | 0.017 | 8.32 | 0.001 |
| Farm type: Con - Org | 0.013 | 7.37 | 0.059 | 0.019 | 9.52 | 0.001 |
| Carrot cultivar | 0.034 | 9.37 | 0.001 | 0.040 | 10.06 | 0.001 |
| CCH: Recent-Yes-No | 0.022 | 6.07 | 0.078 | 0.032 | 8.15 | 0.001 |
| Green manure: Yes - No | 0.036 | 19.81 | 0.001 | 0.040 | 20.32 | 0.001 |
| Rainfall of preharvest week | 0.012 | 6.84 | 0.001 | 0.012 | 6.14 | 0.001 |
| Residual | 0.54 |  |  | 0.59 |  |  |
| Total | 1.00 |  |  | 1.00 |  |  |

F, pseudo-F statistic; Pr, permutation-based *P*-values

### 3.2 Rhizosphere and bulk soil microbes associated with carrot storability

#### Differential abundance analysis – indicators of high and low storability

The average bacterial OTU richness was 7.1 in the rhizosphere and 7.0 in the bulk soil of carrots, regardless of whether they exhibited low or high storability. Similarly, Shannon diversity was nearly identical between low- and high-storability carrots, with values of 3479 and 3377 in the rhizosphere, and 3289 and 3266 in bulk soil. According to the differential abundance analysis comparing the microbial communities of carrot samples with high (>80%) and low (≤80%) storability, 356 and 239 bacterial OTUs were respectively abundant in rhizosphere and bulk soil samples associated with high storability (Figure 4). Bacteria that were most abundant with high storability included e.g. bacteria belonging to *Streptomycetaceae* and Solirubrobacterales (Actinomycetota), *Bacillaceae* and *Aerococcaceae* (Bacillota), *Hyphomicrobiaceae* and *TRA3–20* (Pseudomonadota) and *Gemmatimonadaceae* (Gemmatimonadota), Tepidisphaerales WD2101 (Planctomycetota). Moreover, 103 and 228 bacteria were abundant in both rhizosphere and bulk soil samples associated with low storability including e.g., uncultured *Ktedonobacteraceae* (Chloroflexota) and *Micrococcaceae* (Actinomycetota) (Figure 4).

**Figure 4.**
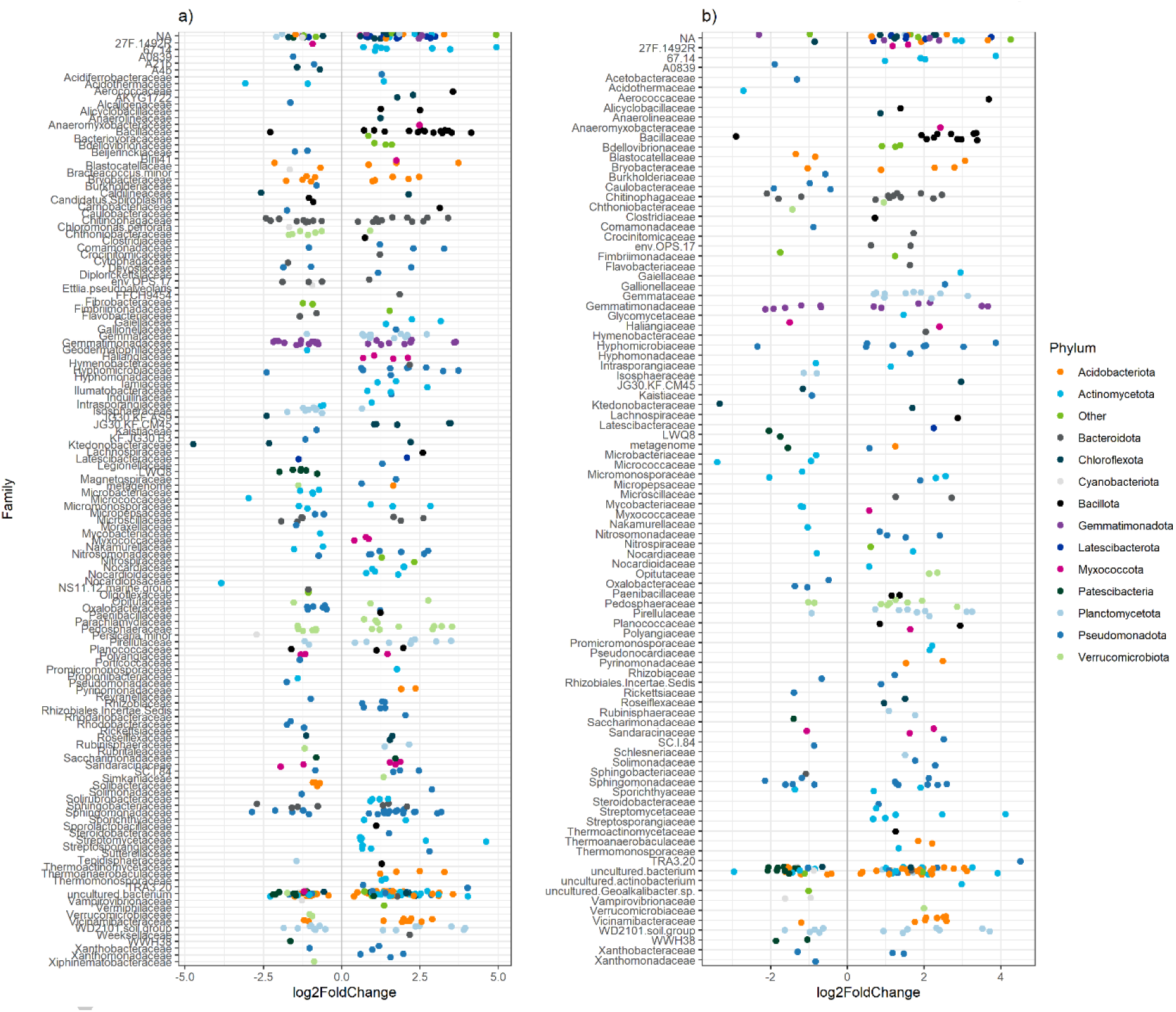
Differential abundance analysis (P<0.05) a) rhizosphere and b) bulk soil indicator bacterial families. Phylum names synonyms Pseudomonadota (syn. Proteobacteria), Actinomycetota (syn. Actinobacteria), Chloroflexota (syn. Chloroflexi) Bacillota (syn. Firmicutes), Cyanobacteriota (syn. Cyanobacteria).

The average fungal OTU richness (511 vs. 450) and Shannon diversity (4.3 vs. 4.0) in the rhizosphere were slightly higher in carrots with high storability compared to those with low storability. In bulk soil, the differences were smaller, with richness values of 499 vs. 475 and Shannon diversity of 4.3 vs. 4.2. Based on the differential abundance analysis, altogether 185 and 121 fungal OTUs were indicative of high storability in rhizosphere and bulk soil. Most indicative (log fold change > 4) of high storability were Ascomycetous *Metarhizium* sp., and unidentified *Pleosporales* and Microascales, Botryotrichum sp., *Chaetomium* sp., and *Tausonia* sp. (Basidiomycota), *Powellomyces* sp. (Chytridiomycota) and *Mortierella* sp. (Mortierellomycota). Furthermore, 38 and 47 fungal OTUs were indicative of carrots with low storability. The differential abundance analysis showed Ascomycetous *Cladorrhinum* sp. to be the most indicative fungi for low storability in both rhizosphere and bulk soil (Figure 5).

**Figure 5.**
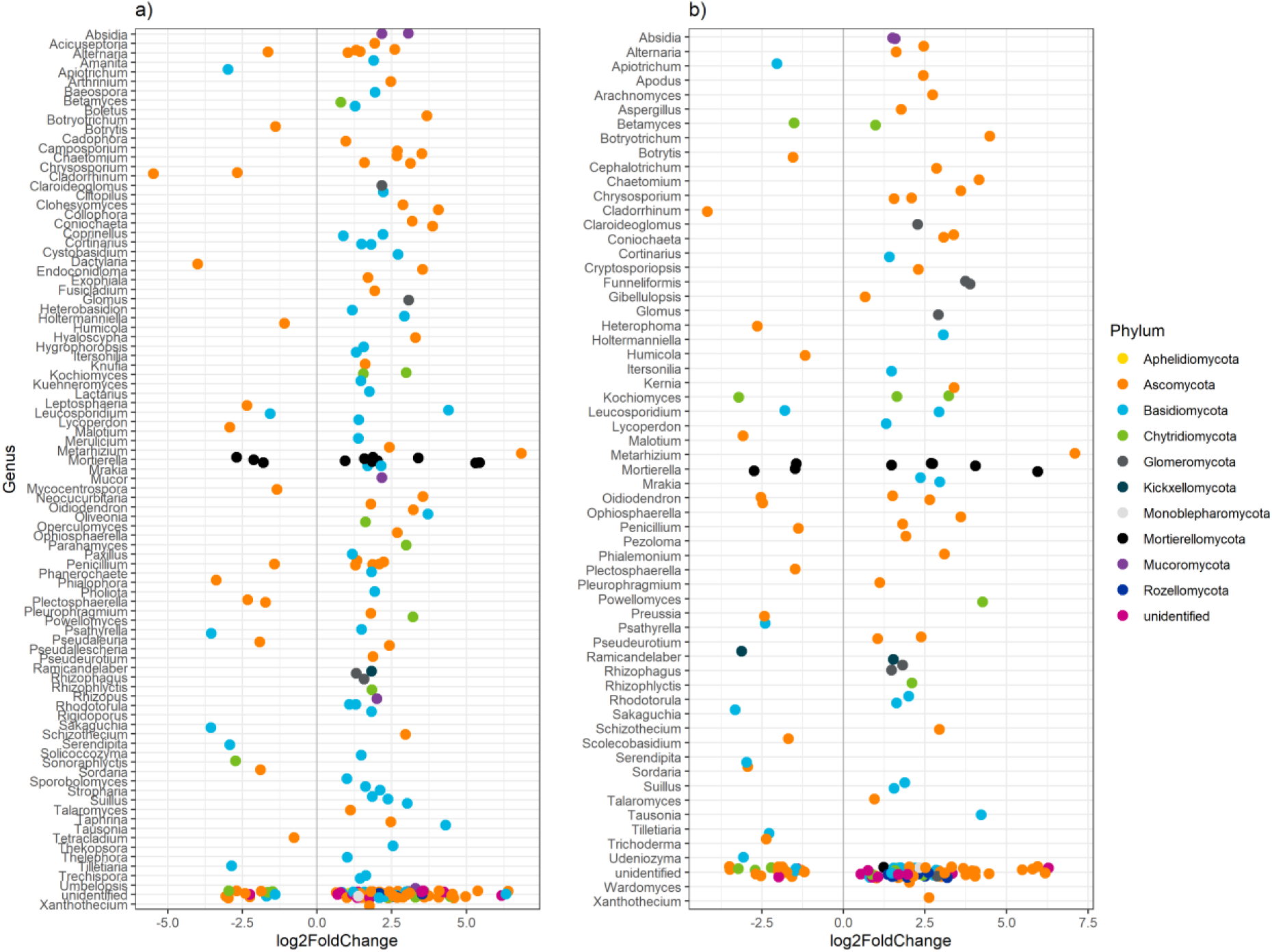
Differential abundance analysis (P<0.05) a) rhizosphere and b) bulk soil indicator fungal genera. Much more fungal OTUs are associated with good storability than with low storability.

#### Microbial Taxa Abundance as a Predictor of Carrot Storability

In the rhizosphere bacteriome, high carrot storability correlated positively (*r* >0.5, *P*<0.001) with abundancy of OTUs belonging to Actinomycetota, Acidobacteriota (*Vicinamibacteraceae*, *Thermoanaerobaculaceae*), Bacillota (*Bacillaceae*), RCP2-54, Planctomycetota (*Gemmataceae*, WD2101), and Verrucomicrobiota (*Pedosphaeraceae*) (Table 3). In the bulk soil bacteriome, high storability correlated positively with abundance of Actinomycetota (unknown 0319-7L14), Planctomycetota (unknown Gemmatales and Planctomycetales, *Rubinisphaeraceae*), Bacillota (*Planococcaceae*) and Acidobacteriota (unknown Vicinamibacterales) (Table 4). Low carrot storability correlated (*r<*-0.5, *P*<0.001) with the presence of Pseudomonadota Burkholderiales agricultural soil bacterium SC-I-84 in both the rhizosphere and bulk soil (Table 3).

**Table 3.**
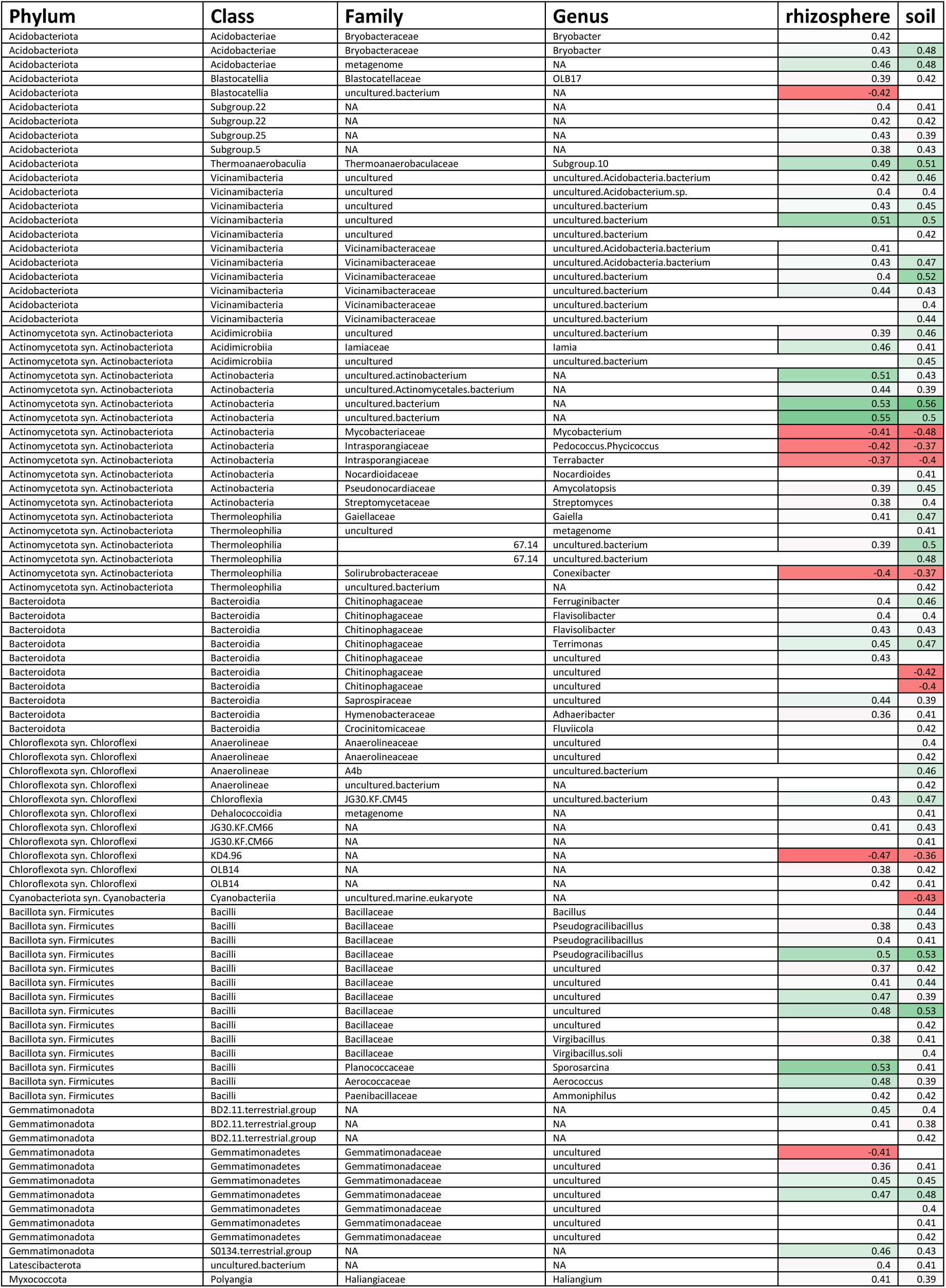
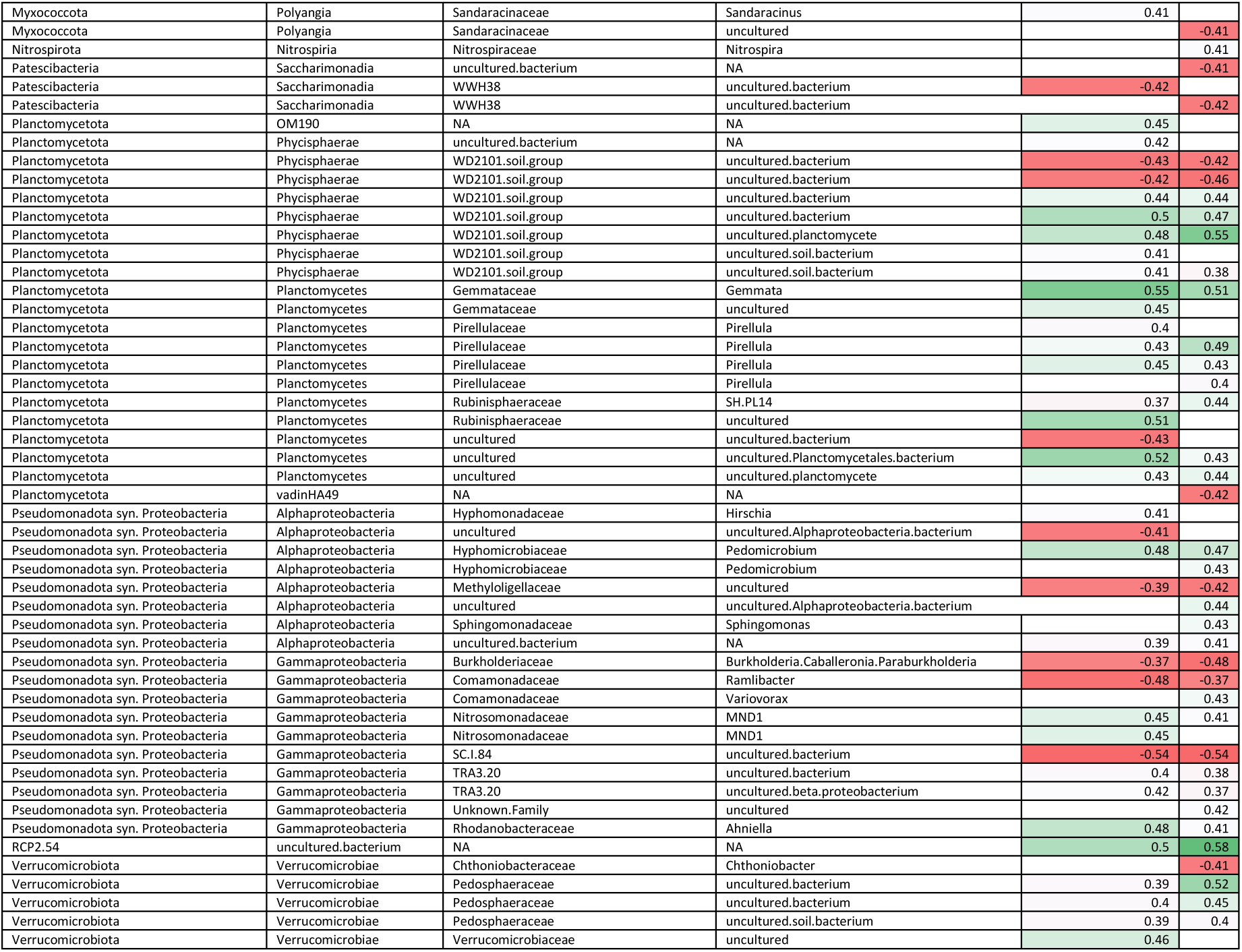
**Correlation analysis between carrot storability and bacterial OTU abundance** based on GMPR-normalized data. Spearman correlation coefficients calculated separately for rhizosphere and bulk soil. Table shows moderate and strong correlations |>0.35|, *P*<0.05.

**Table 4.**
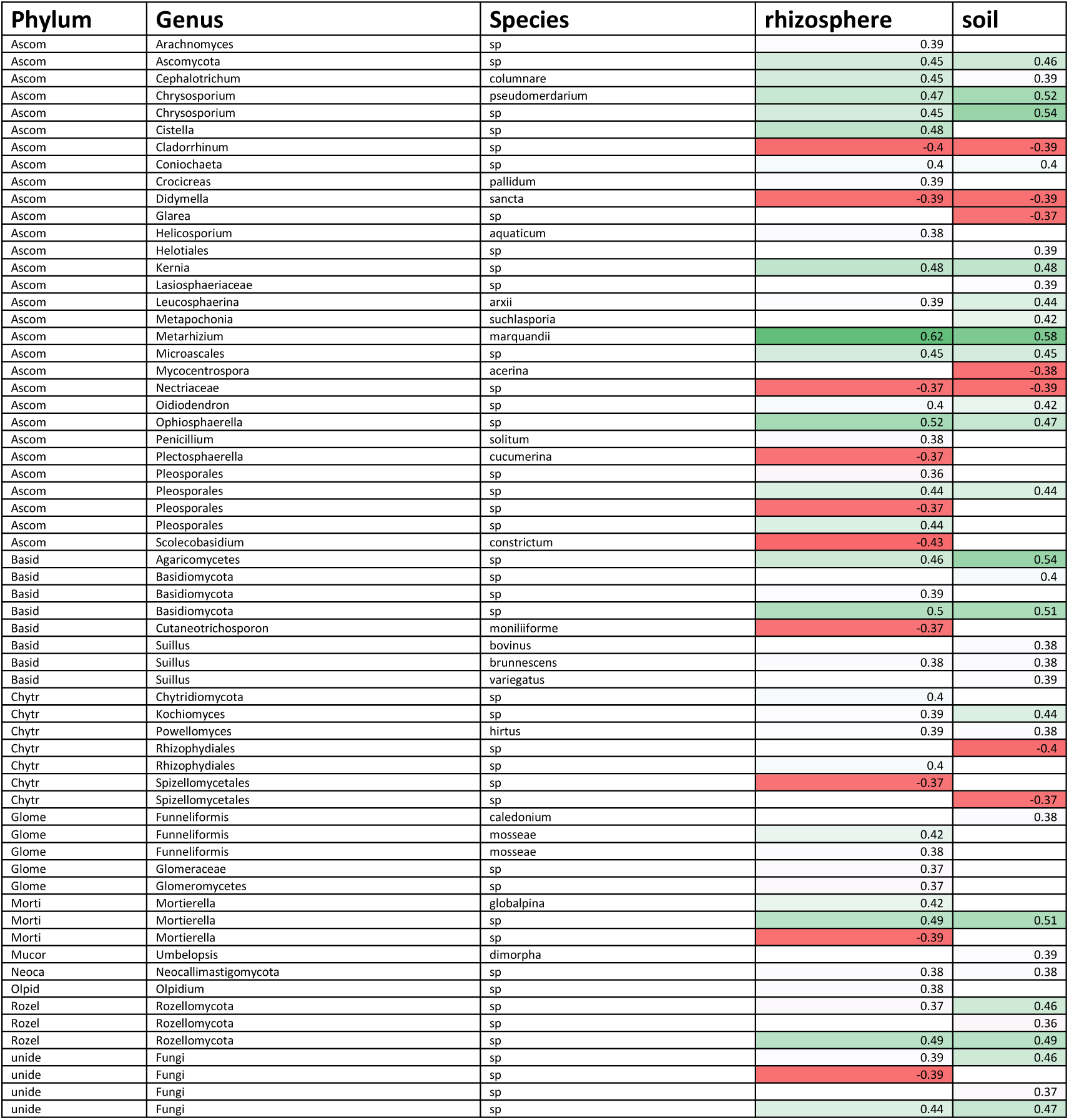
**Correlation analysis between carrot storability and fungal OTU abundance** based on GMPR-normalized data. Spearman correlation coefficients calculated separately for rhizosphere and bulk soil. Table shows moderate and strong correlations |>0.35|, *P*<0.05.

The abundance of ascomycetous fungal OTUs representing Ascomycotous *Chrysosporium* sp., *Metarhizium* sp. and *Ophiosphaerella* sp., Mortierellomycotan *Mortierella* sp., and an unknown Basidiomycota OTU correlated positively (*r* >0.5, *P*<0.001) with high carrot storability, in both rhizosphere and bulk soil (Table 4). Several Rozellomycota, and Glomeromycotan arbuscular mycorrhizal OTUs also had a positive correlation with storability (*r>*0.35, *P*<0.001, Table 4). Low carrot storability was moderately (*r<*-0.35, *P*<0.001) associated with the presence of several Ascomycotous *Mycocentrospora* sp., *Didymella* sp., unidentified *Pleosporales*, *Scolecobasidium* sp., *Glarea* sp., unidentified *Coniochaetales*, *Plectosphaerella* sp., unidentified *Nectriaceae*, and *Cladorrhinum* sp., Basidiomycotan *Cutaneotrichosporon* sp., Chytridiomycotan unidentified *Spizellomycetales*, and Mortierellomycotan *Mortierella* sp. (Table 4).

#### Microbial Taxa Abundance and Soil OM% content

High organic matter content (OM%) of the soil correlated positively (r>0.7, *P* <0.001, Supplementary Table 3) with the abundance of Acidobacteriota (unidentified Acidobacteriae), Actinomycetota (unidentified Acidimicrobiia and Actinobacteria), Bacteroidota Bacteroidia, Chloroflexota KD4-96, Gemmatimonadota Gemmatimonadaceae, Pseudomonadota (Alphaproteobacteria and Gammaproteobacteria) in bulk soil and rhizosphere. Low organic matter was associated with the abundance of Actinomycetota (Actinobacteria, thermoleophilia), Bacillota (Bacilli), Nitrospirota Nitrospiria, Pseudomonadota (unidentified Alphaproteobacteria, Gammaproteobacteria), both in the bulk soil and rhizosphere (Supplementary Table 3).

The abundancy of *Fusicolla* sp., *Mollisia* sp., *Mycosymbioces* sp., *Coniochaeta* sp., *Saitozyma* sp., unidentified *Powellomycetaceae*, and *Umbelopsis* sp. was positively associated with high OM%, whereas the abundancy of *Metarhizium* sp., *Exophala* sp., and unidentified *Mortierellaceae* was negatively associated with the OM% in both rhizosphere and bulk soil (Supplementary Table 4).

## 4 Discussion

### 4.1 Beneficial and pathogenic microbes coexist in carrot rhizosphere and bulk Soil

The results of this study demonstrate that carrot rhizospheres and the surrounding soils contain a complex microbial community comprising both beneficial and potentially pathogenic taxa. Samples were collected from 26 Finnish carrot fields, and the results revealed that neither high nor low storability in carrots was exclusively associated with the presence of pathogens or beneficial microbes. For instance, major carrot pathogens such as *Mycocentrospora* sp. were linked to low storability, yet other fungi commonly considered pathogenic (e.g. *Tilletiaria* sp., *Plectosphaerella* sp.) were detected in fields with both high and low storability carrots. In this study, *Alternaria* sp. were associated with good carrot storability and most *Alternaria* sp. are saprophytes commonly found in soil or on decaying plant tissues (Thomma, 2003). However, *A. dauci*, *A*. *radicina* (not detected in Finland, Latvala et al., 2024) and *A. alternata* are serious pathogens of carrot worldwide. In addition, the presence of beneficial taxa such as *Trichoderma* sp., recognised for their biocontrol potential, was observed in all samples irrespective of storability. These findings further support the hypothesis that the rhizosphere and the surrounding bulk soil harbour a dynamic microbial network wherein both beneficial and pathogenic organisms coexist (Abdelrazek et al., 2020; Kusstatscher et al., 2020). Abdelrazek et al. (2020) reported that carrot taproots host a diverse fungal endophytic community dominated by Ascomycota, largely consistent across management systems. However, they also found that organic management increased richness compared to conventional systems. The reported taxa include pathogens such as *Alternaria dauci*, *Cercospora carotae*, and *Xanthomonas campestris*, along with diverse endophytes reported *Ophiosphaerella*, *Ceratobasidium*, *Colletotrichum*, *Gibberella*, *Cladosporium*, *Aspergillus*, *Cyphellophora*, *Thanatephorus*, *Plectosphaerella*, *Rhizopycnis* and *Phoma* as key members of carrot taproots fungal endophyte community (Abdelrazek et al., 2020).

### 4.2 High microbial diversity might suppress postharvest diseases

In our study postharvest disease incidence could not be predicted by the occurrence of any individual microbial species. Although a high abundance of pathogenic *Mycocentrospora* sp. was correlated with low storability, overall patterns indicated that storability is not solely determined by the presence of carrot pathogens. This finding is consistent with the previous research that indicated indigenous pathogens may only become problematic in conditions of reduced diversity or compositional shifts (Kusstatscher et al., 2020). The data presented herein underscore the notion that storability constitutes a multifactorial trait, influenced by the structure of microbial communities as opposed to individual taxa. This finding aligns with the holobiont concept proposed for postharvest microbiomes (Wisniewski & Droby, 2019).

The high fungal diversity in the rhizosphere was associated with a lower incidence of disease, thus suggesting a possible suppressive effect on postharvest pathogens. Similarly, an association was also observed between bacterial diversity and high storability, particularly when taxa such as *Streptomycetaceae*, *Bacillaceae*, and *Hyphomicrobiaceae* were present in high abundance. These groups are distinguished by their role in nutrient cycling, promotion of plant growth, and antagonism against pathogens through the production of antibiotics (Alipour Kafi et al., 2021; Saxena et al., 2020; Zhang et al., 2024).

However, it was observed that diversity alone was not a universal predictor, as some fields with diverse communities still exhibited disease. This finding suggests that other factors, such as soil physicochemical properties and crop management practices are important for postharvest storability (Batista et al., 2024; Kinkel et al., 2012). As previously reported the rhizosphere microbial composition is shaped by interactions between both biotic and abiotic factors that have substantial, context-dependent effects on rhizosphere microbiomes (Garbeva et al., 2008). Here the bacterial diversity was strongly driven by soil pH, a common phenomenon in acidic soils of boreal field conditions (Fierer & Jackson, 2006; Rousk et al., 2010) and benefitted from organic management.

Fungal diversity was more affected by weather conditions, e.g. rain and temperature, and the use of green manure. This indicates that different management practices are required to support these diverse soil microbial communities.

Organic wood-derived soil amendments are recently reported to enhance beneficial fungal activity in the rhizosphere and promote positive microbial interactions without favouring pathogens (Clocchiatti et al., 2021) or increase fungal biomass in the fields (Rasa et al., 2021). Moreover, high-quality, low C:N-ratio, cover crop residues can improve disease tolerance and microbiome function; major shifts in Proteobacteria and increased abundance of *Bacillaceae* and *Mortierellomycetes* have been reported (Liu et al., 2021). In our study, use of green manure increased the microbial diversity of the soil, but did not per se affect the storability of carrots. Although not statistically significant, the slightly lower storability of carrots from farms using green manure in rotation may reflect multiple factors linked to organic farm management rather than green manure alone; however, our data do not allow further investigation of this.

### 4.3 Good storability is associated with high abundance of beneficial microbial groups

The fields with high-storability carrots demonstrated consistent enrichment of beneficial taxa, including *Devosia* sp., which has been observed to associate with arbuscular mycorrhizal fungi (Zhang et al., 2024), and *Amycolatopsis* sp., which is renowned for producing antibacterial compounds (Kisil et al., 2021). Fungal genera such as *Metarhizium* and *Trichoderma* were linked to enhanced storability, presumably due to their well-known properties in promoting biocontrol and organic matter decomposition (Hermosa et al., 2012; Liao et al., 2014). Conversely, fields with low-storability carrots were dominated by microbes adapted to acidic, low-nutrient soils. While these microbes are not pathogenic, they lack strong disease-suppressive traits. For example, although some members of the Burkholderiaceae family (here associated with low storability) are pathogenic, the majority of them are known as plant growth promoting soil bacteria, and some are even antagonistic to phytopathogenic bacteria and fungi (Coenye & Vandamme, 2003).The results of this study suggest that soil remaining on the carrot surface is important for the storability, and management practices aimed at fostering beneficial microbial groups and enhancing fungal diversity in soil could improve carrot storability (Kalam et al., 2020; Larkin, 2015).

## 5 Conclusion

This study demonstrates that carrot storability is shaped by the composition and diversity of soil and rhizosphere microbiomes rather than the presence of individual pathogens. It is evident that high microbial diversity and the dominance of beneficial taxa are pivotal in the suppression of postharvest diseases and the enhancement of storability. It is recommended that future research endeavours focus on the exploration of functional interactions within these communities and how develop crop management practices to enhance postharvest quality.

## 6 Acknowledgements

This project was funded by the Ministry of Agriculture and Forestry of Finland (VN/4602/2020), and Maiju and Yrjö Rikala’s Horticultural Foundation. We thank researcher Pirjo Kivijärvi for her valuable input in designing and executing this experiment. We are grateful for the technical assistance of Carita Karenius in the laboratory and Johanna Rihtilä in performing the storage experiments and for analyses of storage losses. The authors declare that they have no competing financial interests or personal relationships that could have appeared to influence the work reported in this paper.

## 7 Author contributions

Conceptualization (SV, TP, PK, MH, SL, MP, TSA); Data curation (SV, SL, PK); Formal analysis (SV, TP, PK); Funding acquisition (TP, TSA); Investigation (SV, TP, SL, TSA); Methodology (SV); Project administration (TSA); Resources (TSA); Validation (SV); Visualization (SV); Roles/Writing - original draft (SV, TP, MH, PK); and Writing - review & editing (SL, MP, TSA).

