## Supplementary for "High microbial diversity in the rhizosphere and soil improve carrot (*Daucus carota* L.) postharvest storability"

### Supplementary material – Figures and tables

### Supplementary Figures

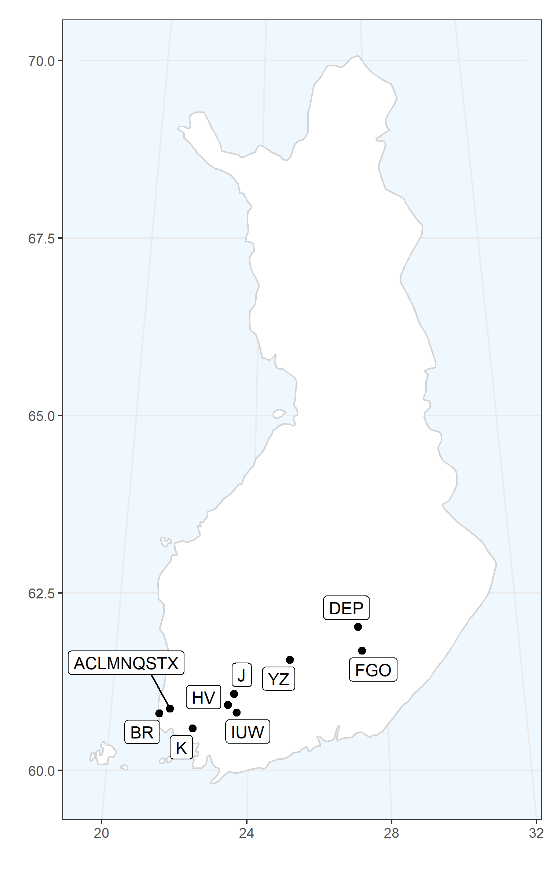

**Supplementary Figure S1** Map of all 26 carrot study sites that were sampled for soil, rhizosphere and carrot samples. The uppercase letters serve as identifier codes for different farm locations (Table 1).

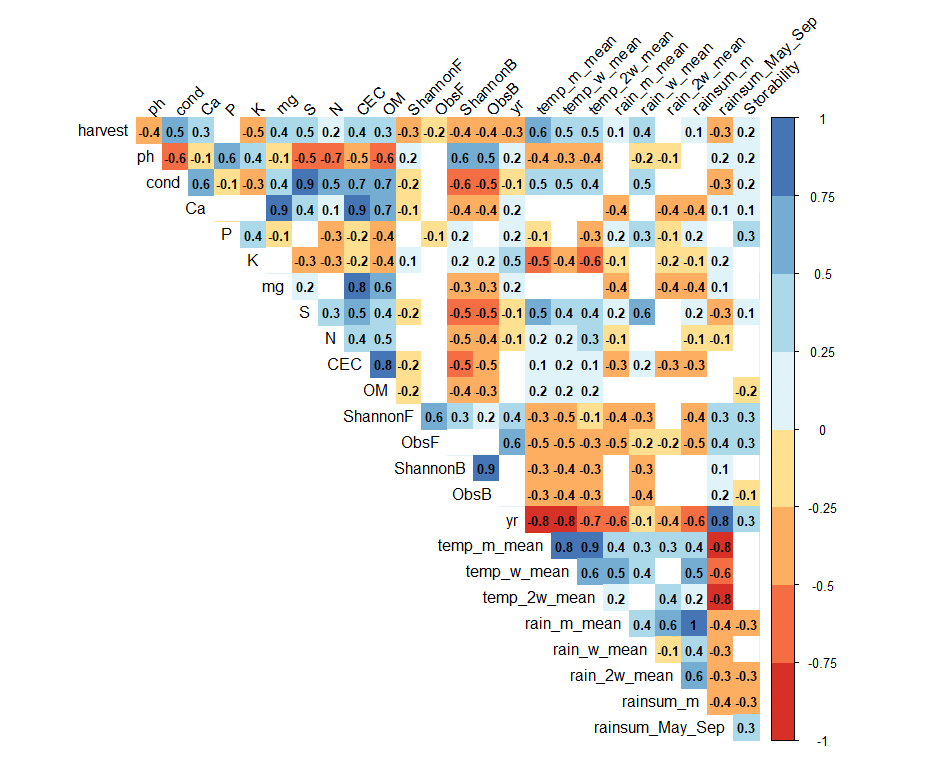

**Supplementary Figure S2** Shows the relationships across soil parameters, and rainfall, temperature, bacterial and fungal diversity, and carrot storability after half year cold storage. Pearson correlation coefficients are presented when *P*<0.05. Carrot storability had weak negative association (*r*= -0.2) with OM%. Soil pH had a strong positive correlation (*r*=0.6) with phosphorus content, and a strong negative correlation with conductivity (*r*=-0.6), nitrogen content (r=-0.7), and OM% (*r*=-0.6). Conductivity had strong positive correlation with soil sulphur (*r*=0.9) and calcium content (*r* =0.6OM% (*r* =0.7,), and moreover, calcium content correlated positively with both OM% (*r* =0.7) and magnesium content (*r* =0.9), and thereby CEC (*r* =0.9) of soil. CEC had a strong positive correlation (*r* =0.8) with soil OM%. Soil phosphorus content had a weak positive correlation (*r* =0.3) with storability. *Monthly mean temperature* of growing season had strong negative correlation (*r* = -0.8) with the total sum of rain during the growing season, and a positive correlation (*r* =0.6) with the total carrot yield. Carrot storability had a weak negative association with mean rainfall (*r* = -0.3). *Preharvest month mean of rainfall* had positive correlation (*r* =0.5) with *preharvest week mean temperature*. *Preharvest week mean of rainfall* correlated positively with soil conductivity and sulphur content (*r*>0.5).

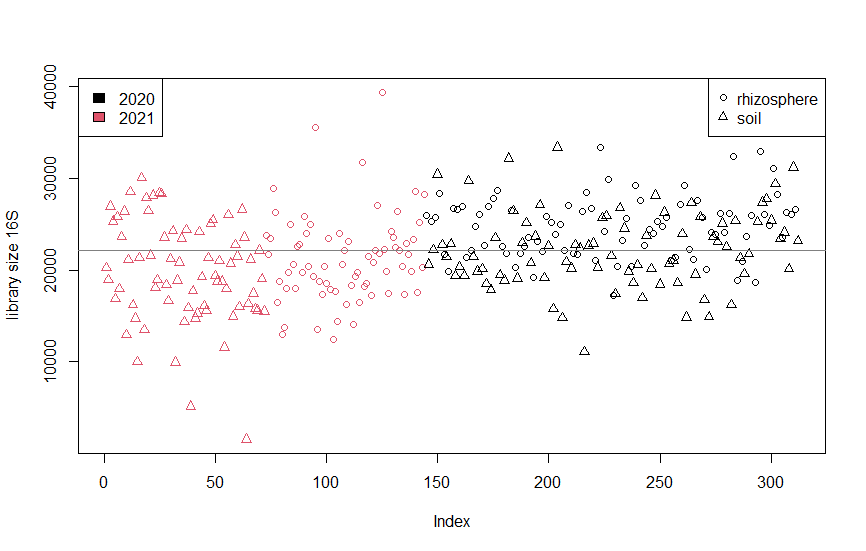

**Supplementary Figure S3** Sequenced 16S library size by sample type and sampling and sequencing year
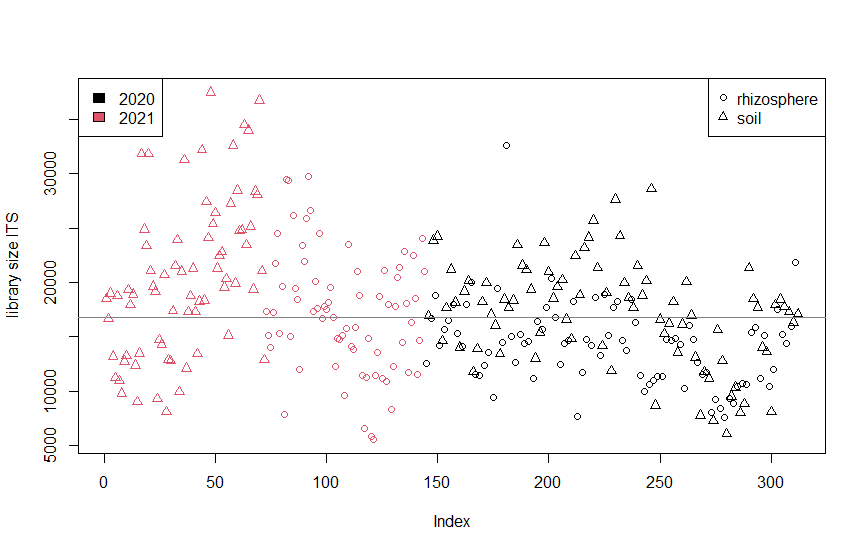

**Supplementary Figure S4** Sequenced ITS library size by sample type and sampling and sequencing year

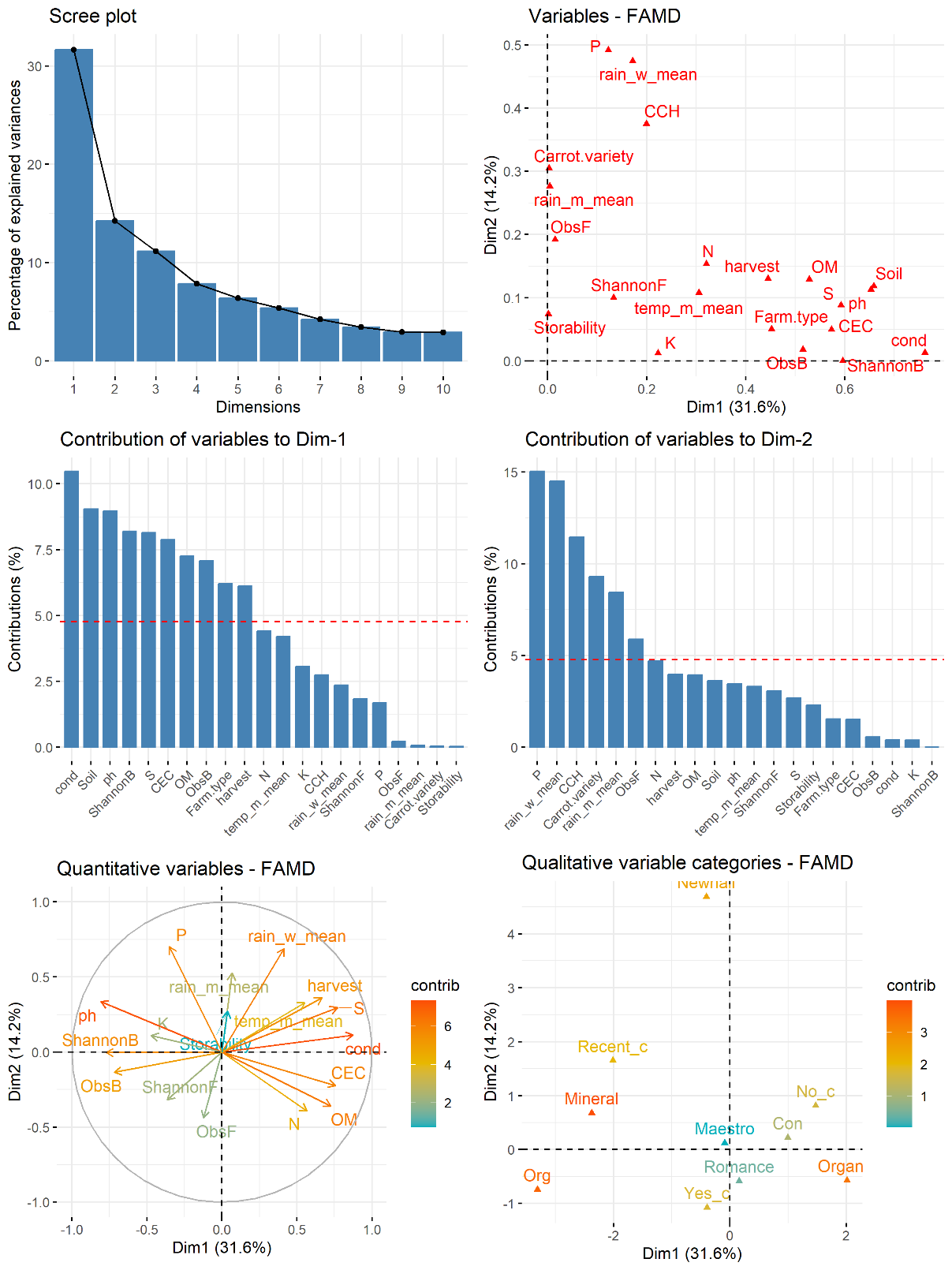

**Supplementary Figure S5 Factor analysis of mixed data** The scree plot (the percentages of inertia explained by each FAMD dimensions, correlation between both quantitative and qualitative variables, the principal dimensions, as well as the contribution of variables to the dimensions 1 and 2. Red dashed line on the graph above indicates the expected average value, if the contributions were uniform. Individuals with similar profiles are close to each other on the factor map. Relationship between variables, the quality of the representation of variables, as well as the correlation between variables and the dimensions, most contributing quantitative variables highlighted.

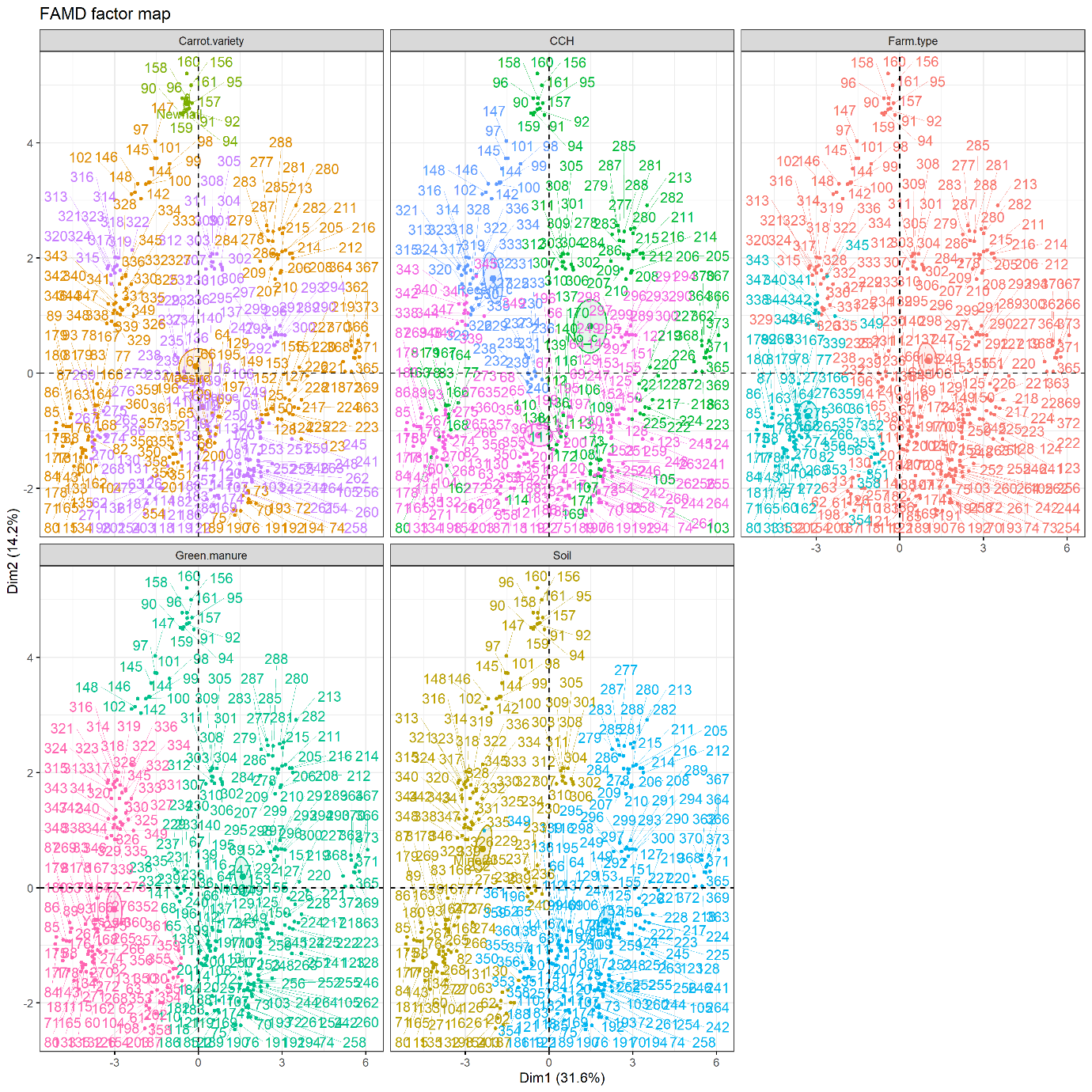

**Supplementary Figure S6 Factor analysis of mixed data** Carrot variety (Newhall on the top, Maestro in the middle, mixed with Romance in the lower half), Carrot in cultivation history (CCH: Recent on the top left, Yes longer time ago on the bottom half, No on the top right), Management (Farm type Conventional on the right Organic on the left), Green manure (Yes on the left, No on the right), and Soil type category (Mineral left, Organic right).

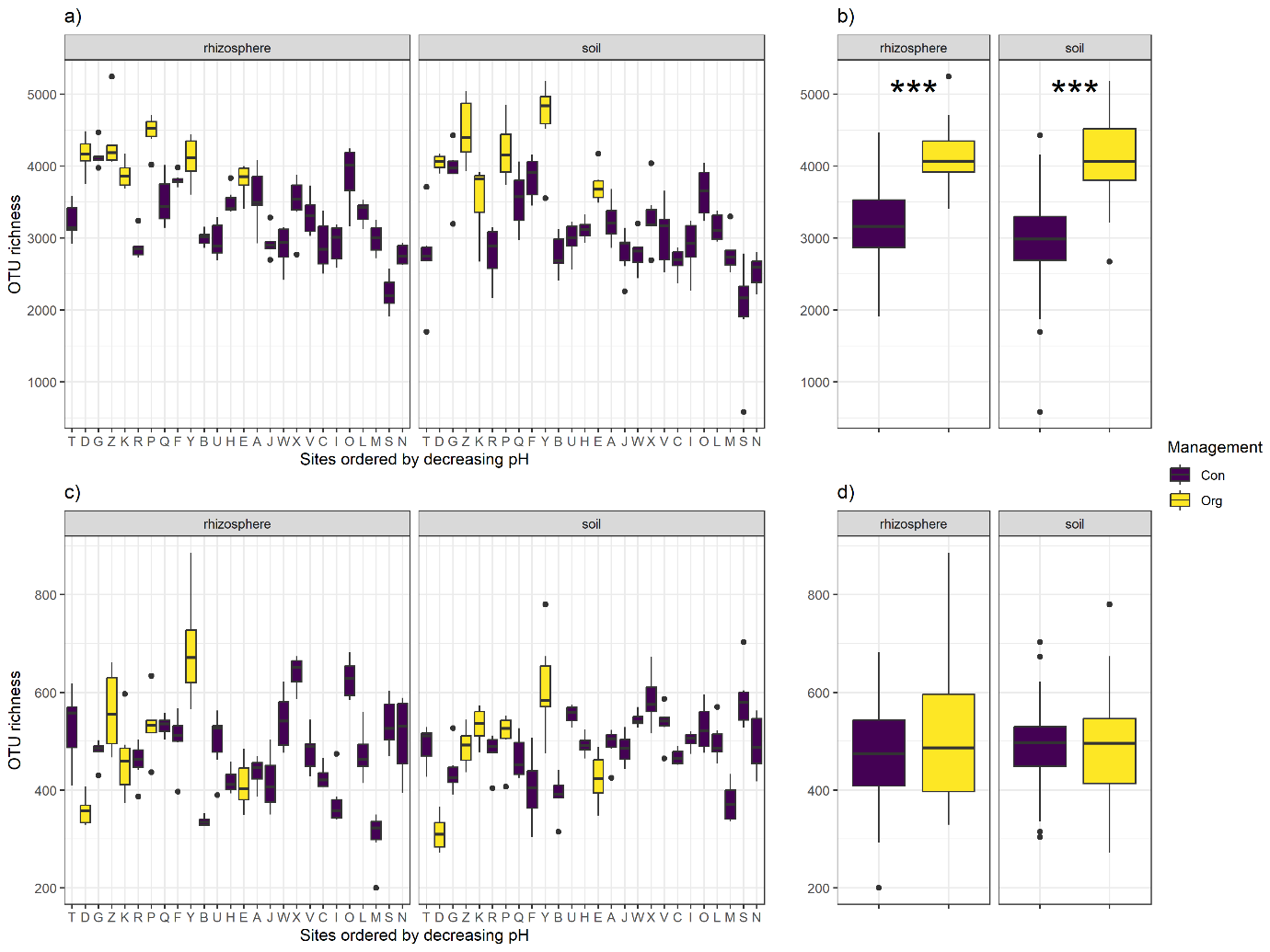

**Supplementary figure S7**. OTU richness of bacterial (a, b) and fungal (c,d) OTUs across different sample types. Boxplots represent the median and interquartile range. (a) Richness grouped by sites, ordered by decreasing soil pH. (b) Richness grouped by management type. *** Statistically significant differences between management types p<0.001.

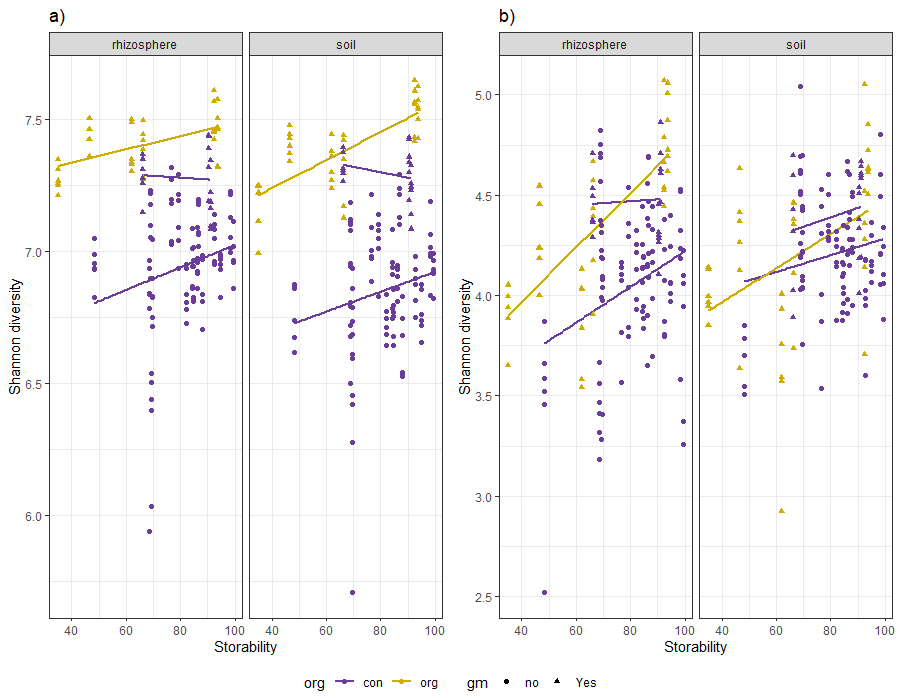

**Supplementary figure S8**. Linear relationships of carrot storability x-axis and Shannon H’ index of a) bacteria and b) fungi. For bacteria only the use of green manure is significant (*P*<0.001). For fungi the Shannon diversity in the rhizosphere increase by use of green manure (*P*<0.01) and is associated to higher storability (*P*<0.05), in bulk soil the is no significant interaction.

### Supplementary Tables

**Supplementary table 1** This table is provided as separate file. Core bacterial and Fungal taxa, 90% prevalence, associated with rhizosphere and bilk soil, and storability.

**Supplementary table 2** Table shows envfit (vegan package, R) results fitting environmental variables onto ordination axes. NMDS1 and NMDS2 are vector coordinates on the first two NMDS axes. r2 is the squared correlation coefficient indicating fit strength. Pr(>r) is the p-value from permutation tests assessing significance. Significant variables (p < 0.05) strongly correlate with the ordination gradients.

|  | Bacteria | NMDS1 | NMDS2 | r2 | Pr(>r) | Fungi | NMDS1 | NMDS2 | r2 | Pr(>r) |
| --- | --- | --- | --- | --- | --- | --- | --- | --- | --- | --- |
| ph |  | -0.99 | 0.17 | 0.78 | 0.001 |  | -0.91 | 0.43 | 0.7 | 0.001 |
| cond |  | 0.81 | 0.58 | 0.3 | 0.001 |  | 0.98 | 0.21 | 0.49 | 0.001 |
| P |  | -0.72 | 0.69 | 0.46 | 0.001 |  | -0.59 | 0.81 | 0.42 | 0.001 |
| K |  | -0.59 | 0.81 | 0.2 | 0.001 |  | -0.86 | 0.51 | 0.2 | 0.001 |
| S |  | 0.62 | 0.78 | 0.21 | 0.001 |  | 0.87 | 0.5 | 0.4 | 0.001 |
| N |  | 0.94 | -0.35 | 0.42 | 0.001 |  | 0.74 | -0.68 | 0.40 | 0.001 |
| OM |  | 0.9 | -0.43 | 0.42 | 0.001 |  | 0.88 | -0.47 | 0.54 | 0.001 |
| CEC |  | 0.84 | 0.54 | 0.33 | 0.001 |  | 1 | 0.04 | 0.5 | 0.001 |
| Storability |  | -0.43 | 0.9 | 0.09 | 0.001 |  | -0.02 | 1 | 0.03 | 0.007 |
| Rain |  | 0.17 | 0.99 | 0.17 | 0.001 |  | 0.42 | 0.91 | 0.25 | 0.001 |
| GM, no |  | 0.18 | 0.1 | 0.31 | 0.001 |  | 0.24 | 0.07 |  | 0.001 |
| GM, yes |  | -0.33 | -0.19 | 0.31 | 0.001 |  | -0.45 | -0.14 | 0.46 | 0.001 |

**Supplementary table 3** Correlation analysis between soil OM% and bacterial OTU abundance based on GMPR-normalized data. Spearman correlation coefficients calculated separately for rhizosphere and bulk soil. Table shows moderate and strong correlations |>0.5|, P<0.05.

| phylum | order | family | genus | soil | rhizo |
| --- | --- | --- | --- | --- | --- |
| Abditibacteriota | Abditibacteriales | Abditibacteriaceae | Abditibacterium |  | -0.53 |
| Acidobacteriota | Acidobacteriales | Acidobacteriaceae..Subgroup.1. | Acidipila.Silvibacterium | 0.54 |  |
|  | Acidobacteriales | Acidobacteriaceae..Subgroup.1. | Acidipila.Silvibacterium | 0.56 | 0.54 |
|  | Acidobacteriales | Acidobacteriaceae..Subgroup.1. | Edaphobacter | 0.60 | 0.52 |
|  | Acidobacteriales | Acidobacteriaceae..Subgroup.1. | Granulicella | 0.59 | 0.52 |
|  | Acidobacteriales | Acidobacteriaceae..Subgroup.1. | Granulicella | 0.54 | 0.53 |
|  | Acidobacteriales | Acidobacteriaceae..Subgroup.1. | Occallatibacter | 0.62 | 0.56 |
|  | Acidobacteriales | Acidobacteriaceae..Subgroup.1. | Terracidiphilus | 0.60 | 0.57 |
|  | Acidobacteriales | Acidobacteriaceae..Subgroup.1. | uncultured | 0.59 | 0.57 |
|  | Acidobacteriales | Acidobacteriaceae..Subgroup.1. | uncultured | 0.53 | 0.52 |
|  | Acidobacteriales | Koribacteraceae | Candidatus.Koribacter | 0.73 | 0.68 |
|  | Acidobacteriales | Koribacteraceae | Candidatus.Koribacter |  | 0.57 |
|  | Acidobacteriales | Koribacteraceae | Candidatus.Koribacter | 0.56 | 0.57 |
|  | Acidobacteriales | Koribacteraceae | Candidatus.Koribacter | 0.52 | 0.51 |
|  | Acidobacteriales | uncultured | uncultured.Acidobacteria.bacterium | 0.70 | 0.71 |
|  | Acidobacteriales | uncultured | uncultured.Acidobacteria.bacterium | 0.60 | 0.54 |
|  | Acidobacteriales | uncultured | uncultured.Acidobacteria.bacterium | 0.70 | 0.72 |
|  | Acidobacteriales | uncultured | uncultured.Acidobacteria.bacterium | 0.60 | 0.55 |
|  | Acidobacteriales | uncultured | uncultured.Acidobacteria.bacterium | 0.60 | 0.59 |
|  | Acidobacteriales | uncultured | uncultured.Acidobacteria.bacterium | 0.58 |  |
|  | Acidobacteriales | uncultured | uncultured.Acidobacteria.bacterium | 0.64 | 0.61 |
|  | Acidobacteriales | uncultured | uncultured.Acidobacteria.bacterium | 0.60 | 0.58 |
|  | Acidobacteriales | uncultured | uncultured.bacterium | 0.59 | 0.61 |
|  | Acidobacteriales | uncultured | uncultured.bacterium | 0.64 | 0.65 |
|  | Acidobacteriales | uncultured | uncultured.bacterium | 0.66 | 0.66 |
|  | Acidobacteriales | uncultured | uncultured.bacterium | 0.61 | 0.62 |
|  | Acidobacteriales | uncultured | uncultured.bacterium | 0.72 | 0.73 |
|  | Acidobacteriales | uncultured | uncultured.bacterium | 0.60 | 0.60 |
|  | Acidobacteriales | uncultured | uncultured.bacterium | 0.69 | 0.67 |
|  | Acidobacteriales | uncultured | uncultured.bacterium | 0.67 | 0.69 |
|  | Acidobacteriales | uncultured | uncultured.bacterium | 0.50 |  |
|  | Acidobacteriales | uncultured | uncultured.bacterium | 0.55 | 0.60 |
|  | Acidobacteriales | uncultured | uncultured.bacterium | 0.55 | 0.56 |
|  | Acidobacteriales | uncultured | uncultured.bacterium | 0.53 |  |
|  | Acidobacteriales | uncultured | uncultured.bacterium | 0.64 | 0.60 |
|  | Acidobacteriales | uncultured | uncultured.bacterium | 0.62 | 0.55 |
|  | Acidobacteriales | uncultured | uncultured.bacterium | 0.56 | 0.60 |
|  | Acidobacteriales | uncultured | uncultured.bacterium | 0.62 | 0.59 |
|  | Acidobacteriales | uncultured | uncultured.bacterium | -0.59 | -0.55 |
|  | Acidobacteriales | uncultured | uncultured.bacterium | 0.58 |  |
|  | Bryobacterales | Bryobacteraceae | Bryobacter | 0.56 | 0.55 |
|  | Bryobacterales | Bryobacteraceae | Bryobacter | -0.57 |  |
|  | Bryobacterales | Bryobacteraceae | Bryobacter | 0.65 | 0.60 |
|  | Bryobacterales | Bryobacteraceae | Bryobacter | 0.61 | 0.61 |
|  | Bryobacterales | Bryobacteraceae | Bryobacter | 0.57 | 0.54 |
|  | Bryobacterales | Bryobacteraceae | Bryobacter | 0.72 | 0.69 |
|  | Bryobacterales | Bryobacteraceae | Bryobacter | 0.64 | 0.66 |
|  | Bryobacterales | Bryobacteraceae | Bryobacter | 0.60 | 0.65 |
|  | Bryobacterales | Bryobacteraceae | Bryobacter | 0.66 | 0.66 |
|  | Bryobacterales | Bryobacteraceae | Bryobacter | 0.66 | 0.63 |
|  | Bryobacterales | Bryobacteraceae | Bryobacter | 0.62 | 0.60 |
|  | Bryobacterales | Bryobacteraceae | Bryobacter | -0.67 | -0.56 |
|  | Bryobacterales | Bryobacteraceae | Bryobacter | 0.53 | 0.55 |
|  | Bryobacterales | Bryobacteraceae | Bryobacter |  | 0.56 |
|  | Bryobacterales | Bryobacteraceae | Bryobacter | 0.56 | 0.57 |
|  | Bryobacterales | Bryobacteraceae | Bryobacter | -0.59 | -0.53 |
|  | Bryobacterales | Bryobacteraceae | Bryobacter | -0.53 | -0.63 |
|  | Bryobacterales | Bryobacteraceae | Bryobacter | 0.59 |  |
|  | Bryobacterales | Bryobacteraceae | Bryobacter | 0.50 |  |
|  | Bryobacterales | Bryobacteraceae | Bryobacter |  | 0.51 |
|  | Bryobacterales | Bryobacteraceae | Bryobacter | -0.59 | -0.59 |
|  | Bryobacterales | Bryobacteraceae | Bryobacter |  | -0.53 |
|  | PAUC26f | uncultured.bacterium | NA | -0.50 |  |
|  | Solibacterales | Solibacteraceae | Candidatus.Solibacter | 0.58 | 0.61 |
|  | Solibacterales | Solibacteraceae | Candidatus.Solibacter | 0.67 | 0.66 |
|  | Solibacterales | Solibacteraceae | Candidatus.Solibacter | 0.57 | 0.55 |
|  | Solibacterales | Solibacteraceae | Candidatus.Solibacter | 0.70 | 0.69 |
|  | Solibacterales | Solibacteraceae | Candidatus.Solibacter |  | 0.54 |
|  | Solibacterales | Solibacteraceae | Candidatus.Solibacter | 0.62 | 0.58 |
|  | Solibacterales | Solibacteraceae | Candidatus.Solibacter | 0.75 | 0.76 |
|  | Solibacterales | Solibacteraceae | Candidatus.Solibacter | 0.61 | 0.55 |
|  | Solibacterales | Solibacteraceae | Candidatus.Solibacter |  | 0.53 |
|  | Solibacterales | Solibacteraceae | Candidatus.Solibacter | 0.63 | 0.63 |
|  | Solibacterales | Solibacteraceae | Candidatus.Solibacter | 0.62 | 0.53 |
|  | Solibacterales | Solibacteraceae | Candidatus.Solibacter | 0.66 | 0.60 |
|  | Solibacterales | Solibacteraceae | Candidatus.Solibacter | 0.57 | 0.54 |
|  | Solibacterales | Solibacteraceae | Candidatus.Solibacter | 0.57 | 0.53 |
|  | Solibacterales | Solibacteraceae | Candidatus.Solibacter | 0.56 | 0.65 |
|  | Solibacterales | Solibacteraceae | Candidatus.Solibacter |  | -0.51 |
|  | Solibacterales | Solibacteraceae | Candidatus.Solibacter | 0.51 |  |
|  | Solibacterales | Solibacteraceae | Candidatus.Solibacter | -0.57 | -0.55 |
|  | Solibacterales | Solibacteraceae | Candidatus.Solibacter | 0.56 |  |
|  | Solibacterales | Solibacteraceae | Candidatus.Solibacter | 0.52 |  |
|  | Subgroup.13 | uncultured.bacterium | NA |  | 0.51 |
|  | Subgroup.13 | uncultured.Holophaga.sp. | NA | 0.53 |  |
|  | Subgroup.2 | uncultured.Acidobacteria.bacterium | NA | 0.63 | 0.61 |
|  | Subgroup.2 | uncultured.Acidobacteria.bacterium | NA | 0.81 | 0.79 |
|  | Subgroup.2 | uncultured.Acidobacteria.bacterium | NA | 0.57 | 0.56 |
|  | Subgroup.2 | uncultured.bacterium | NA | 0.68 | 0.73 |
|  | Subgroup.2 | uncultured.bacterium | NA | 0.70 | 0.72 |
|  | Subgroup.2 | uncultured.bacterium | NA | 0.74 | 0.78 |
|  | Subgroup.2 | uncultured.bacterium | NA | 0.66 | 0.69 |
|  | Subgroup.2 | uncultured.bacterium | NA | 0.66 | 0.68 |
|  | Subgroup.2 | uncultured.bacterium | NA | 0.64 | 0.66 |
|  | Subgroup.2 | uncultured.bacterium | NA | 0.65 | 0.66 |
|  | Subgroup.2 | uncultured.bacterium | NA | 0.64 | 0.62 |
|  | Subgroup.2 | uncultured.bacterium | NA | 0.56 | 0.55 |
|  | Subgroup.2 | uncultured.bacterium | NA | 0.57 |  |
|  | Subgroup.2 | uncultured.bacterium | NA | 0.55 | 0.54 |
|  | Subgroup.2 | uncultured.bacterium | NA | 0.54 | 0.56 |
|  | Subgroup.2 | uncultured.bacterium | NA | 0.70 | 0.67 |
|  | Subgroup.2 | uncultured.bacterium | NA | 0.57 | 0.50 |
|  | uncultured.Acidobacteriaceae.bacterium | NA | NA | -0.54 | -0.57 |
|  | 45597 | uncultured.Acidobacteria.bacterium | NA | -0.63 | -0.56 |
|  | 45597 | uncultured.Acidobacterium.sp. | NA | -0.52 |  |
|  | 45597 | uncultured.bacterium | NA | -0.58 | -0.50 |
|  | 45597 | uncultured.bacterium | NA | -0.53 |  |
|  | 45597 | uncultured.bacterium | NA |  | -0.50 |
|  | Blastocatellales | Blastocatellaceae | JGI.0001001.H03 | -0.50 | -0.54 |
|  | Blastocatellales | Blastocatellaceae | JGI.0001001.H03 | -0.52 |  |
|  | Blastocatellales | Blastocatellaceae | JGI.0001001.H03 | 0.62 | 0.62 |
|  | Blastocatellales | Blastocatellaceae | uncultured | -0.54 |  |
|  | Blastocatellales | Blastocatellaceae | uncultured |  | -0.54 |
|  | DS.100 | uncultured.bacterium | NA |  | -0.51 |
|  | DS.100 | uncultured.bacterium | NA | -0.54 |  |
|  | Pyrinomonadales | Pyrinomonadaceae | RB41 |  | -0.52 |
|  | Pyrinomonadales | Pyrinomonadaceae | RB41 | -0.58 | -0.54 |
|  | Pyrinomonadales | Pyrinomonadaceae | RB41 | -0.64 | -0.64 |
|  | Pyrinomonadales | Pyrinomonadaceae | RB41 | -0.67 | -0.64 |
|  | Pyrinomonadales | Pyrinomonadaceae | RB41 |  | 0.50 |
|  | Pyrinomonadales | Pyrinomonadaceae | RB41 | -0.52 | -0.64 |
|  | Pyrinomonadales | Pyrinomonadaceae | RB41 | -0.59 | -0.56 |
|  | Pyrinomonadales | Pyrinomonadaceae | RB41 | -0.52 | -0.54 |
|  | Pyrinomonadales | Pyrinomonadaceae | RB41 | -0.51 |  |
|  | Subgroup.7 | metagenome | NA | -0.51 |  |
|  | Subgroup.7 | uncultured.bacterium | NA | 0.67 | 0.65 |
|  | Subgroup.7 | uncultured.bacterium | NA | -0.65 | -0.59 |
|  | Subgroup.7 | uncultured.bacterium | NA |  | -0.53 |
|  | Subgroup.7 | uncultured.delta.proteobacterium | NA | -0.56 | -0.55 |
|  | uncultured.bacterium | NA | NA | -0.64 | -0.62 |
|  | uncultured.bacterium | NA | NA | -0.58 |  |
|  | uncultured.bacterium | NA | NA | -0.54 |  |
|  | uncultured.soil.bacterium | NA | NA | -0.52 |  |
|  | metagenome | NA | NA | 0.51 |  |
|  | uncultured.Acidobacteria.bacterium | NA | NA | -0.54 | -0.53 |
|  | uncultured.Acidobacteria.bacterium | NA | NA | -0.50 |  |
|  | uncultured.Acidobacteriales.bacterium | NA | NA | -0.55 |  |
|  | uncultured.bacterium | NA | NA |  | -0.58 |
|  | Thermoanaerobaculales | Thermoanaerobaculaceae | Subgroup.10 | -0.55 |  |
|  | Thermoanaerobaculales | Thermoanaerobaculaceae | Subgroup.10 | -0.60 |  |
|  | Thermoanaerobaculales | Thermoanaerobaculaceae | Subgroup.10 | -0.57 | -0.52 |
|  | Thermoanaerobaculales | Thermoanaerobaculaceae | Subgroup.10 | -0.51 |  |
|  | Subgroup.17 | uncultured.Acidobacteria.bacterium | NA | -0.60 | -0.57 |
|  | Subgroup.17 | uncultured.Acidobacteria.bacterium | NA | -0.60 | -0.53 |
|  | Subgroup.17 | uncultured.Acidobacteriales.bacterium | NA | -0.62 | -0.57 |
|  | Subgroup.17 | uncultured.Acidobacteriales.bacterium | NA | -0.56 |  |
|  | Subgroup.17 | uncultured.bacterium | NA | -0.56 | -0.50 |
|  | Subgroup.17 | uncultured.bacterium | NA | -0.54 |  |
|  | Subgroup.17 | uncultured.bacterium | NA | -0.53 |  |
|  | Vicinamibacterales | uncultured | metagenome | -0.54 |  |
|  | Vicinamibacterales | uncultured | uncultured.Acidobacteria.bacterium | 0.61 | 0.58 |
|  | Vicinamibacterales | uncultured | uncultured.Acidobacteria.bacterium | -0.55 | -0.53 |
|  | Vicinamibacterales | uncultured | uncultured.Acidobacteria.bacterium | -0.55 | -0.52 |
|  | Vicinamibacterales | uncultured | uncultured.Acidobacteria.bacterium | -0.51 |  |
|  | Vicinamibacterales | uncultured | uncultured.Acidobacteria.bacterium | -0.60 | -0.55 |
|  | Vicinamibacterales | uncultured | uncultured.Acidobacteria.bacterium | 0.68 | 0.62 |
|  | Vicinamibacterales | uncultured | uncultured.Acidobacteria.bacterium | -0.55 |  |
|  | Vicinamibacterales | uncultured | uncultured.Acidobacteria.bacterium | -0.60 | -0.60 |
|  | Vicinamibacterales | uncultured | uncultured.Acidobacteria.bacterium | -0.56 |  |
|  | Vicinamibacterales | uncultured | uncultured.Acidobacteria.bacterium | -0.60 | -0.56 |
|  | Vicinamibacterales | uncultured | uncultured.Acidobacteria.bacterium | -0.57 | -0.53 |
|  | Vicinamibacterales | uncultured | uncultured.Acidobacteria.bacterium | -0.56 | -0.53 |
|  | Vicinamibacterales | uncultured | uncultured.Acidobacteria.bacterium | -0.59 | -0.51 |
|  | Vicinamibacterales | uncultured | uncultured.Acidobacteriaceae.bacterium | -0.51 |  |
|  | Vicinamibacterales | uncultured | uncultured.Acidobacteriales.bacterium | 0.71 | 0.58 |
|  | Vicinamibacterales | uncultured | uncultured.Acidobacteriales.bacterium | -0.55 |  |
|  | Vicinamibacterales | uncultured | uncultured.bacterium | 0.70 | 0.71 |
|  | Vicinamibacterales | uncultured | uncultured.bacterium | -0.56 |  |
|  | Vicinamibacterales | uncultured | uncultured.bacterium | -0.59 | -0.54 |
|  | Vicinamibacterales | uncultured | uncultured.bacterium | -0.64 | -0.64 |
|  | Vicinamibacterales | uncultured | uncultured.bacterium | -0.55 |  |
|  | Vicinamibacterales | uncultured | uncultured.bacterium | 0.52 |  |
|  | Vicinamibacterales | uncultured | uncultured.bacterium | 0.71 | 0.72 |
|  | Vicinamibacterales | uncultured | uncultured.bacterium | 0.68 | 0.66 |
|  | Vicinamibacterales | uncultured | uncultured.bacterium | -0.73 | -0.71 |
|  | Vicinamibacterales | uncultured | uncultured.bacterium | -0.69 | -0.65 |
|  | Vicinamibacterales | uncultured | uncultured.bacterium | -0.70 | -0.68 |
|  | Vicinamibacterales | uncultured | uncultured.bacterium | -0.56 | -0.56 |
|  | Vicinamibacterales | uncultured | uncultured.bacterium | -0.56 | -0.51 |
|  | Vicinamibacterales | uncultured | uncultured.bacterium |  | -0.51 |
|  | Vicinamibacterales | uncultured | uncultured.bacterium | 0.51 |  |
|  | Vicinamibacterales | uncultured | uncultured.bacterium | -0.51 |  |
|  | Vicinamibacterales | uncultured | uncultured.bacterium.gp6 | -0.59 | -0.63 |
|  | Vicinamibacterales | uncultured | uncultured.soil.bacterium |  | -0.51 |
|  | Vicinamibacterales | uncultured.bacterium | NA | -0.50 |  |
|  | Vicinamibacterales | Vicinamibacteraceae | Luteitalea | -0.61 |  |
|  | Vicinamibacterales | Vicinamibacteraceae | metagenome | -0.69 | -0.66 |
|  | Vicinamibacterales | Vicinamibacteraceae | metagenome | -0.54 | -0.61 |
|  | Vicinamibacterales | Vicinamibacteraceae | uncultured | -0.67 | -0.68 |
|  | Vicinamibacterales | Vicinamibacteraceae | uncultured |  | -0.52 |
|  | Vicinamibacterales | Vicinamibacteraceae | uncultured | -0.51 | -0.55 |
|  | Vicinamibacterales | Vicinamibacteraceae | uncultured.Acidobacteria.bacterium | -0.65 | -0.59 |
|  | Vicinamibacterales | Vicinamibacteraceae | uncultured.Acidobacteria.bacterium | -0.65 | -0.57 |
|  | Vicinamibacterales | Vicinamibacteraceae | uncultured.Acidobacteria.bacterium |  | -0.52 |
|  | Vicinamibacterales | Vicinamibacteraceae | uncultured.Acidobacteria.bacterium | -0.51 |  |
|  | Vicinamibacterales | Vicinamibacteraceae | uncultured.Acidobacteria.bacterium | -0.54 | -0.58 |
|  | Vicinamibacterales | Vicinamibacteraceae | uncultured.Acidobacteria.bacterium | -0.52 | -0.57 |
|  | Vicinamibacterales | Vicinamibacteraceae | uncultured.bacterium | -0.57 | -0.60 |
|  | Vicinamibacterales | Vicinamibacteraceae | uncultured.bacterium | -0.62 | -0.56 |
|  | Vicinamibacterales | Vicinamibacteraceae | uncultured.bacterium | -0.62 | -0.60 |
|  | Vicinamibacterales | Vicinamibacteraceae | uncultured.bacterium | -0.60 | -0.55 |
|  | Vicinamibacterales | Vicinamibacteraceae | uncultured.bacterium | -0.69 | -0.63 |
|  | Vicinamibacterales | Vicinamibacteraceae | uncultured.bacterium | -0.72 | -0.65 |
|  | Vicinamibacterales | Vicinamibacteraceae | uncultured.bacterium | -0.54 | -0.54 |
|  | Vicinamibacterales | Vicinamibacteraceae | uncultured.bacterium | -0.63 | -0.54 |
|  | Vicinamibacterales | Vicinamibacteraceae | uncultured.bacterium | -0.50 |  |
|  | Vicinamibacterales | Vicinamibacteraceae | Vicinamibacter | -0.60 | -0.62 |
|  | Vicinamibacterales | Vicinamibacteraceae | Vicinamibacter | -0.53 |  |
| Actinomycetota syn. Actinobacteria | Actinomarinales | uncultured | uncultured.bacterium | -0.66 | -0.62 |
|  | IMCC26256 | bacterium.enrichment.culture.clone.auto73.4W | NA | 0.52 | 0.51 |
|  | IMCC26256 | uncultured.Acidimicrobiales.bacterium | NA | 0.58 | 0.62 |
|  | IMCC26256 | uncultured.Aciditerrimonas.sp. | NA | 0.62 | 0.50 |
|  | IMCC26256 | uncultured.actinobacterium | NA | 0.64 | 0.65 |
|  | IMCC26256 | uncultured.actinobacterium | NA | -0.62 | -0.56 |
|  | IMCC26256 | uncultured.actinobacterium | NA | -0.56 | -0.56 |
|  | IMCC26256 | uncultured.actinobacterium | NA | -0.52 |  |
|  | IMCC26256 | uncultured.bacterium | NA | 0.72 | 0.78 |
|  | IMCC26256 | uncultured.bacterium | NA | 0.58 | 0.63 |
|  | IMCC26256 | uncultured.bacterium | NA | 0.59 | 0.64 |
|  | IMCC26256 | uncultured.bacterium | NA | 0.63 | 0.65 |
|  | IMCC26256 | uncultured.bacterium | NA |  | -0.56 |
|  | IMCC26256 | uncultured.bacterium | NA | 0.51 |  |
|  | IMCC26256 | uncultured.bacterium | NA | 0.61 |  |
|  | IMCC26256 | uncultured.bacterium | NA |  | -0.57 |
|  | Microtrichales | Iamiaceae | Iamia | -0.52 |  |
|  | Microtrichales | Iamiaceae | Iamia | -0.56 |  |
|  | Microtrichales | Ilumatobacteraceae | CL500.29.marine.group |  | -0.62 |
|  | Microtrichales | Ilumatobacteraceae | Ilumatobacter | -0.51 |  |
|  | Microtrichales | Ilumatobacteraceae | Ilumatobacter | -0.52 |  |
|  | Microtrichales | Ilumatobacteraceae | uncultured | -0.69 | -0.69 |
|  | Microtrichales | Ilumatobacteraceae | uncultured | 0.55 | 0.56 |
|  | Microtrichales | Ilumatobacteraceae | uncultured | -0.60 |  |
|  | Microtrichales | Ilumatobacteraceae | uncultured | -0.52 |  |
|  | Microtrichales | uncultured | metagenome | -0.66 | -0.59 |
|  | Microtrichales | uncultured | uncultured.bacterium | -0.52 |  |
|  | uncultured | uncultured.actinobacterium | NA | -0.51 | -0.58 |
|  | uncultured | uncultured.bacterium | NA | -0.64 | -0.61 |
|  | uncultured | uncultured.bacterium | NA | 0.64 | 0.60 |
|  | uncultured | uncultured.bacterium | NA | 0.56 | 0.55 |
|  | Corynebacteriales | Mycobacteriaceae | Mycobacterium | 0.68 | 0.67 |
|  | Corynebacteriales | Mycobacteriaceae | Mycobacterium |  | -0.52 |
|  | Corynebacteriales | Mycobacteriaceae | Mycobacterium | 0.55 |  |
|  | Corynebacteriales | Nocardiaceae | Nocardia | 0.59 | 0.54 |
|  | Corynebacteriales | Nocardiaceae | Rhodococcus | -0.53 | -0.52 |
|  | Corynebacteriales | Nocardiaceae | Rhodococcus | -0.56 |  |
|  | Corynebacteriales | Nocardiaceae | Rhodococcus |  | -0.55 |
|  | Corynebacteriales | Nocardiaceae | Rhodococcus | -0.64 | -0.56 |
|  | Frankiales | Acidothermaceae | Acidothermus | 0.71 | 0.69 |
|  | Frankiales | Acidothermaceae | Acidothermus | 0.76 | 0.74 |
|  | Frankiales | Acidothermaceae | Acidothermus | 0.67 | 0.66 |
|  | Frankiales | Acidothermaceae | Acidothermus | 0.74 | 0.70 |
|  | Frankiales | Acidothermaceae | Acidothermus | 0.66 | 0.64 |
|  | Frankiales | Acidothermaceae | Acidothermus | 0.51 |  |
|  | Frankiales | Acidothermaceae | Acidothermus | 0.71 | 0.70 |
|  | Frankiales | Acidothermaceae | Acidothermus | 0.62 | 0.57 |
|  | Frankiales | Acidothermaceae | Acidothermus | 0.52 |  |
|  | Frankiales | Acidothermaceae | Acidothermus | 0.63 | 0.62 |
|  | Frankiales | Acidothermaceae | Acidothermus | 0.56 | 0.57 |
|  | Frankiales | Acidothermaceae | Acidothermus | 0.63 | 0.57 |
|  | Frankiales | Acidothermaceae | Acidothermus | 0.69 | 0.67 |
|  | Frankiales | Acidothermaceae | Acidothermus | 0.64 | 0.54 |
|  | Frankiales | Acidothermaceae | Acidothermus | 0.58 | 0.60 |
|  | Frankiales | Acidothermaceae | Acidothermus | 0.52 | 0.55 |
|  | Frankiales | Acidothermaceae | Acidothermus | 0.54 |  |
|  | Frankiales | Acidothermaceae | Acidothermus | -0.55 |  |
|  | Frankiales | Acidothermaceae | Acidothermus | 0.55 |  |
|  | Frankiales | Acidothermaceae | Acidothermus | 0.51 |  |
|  | Frankiales | Frankiaceae | Frankia | 0.58 | 0.54 |
|  | Frankiales | Frankiaceae | Jatrophihabitans |  | 0.53 |
|  | Frankiales | Frankiaceae | Jatrophihabitans | 0.69 | 0.65 |
|  | Frankiales | Frankiaceae | Jatrophihabitans | -0.53 |  |
|  | Frankiales | Geodermatophilaceae | Blastococcus | 0.59 | 0.61 |
|  | Frankiales | Geodermatophilaceae | Blastococcus | 0.52 | 0.54 |
|  | Frankiales | Geodermatophilaceae | Blastococcus | -0.59 | -0.57 |
|  | Frankiales | Geodermatophilaceae | Blastococcus | -0.55 | -0.55 |
|  | Frankiales | Geodermatophilaceae | Klenkia | -0.58 | -0.56 |
|  | Frankiales | Nakamurellaceae | Nakamurella | -0.51 |  |
|  | Frankiales | Nakamurellaceae | Nakamurella | -0.74 | -0.68 |
|  | Frankiales | Nakamurellaceae | Nakamurella |  | -0.52 |
|  | Frankiales | Sporichthyaceae | uncultured | -0.54 | -0.50 |
|  | Micrococcales | Cellulomonadaceae | Cellulomonas | -0.63 | -0.61 |
|  | Micrococcales | Cellulomonadaceae | Cellulomonas | -0.61 |  |
|  | Micrococcales | Intrasporangiaceae | Kribbia | -0.76 | -0.70 |
|  | Micrococcales | Intrasporangiaceae | Oryzihumus |  | 0.51 |
|  | Micrococcales | Intrasporangiaceae | Pedococcus.Phycicoccus | -0.68 | -0.68 |
|  | Micrococcales | Microbacteriaceae | Leifsonia | -0.50 | -0.62 |
|  | Micrococcales | Microbacteriaceae | Microbacterium |  | -0.51 |
|  | Micrococcales | Microbacteriaceae | Plantibacter | -0.52 | -0.52 |
|  | Micromonosporales | Micromonosporaceae | Actinoplanes | -0.53 | -0.52 |
|  | Micromonosporales | Micromonosporaceae | Catellatospora | -0.68 |  |
|  | Micromonosporales | Micromonosporaceae | Dactylosporangium | -0.53 |  |
|  | Micromonosporales | Micromonosporaceae | Micromonospora | 0.52 |  |
|  | Micromonosporales | Micromonosporaceae | Rhizocola | -0.61 | -0.51 |
|  | Micromonosporales | Micromonosporaceae | Salinispora | -0.50 |  |
|  | Propionibacteriales | Nocardioidaceae | Marmoricola |  | 0.53 |
|  | Propionibacteriales | Nocardioidaceae | Nocardioides | 0.55 | 0.58 |
|  | Propionibacteriales | Nocardioidaceae | Nocardioides | -0.75 | -0.72 |
|  | Propionibacteriales | Nocardioidaceae | Nocardioides | -0.60 | -0.51 |
|  | Propionibacteriales | Nocardioidaceae | Nocardioides | -0.61 | -0.56 |
|  | Propionibacteriales | Nocardioidaceae | Nocardioides | -0.69 | -0.67 |
|  | Propionibacteriales | Nocardioidaceae | Nocardioides | -0.64 | -0.57 |
|  | Propionibacteriales | Nocardioidaceae | Nocardioides |  | -0.52 |
|  | Propionibacteriales | Nocardioidaceae | Nocardioides | -0.50 |  |
|  | Propionibacteriales | Nocardioidaceae | Nocardioides | -0.68 | -0.64 |
|  | Propionibacteriales | Nocardioidaceae | Nocardioides | -0.61 | -0.60 |
|  | Propionibacteriales | Nocardioidaceae | Nocardioides | 0.63 | 0.57 |
|  | Propionibacteriales | Nocardioidaceae | Nocardioides |  | -0.59 |
|  | Propionibacteriales | Propionibacteriaceae | Microlunatus | -0.57 | -0.59 |
|  | Pseudonocardiales | Pseudonocardiaceae | Pseudonocardia | -0.69 | -0.74 |
|  | Pseudonocardiales | Pseudonocardiaceae | Pseudonocardia | -0.69 | -0.69 |
|  | Pseudonocardiales | Pseudonocardiaceae | Pseudonocardia | -0.55 | -0.56 |
|  | Streptomycetales | Streptomycetaceae | Streptomyces | 0.64 | 0.60 |
|  | Streptosporangiales | Streptosporangiaceae | Sphaerimonospora | -0.51 | -0.54 |
|  | Streptosporangiales | Streptosporangiaceae | Streptosporangium | -0.55 | -0.55 |
|  | Streptosporangiales | Streptosporangiaceae | Thermopolyspora | -0.51 |  |
|  | metagenome | NA | NA | -0.73 | -0.72 |
|  | uncultured.actinobacterium | NA | NA | -0.59 | -0.64 |
|  | uncultured.bacterium | NA | NA | 0.57 | 0.56 |
|  | uncultured.bacterium | NA | NA | -0.70 | -0.67 |
|  | uncultured.bacterium | NA | NA | -0.51 |  |
|  | uncultured.bacterium | NA | NA | -0.68 | -0.62 |
|  | uncultured.bacterium | NA | NA | -0.59 | -0.55 |
|  | uncultured.bacterium | NA | NA | 0.57 | 0.57 |
|  | uncultured.Catenulispora.sp. | NA | NA |  | -0.53 |
|  | uncultured.soil.bacterium | NA | NA | -0.60 | -0.55 |
|  | Gaiellales | Gaiellaceae | Gaiella | -0.52 |  |
|  | Gaiellales | Gaiellaceae | Gaiella | -0.65 | -0.64 |
|  | Gaiellales | Gaiellaceae | Gaiella | -0.75 | -0.75 |
|  | Gaiellales | Gaiellaceae | Gaiella | -0.67 | -0.65 |
|  | Gaiellales | Gaiellaceae | Gaiella | -0.76 | -0.73 |
|  | Gaiellales | Gaiellaceae | Gaiella | -0.67 | -0.65 |
|  | Gaiellales | uncultured | bacterium.Ellin6517 | 0.69 | 0.70 |
|  | Gaiellales | uncultured | metagenome | 0.66 | 0.65 |
|  | Gaiellales | uncultured | metagenome | -0.57 | -0.52 |
|  | Gaiellales | uncultured | uncultured.actinobacterium | 0.64 | 0.68 |
|  | Gaiellales | uncultured | uncultured.actinobacterium | 0.69 | 0.68 |
|  | Gaiellales | uncultured | uncultured.bacterium | 0.68 | 0.71 |
|  | Gaiellales | uncultured | uncultured.bacterium | 0.57 | 0.63 |
|  | Gaiellales | uncultured | uncultured.bacterium | 0.70 | 0.73 |
|  | Gaiellales | uncultured | uncultured.bacterium | -0.53 |  |
|  | Gaiellales | uncultured | uncultured.bacterium | -0.73 | -0.72 |
|  | Gaiellales | uncultured | uncultured.bacterium | -0.73 | -0.71 |
|  | Gaiellales | uncultured | uncultured.bacterium | 0.70 | 0.69 |
|  | Gaiellales | uncultured | uncultured.bacterium | 0.55 | 0.55 |
|  | Gaiellales | uncultured | uncultured.bacterium | 0.59 | 0.55 |
|  | Gaiellales | uncultured | uncultured.bacterium | 0.61 | 0.57 |
|  | Gaiellales | uncultured | uncultured.bacterium | -0.67 | -0.60 |
|  | Gaiellales | uncultured | uncultured.bacterium | 0.68 | 0.68 |
|  | Gaiellales | uncultured | uncultured.bacterium | 0.72 | 0.67 |
|  | Gaiellales | uncultured | uncultured.bacterium | 0.64 | 0.63 |
|  | Gaiellales | uncultured | uncultured.bacterium |  | -0.53 |
|  | Gaiellales | uncultured | uncultured.bacterium | -0.55 |  |
|  | Gaiellales | uncultured | uncultured.bacterium | -0.62 | -0.61 |
|  | Gaiellales | uncultured | uncultured.bacterium | -0.71 | -0.55 |
|  | Gaiellales | uncultured | uncultured.bacterium | -0.50 |  |
|  | Gaiellales | uncultured | uncultured.bacterium |  | 0.50 |
|  | Gaiellales | uncultured | uncultured.Conexibacteraceae.bacterium | 0.68 | 0.65 |
|  | Gaiellales | uncultured | uncultured.Conexibacteraceae.bacterium | 0.53 | 0.50 |
|  | Gaiellales | uncultured | uncultured.Rubrobacteria.bacterium | -0.59 | -0.58 |
|  | Solirubrobacterales | 67.14 | metagenome | 0.62 | 0.54 |
|  | Solirubrobacterales | 67.14 | metagenome | -0.51 |  |
|  | Solirubrobacterales | 67.14 | uncultured.actinobacterium | -0.57 | -0.58 |
|  | Solirubrobacterales | 67.14 | uncultured.bacterium | 0.57 | 0.55 |
|  | Solirubrobacterales | 67.14 | uncultured.bacterium | 0.50 |  |
|  | Solirubrobacterales | 67.14 | uncultured.bacterium | 0.54 |  |
|  | Solirubrobacterales | 67.14 | uncultured.bacterium | -0.51 |  |
|  | Solirubrobacterales | 67.14 | uncultured.bacterium | -0.55 | -0.54 |
|  | Solirubrobacterales | 67.14 | uncultured.bacterium | -0.52 | -0.52 |
|  | Solirubrobacterales | 67.14 | uncultured.bacterium | -0.55 | -0.53 |
|  | Solirubrobacterales | 67.14 | uncultured.bacterium |  | -0.51 |
|  | Solirubrobacterales | 67.14 | uncultured.bacterium | -0.66 | -0.63 |
|  | Solirubrobacterales | 67.14 | uncultured.bacterium | -0.61 | -0.55 |
|  | Solirubrobacterales | 67.14 | uncultured.bacterium | -0.57 | -0.56 |
|  | Solirubrobacterales | Solirubrobacteraceae | Conexibacter | 0.60 | 0.60 |
|  | Solirubrobacterales | Solirubrobacteraceae | Conexibacter |  | -0.50 |
|  | Solirubrobacterales | Solirubrobacteraceae | Conexibacter | 0.57 | 0.60 |
|  | Solirubrobacterales | Solirubrobacteraceae | Conexibacter |  | 0.58 |
|  | Solirubrobacterales | Solirubrobacteraceae | Conexibacter | 0.53 |  |
|  | Solirubrobacterales | Solirubrobacteraceae | Conexibacter | -0.50 | -0.58 |
|  | Solirubrobacterales | Solirubrobacteraceae | Solirubrobacter | -0.56 |  |
|  | Solirubrobacterales | Solirubrobacteraceae | Solirubrobacter | -0.58 |  |
|  | Solirubrobacterales | Solirubrobacteraceae | Solirubrobacter | -0.54 | -0.56 |
|  | Solirubrobacterales | Solirubrobacteraceae | Solirubrobacter | -0.67 | -0.64 |
|  | Solirubrobacterales | Solirubrobacteraceae | Solirubrobacter | -0.71 | -0.72 |
|  | Solirubrobacterales | Solirubrobacteraceae | Solirubrobacter | -0.66 | -0.56 |
|  | Solirubrobacterales | Solirubrobacteraceae | uncultured | 0.68 | 0.70 |
|  | Solirubrobacterales | Solirubrobacteraceae | uncultured | 0.65 | 0.62 |
|  | Solirubrobacterales | Solirubrobacteraceae | uncultured |  | -0.51 |
|  | Solirubrobacterales | Solirubrobacteraceae | uncultured | -0.52 |  |
|  | uncultured | uncultured.bacterium | NA | -0.61 | -0.55 |
|  | uncultured | uncultured.bacterium | NA | -0.55 | -0.50 |
|  | uncultured | uncultured.bacterium | NA | -0.52 | -0.53 |
|  | uncultured | uncultured.bacterium | NA | -0.60 | -0.59 |
| Armatimonadota | Chthonomonadales | Chthonomonadaceae | Chthonomonas | 0.60 | 0.57 |
|  | Chthonomonadales | Chthonomonadaceae | Chthonomonas | 0.60 | 0.54 |
|  | Chthonomonadales | Chthonomonadaceae | Chthonomonas | 0.51 |  |
|  | Fimbriimonadales | Fimbriimonadaceae | metagenome |  | -0.58 |
|  | Fimbriimonadales | Fimbriimonadaceae | uncultured.bacterium | -0.62 | -0.55 |
|  | Fimbriimonadales | Fimbriimonadaceae | uncultured.bacterium |  | -0.52 |
|  | uncultured.actinobacterium | NA | NA | -0.51 |  |
|  | uncultured.bacterium | NA | NA | -0.58 | -0.54 |
|  | uncultured.bacterium | NA | NA |  | -0.51 |
|  | uncultured.Firmicutes.bacterium | NA | NA | -0.52 |  |
| Bacteroidota | Chitinophagales | Chitinophagaceae | Ferruginibacter | 0.69 | 0.70 |
|  | Chitinophagales | Chitinophagaceae | Ferruginibacter | -0.59 | -0.59 |
|  | Chitinophagales | Chitinophagaceae | Flavisolibacter |  | 0.55 |
|  | Chitinophagales | Chitinophagaceae | Flavisolibacter | 0.66 | 0.68 |
|  | Chitinophagales | Chitinophagaceae | Flavisolibacter | 0.55 | 0.64 |
|  | Chitinophagales | Chitinophagaceae | Niastella | -0.60 | -0.62 |
|  | Chitinophagales | Chitinophagaceae | Parafilimonas | 0.51 |  |
|  | Chitinophagales | Chitinophagaceae | Parafilimonas |  | 0.52 |
|  | Chitinophagales | Chitinophagaceae | Puia | 0.63 | 0.62 |
|  | Chitinophagales | Chitinophagaceae | Puia | 0.75 | 0.74 |
|  | Chitinophagales | Chitinophagaceae | Puia | 0.60 | 0.54 |
|  | Chitinophagales | Chitinophagaceae | Puia | 0.72 | 0.68 |
|  | Chitinophagales | Chitinophagaceae | Terrimonas | 0.59 | 0.52 |
|  | Chitinophagales | Chitinophagaceae | Terrimonas | 0.59 | 0.53 |
|  | Chitinophagales | Chitinophagaceae | Terrimonas |  | -0.51 |
|  | Chitinophagales | Chitinophagaceae | uncultured | 0.72 | 0.71 |
|  | Chitinophagales | Chitinophagaceae | uncultured | 0.69 | 0.73 |
|  | Chitinophagales | Chitinophagaceae | uncultured | 0.52 | 0.55 |
|  | Chitinophagales | Chitinophagaceae | uncultured | 0.65 | 0.58 |
|  | Chitinophagales | Chitinophagaceae | uncultured |  | 0.51 |
|  | Chitinophagales | Chitinophagaceae | uncultured |  | 0.51 |
|  | Chitinophagales | Chitinophagaceae | uncultured | -0.51 |  |
|  | Chitinophagales | Chitinophagaceae | uncultured | -0.54 | -0.53 |
|  | Chitinophagales | Chitinophagaceae | UTBCD1 | 0.51 |  |
|  | Chitinophagales | uncultured | uncultured.bacterium | 0.55 | 0.60 |
|  | Chitinophagales | uncultured | uncultured.bacterium | 0.51 |  |
|  | Cytophagales | Microscillaceae | Chryseolinea | -0.53 | -0.57 |
|  | Cytophagales | Microscillaceae | uncultured | -0.57 | -0.52 |
|  | Cytophagales | Microscillaceae | uncultured | -0.56 | -0.56 |
|  | Cytophagales | Microscillaceae | uncultured | -0.51 |  |
|  | Cytophagales | Spirosomaceae | Fibrella |  | -0.56 |
|  | Cytophagales | Spirosomaceae | Spirosoma |  | -0.55 |
|  | Cytophagales | uncultured | uncultured.bacterium | -0.51 | -0.53 |
|  | Cytophagales | uncultured | uncultured.bacterium |  | -0.54 |
|  | Flavobacteriales | Flavobacteriaceae | Flavobacterium |  | 0.53 |
|  | Sphingobacteriales | AKYH767 | uncultured.bacterium |  | -0.58 |
|  | Sphingobacteriales | FFCH9454 | uncultured.bacterium | -0.63 | -0.55 |
|  | Sphingobacteriales | FFCH9454 | uncultured.bacterium | -0.51 |  |
|  | Sphingobacteriales | Sphingobacteriaceae | Mucilaginibacter |  | 0.60 |
|  | Sphingobacteriales | Sphingobacteriaceae | Mucilaginibacter | 0.50 |  |
|  | Sphingobacteriales | Sphingobacteriaceae | Mucilaginibacter | 0.51 | 0.58 |
|  | Sphingobacteriales | Sphingobacteriaceae | Mucilaginibacter |  | 0.52 |
|  | Sphingobacteriales | Sphingobacteriaceae | Pedobacter | -0.67 | -0.56 |
|  | Sphingobacteriales | Sphingobacteriaceae | uncultured |  | 0.51 |
|  | Sphingobacteriales | Sphingobacteriaceae | uncultured | -0.58 | -0.58 |
|  | Sphingobacteriales | Sphingobacteriaceae | uncultured |  | -0.51 |
|  | Kapabacteriales | uncultured.bacterium | NA | -0.54 |  |
|  | Kryptoniales | BSV26 | uncultured.bacterium | -0.57 | -0.52 |
| Bdellovibrionota | Bdellovibrionales | Bdellovibrionaceae | OM27.clade | -0.59 | -0.54 |
|  | Bdellovibrionales | Bdellovibrionaceae | OM27.clade | -0.52 | -0.54 |
|  | 0319.6G20 | uncultured.bacterium | NA | -0.61 |  |
|  | 0319.6G20 | uncultured.Syntrophobacteraceae.bacterium | NA | -0.53 | -0.52 |
|  | Oligoflexales | Oligoflexaceae | Oligoflexus | -0.75 | -0.64 |
| Chloroflexota syn. Chloroflexi | uncultured.bacterium | NA | NA | -0.55 | -0.56 |
|  | uncultured.Chloroflexi.bacterium | NA | NA | 0.71 | 0.69 |
|  | Anaerolineales | Anaerolineaceae | uncultured | -0.55 | -0.59 |
|  | Anaerolineales | Anaerolineaceae | uncultured | 0.54 |  |
|  | Anaerolineales | Anaerolineaceae | uncultured | -0.50 |  |
|  | Anaerolineales | Anaerolineaceae | uncultured | -0.51 | -0.50 |
|  | Ardenticatenales | uncultured | uncultured.bacterium | -0.51 | -0.52 |
|  | Caldilineales | Caldilineaceae | uncultured | -0.61 | -0.52 |
|  | Caldilineales | Caldilineaceae | uncultured | -0.52 |  |
|  | RBG.13.54.9 | uncultured.bacterium | NA | -0.59 | -0.56 |
|  | RBG.13.54.9 | uncultured.bacterium | NA | -0.59 | -0.58 |
|  | RBG.13.54.9 | uncultured.bacterium | NA | -0.54 | -0.56 |
|  | RBG.13.54.9 | uncultured.bacterium | NA | -0.51 |  |
|  | RBG.13.54.9 | uncultured.bacterium | NA | -0.59 | -0.61 |
|  | RBG.13.54.9 | uncultured.bacterium | NA | -0.52 | -0.50 |
|  | SBR1031 | A4b | uncultured.bacterium | 0.58 | 0.50 |
|  | SBR1031 | A4b | uncultured.bacterium | 0.56 | 0.51 |
|  | SBR1031 | A4b | uncultured.bacterium | -0.60 |  |
|  | SBR1031 | A4b | uncultured.bacterium | -0.56 | -0.52 |
|  | SBR1031 | A4b | uncultured.bacterium | -0.53 | -0.52 |
|  | SBR1031 | A4b | uncultured.soil.bacterium |  | -0.51 |
|  | SBR1031 | metagenome | NA | -0.52 |  |
|  | SBR1031 | uncultured.bacterium | NA | 0.54 |  |
|  | SBR1031 | uncultured.bacterium | NA |  | 0.57 |
|  | SBR1031 | uncultured.bacterium | NA |  | -0.51 |
|  | SBR1031 | uncultured.bacterium | NA |  | -0.52 |
|  | SBR1031 | uncultured.bacterium | NA | -0.53 | -0.59 |
|  | SBR1031 | uncultured.bacterium | NA |  | -0.54 |
|  | SBR1031 | uncultured.Chloroflexi.bacterium | NA |  | -0.60 |
|  | SBR1031 | uncultured.Gemmatimonadetes.bacterium | NA |  | -0.52 |
|  | SBR1031 | wastewater.metagenome | NA |  | -0.53 |
|  | Chloroflexales | Chloroflexaceae | FFCH7168 | -0.65 | -0.63 |
|  | Chloroflexales | Herpetosiphonaceae | Herpetosiphon | -0.53 |  |
|  | Chloroflexales | Roseiflexaceae | uncultured | -0.63 | -0.65 |
|  | Chloroflexales | Roseiflexaceae | uncultured | -0.53 | -0.57 |
|  | Chloroflexales | Roseiflexaceae | uncultured | -0.66 | -0.66 |
|  | Chloroflexales | Roseiflexaceae | uncultured | -0.68 | -0.66 |
|  | Chloroflexales | Roseiflexaceae | uncultured | -0.56 |  |
|  | Chloroflexales | Roseiflexaceae | uncultured | -0.55 | -0.59 |
|  | Chloroflexales | Roseiflexaceae | uncultured | -0.61 | -0.55 |
|  | Chloroflexales | Roseiflexaceae | uncultured | -0.60 | -0.56 |
|  | Chloroflexales | Roseiflexaceae | uncultured | -0.63 | -0.61 |
|  | Chloroflexales | Roseiflexaceae | uncultured |  | -0.51 |
|  | Elev.1554 | uncultured.bacterium | NA | 0.55 | 0.50 |
|  | Thermomicrobiales | AKYG1722 | uncultured.bacterium | -0.54 |  |
|  | Thermomicrobiales | AKYG1722 | uncultured.Chloroflexi.bacterium | -0.59 | -0.60 |
|  | Thermomicrobiales | JG30.KF.CM45 | uncultured.bacterium | -0.54 |  |
|  | Thermomicrobiales | JG30.KF.CM45 | uncultured.bacterium | -0.57 | -0.62 |
|  | Thermomicrobiales | JG30.KF.CM45 | uncultured.bacterium | -0.51 |  |
|  | Thermomicrobiales | JG30.KF.CM45 | uncultured.bacterium | -0.61 | -0.53 |
|  | Thermomicrobiales | JG30.KF.CM45 | uncultured.bacterium | -0.63 | -0.59 |
|  | Thermomicrobiales | JG30.KF.CM45 | uncultured.bacterium | -0.53 |  |
|  | S085 | uncultured.bacterium | NA | -0.61 | -0.55 |
|  | S085 | uncultured.bacterium | NA | -0.54 | -0.54 |
|  | S085 | uncultured.bacterium.5G4 | NA | 0.59 | 0.59 |
|  | uncultured.bacterium | NA | NA | -0.60 | -0.58 |
|  | uncultured.Anaerolineales.bacterium | NA | NA | -0.51 |  |
|  | uncultured.bacterium | NA | NA | 0.66 | 0.65 |
|  | uncultured.bacterium | NA | NA | -0.59 | -0.55 |
|  | uncultured.bacterium | NA | NA | -0.51 | -0.51 |
|  | uncultured.bacterium | NA | NA | 0.55 |  |
|  | uncultured.Caldilinea.sp. | NA | NA | 0.65 | 0.63 |
|  | uncultured.soil.bacterium | NA | NA | -0.53 |  |
|  | uncultured.bacterium | NA | NA | -0.56 |  |
|  | uncultured.bacterium | NA | NA | 0.78 | 0.75 |
|  | uncultured.bacterium | NA | NA | -0.67 | -0.61 |
|  | uncultured.bacterium | NA | NA | -0.67 | -0.62 |
|  | uncultured.bacterium | NA | NA |  | 0.51 |
|  | uncultured.bacterium | NA | NA | -0.51 | -0.61 |
|  | uncultured.bacterium | NA | NA | -0.52 |  |
|  | uncultured.bacterium | NA | NA | -0.61 | -0.50 |
|  | uncultured.Chloroflexi.bacterium | NA | NA | -0.66 | -0.66 |
|  | uncultured.Chloroflexi.bacterium | NA | NA | -0.58 | -0.56 |
|  | uncultured.Longilinea.sp. | NA | NA | 0.59 | 0.58 |
|  | C0119 | uncultured.candidate.division.SAM.bacterium | NA | 0.61 | 0.56 |
|  | C0119 | uncultured.Chloroflexi.bacterium | NA | 0.57 | 0.61 |
|  | C0119 | uncultured.Chloroflexi.bacterium | NA | -0.54 | -0.53 |
|  | Ktedonobacterales | JG30.KF.AS9 | Chloroflexi.bacterium.Ellin7237 | 0.66 | 0.63 |
|  | Ktedonobacterales | JG30.KF.AS9 | uncultured.bacterium | 0.56 |  |
|  | Ktedonobacterales | JG30.KF.AS9 | uncultured.bacterium | 0.60 | 0.57 |
|  | Ktedonobacterales | JG30.KF.AS9 | uncultured.bacterium | 0.56 | 0.59 |
|  | Ktedonobacterales | JG30.KF.AS9 | uncultured.bacterium | 0.64 | 0.64 |
|  | Ktedonobacterales | JG30.KF.AS9 | uncultured.bacterium | 0.52 | 0.51 |
|  | Ktedonobacterales | JG30.KF.AS9 | uncultured.Thermosporothrix.sp. | 0.54 | 0.50 |
|  | Ktedonobacterales | Ktedonobacteraceae | 1921.2 | 0.67 | 0.66 |
|  | Ktedonobacterales | Ktedonobacteraceae | 1921.2 | 0.71 | 0.72 |
|  | Ktedonobacterales | Ktedonobacteraceae | HSB.OF53.F07 | 0.63 | 0.61 |
|  | Ktedonobacterales | Ktedonobacteraceae | JG30a.KF.32 | 0.62 | 0.63 |
|  | Ktedonobacterales | Ktedonobacteraceae | uncultured | 0.63 | 0.58 |
|  | Ktedonobacterales | Ktedonobacteraceae | uncultured | 0.61 | 0.61 |
|  | uncultured.bacterium | NA | NA | 0.74 | 0.71 |
|  | uncultured.bacterium | NA | NA | -0.69 | -0.72 |
|  | uncultured.bacterium | NA | NA | -0.65 | -0.59 |
|  | uncultured.Chloroflexi.bacterium | NA | NA | -0.70 | -0.63 |
|  | uncultured.bacterium | NA | NA | -0.54 |  |
|  | metagenome | NA | NA | -0.55 | -0.53 |
|  | uncultured.bacterium | NA | NA | 0.64 | 0.58 |
|  | uncultured.bacterium | NA | NA | 0.58 | 0.62 |
|  | uncultured.bacterium | NA | NA | 0.59 | 0.59 |
|  | uncultured.bacterium | NA | NA | -0.54 |  |
|  | uncultured.bacterium | NA | NA | -0.62 | -0.51 |
|  | uncultured.bacterium | NA | NA | -0.59 |  |
|  | uncultured.Chloroflexi.bacterium | NA | NA | 0.51 | 0.52 |
| Cyanobacteria | Chloroplast | uncultured.bacterium | NA | -0.64 | -0.61 |
|  | Chloroplast | uncultured.bacterium | NA | -0.65 | -0.61 |
|  | Chloroplast | uncultured.bacterium | NA |  | -0.56 |
|  | Chloroplast | uncultured.bacterium | NA | -0.51 |  |
|  | Cyanobacteriales | Nostocaceae | Cylindrospermum.PCC.7417 | -0.58 | -0.56 |
|  | Cyanobacteriales | Nostocaceae | Nostoc.PCC.7107 | -0.60 | -0.64 |
|  | Cyanobacteriales | Nostocaceae | Nostoc.PCC.73102 | -0.68 | -0.69 |
|  | Cyanobacteriales | Nostocaceae | Nostoc.PCC.73102 | -0.72 | -0.71 |
|  | Cyanobacteriales | Phormidiaceae | Tychonema.CCAP.1459.11B | -0.55 | -0.53 |
|  | Oxyphotobacteria.Incertae.Sedis | Unknown.Family | Leptolyngbya.EcFYyyy.00 | -0.55 | -0.54 |
| Desulfobacterota | Desulfuromonadales | Desulfuromonadaceae | Desulfuromonas | -0.58 | -0.53 |
|  | Geobacterales | Geobacteraceae | Geobacter |  | -0.55 |
|  | uncultured.bacterium | NA | NA | -0.69 | -0.63 |
| Elusimicrobiota | uncultured.bacterium | NA | NA | 0.57 |  |
|  | uncultured.Chitinophagaceae.bacterium | NA | NA | 0.52 |  |
| Entotheonellaeota | Entotheonellales | Entotheonellaceae | metagenome | -0.57 | -0.60 |
| Fibrobacterota | Fibrobacterales | Fibrobacteraceae | possible.genus.04 |  | -0.50 |
| Bacillota syn. Firmicutes | Alicyclobacillales | Alicyclobacillaceae | Tumebacillus | -0.77 | -0.77 |
|  | Alicyclobacillales | Alicyclobacillaceae | Tumebacillus | -0.66 | -0.65 |
|  | Alicyclobacillales | Alicyclobacillaceae | Tumebacillus |  | -0.51 |
|  | Bacillales | Bacillaceae | Bacillus | -0.58 | -0.58 |
|  | Bacillales | Bacillaceae | Bacillus | -0.61 | -0.52 |
|  | Bacillales | Bacillaceae | Bacillus | -0.53 |  |
|  | Bacillales | Bacillaceae | Bacillus | -0.57 | -0.56 |
|  | Bacillales | Bacillaceae | Bacillus | -0.53 |  |
|  | Bacillales | Bacillaceae | Oceanobacillus | -0.58 | -0.63 |
|  | Bacillales | Planococcaceae | Sporosarcina | -0.56 | -0.52 |
|  | Bacillales | Planococcaceae | Sporosarcina | -0.52 |  |
|  | Paenibacillales | Paenibacillaceae | Cohnella |  | -0.52 |
|  | Paenibacillales | Paenibacillaceae | Cohnella | -0.51 |  |
|  | Paenibacillales | Paenibacillaceae | Paenibacillus | -0.54 | -0.56 |
|  | Paenibacillales | Paenibacillaceae | Paenibacillus |  | -0.54 |
|  | Paenibacillales | Paenibacillaceae | Thermobacillus | -0.56 |  |
|  | Thermoactinomycetales | Thermoactinomycetaceae | Novibacillus |  | -0.53 |
|  | Thermoactinomycetales | Thermoactinomycetaceae | Planifilum | -0.58 |  |
|  | uncultured | uncultured.bacterium | NA | -0.53 |  |
|  | Clostridiales | Clostridiaceae | Clostridium.sensu.stricto.12 | 0.54 |  |
|  | Peptostreptococcales.Tissierellales | Peptostreptococcaceae | Paraclostridium | -0.61 |  |
|  | Limnochordales | Limnochordaceae | uncultured.bacterium | -0.50 |  |
|  | Limnochordales | Limnochordaceae | uncultured.bacterium |  | -0.51 |
|  | Limnochordales | Limnochordaceae | uncultured.compost.bacterium | -0.58 |  |
|  | Limnochordales | Limnochordaceae | uncultured.compost.bacterium | -0.51 | -0.52 |
|  | Symbiobacteriales | Symbiobacteraceae | Symbiobacterium | -0.65 | -0.51 |
|  | Thermacetogeniales | Thermacetogeniaceae | Syntrophaceticus | 0.59 | 0.51 |
| GAL15 | NA | NA | NA | -0.60 |  |
| Gemmatimonadota | Gemmatimonadales | Gemmatimonadaceae | Gemmatimonas | 0.64 | 0.59 |
|  | Gemmatimonadales | Gemmatimonadaceae | Gemmatimonas | 0.68 | 0.64 |
|  | Gemmatimonadales | Gemmatimonadaceae | Gemmatimonas | 0.51 | 0.58 |
|  | Gemmatimonadales | Gemmatimonadaceae | Gemmatimonas | 0.66 | 0.63 |
|  | Gemmatimonadales | Gemmatimonadaceae | Gemmatimonas | 0.67 | 0.65 |
|  | Gemmatimonadales | Gemmatimonadaceae | Gemmatimonas | 0.58 | 0.66 |
|  | Gemmatimonadales | Gemmatimonadaceae | Gemmatimonas | 0.69 | 0.71 |
|  | Gemmatimonadales | Gemmatimonadaceae | Gemmatimonas | -0.50 |  |
|  | Gemmatimonadales | Gemmatimonadaceae | Gemmatimonas | 0.56 | 0.53 |
|  | Gemmatimonadales | Gemmatimonadaceae | Gemmatimonas | 0.56 | 0.54 |
|  | Gemmatimonadales | Gemmatimonadaceae | Gemmatimonas | -0.51 |  |
|  | Gemmatimonadales | Gemmatimonadaceae | Gemmatimonas | -0.52 |  |
|  | Gemmatimonadales | Gemmatimonadaceae | Gemmatimonas | -0.52 |  |
|  | Gemmatimonadales | Gemmatimonadaceae | Gemmatimonas |  | 0.54 |
|  | Gemmatimonadales | Gemmatimonadaceae | Gemmatimonas | -0.52 | -0.58 |
|  | Gemmatimonadales | Gemmatimonadaceae | Gemmatirosa |  | 0.56 |
|  | Gemmatimonadales | Gemmatimonadaceae | uncultured | -0.66 | -0.66 |
|  | Gemmatimonadales | Gemmatimonadaceae | uncultured | 0.75 | 0.79 |
|  | Gemmatimonadales | Gemmatimonadaceae | uncultured | -0.67 |  |
|  | Gemmatimonadales | Gemmatimonadaceae | uncultured | -0.68 | -0.66 |
|  | Gemmatimonadales | Gemmatimonadaceae | uncultured | 0.68 | 0.68 |
|  | Gemmatimonadales | Gemmatimonadaceae | uncultured | -0.64 | -0.64 |
|  | Gemmatimonadales | Gemmatimonadaceae | uncultured | -0.67 | -0.68 |
|  | Gemmatimonadales | Gemmatimonadaceae | uncultured | -0.69 | -0.61 |
|  | Gemmatimonadales | Gemmatimonadaceae | uncultured | -0.68 | -0.67 |
|  | Gemmatimonadales | Gemmatimonadaceae | uncultured | -0.64 | -0.62 |
|  | Gemmatimonadales | Gemmatimonadaceae | uncultured |  | 0.58 |
|  | Gemmatimonadales | Gemmatimonadaceae | uncultured | -0.54 | -0.52 |
|  | Gemmatimonadales | Gemmatimonadaceae | uncultured | -0.53 |  |
|  | Gemmatimonadales | Gemmatimonadaceae | uncultured | 0.56 |  |
|  | Gemmatimonadales | Gemmatimonadaceae | uncultured | -0.53 |  |
|  | Gemmatimonadales | Gemmatimonadaceae | uncultured |  | 0.50 |
|  | Gemmatimonadales | Gemmatimonadaceae | uncultured | -0.70 | -0.65 |
|  | Gemmatimonadales | Gemmatimonadaceae | uncultured | -0.63 | -0.68 |
|  | Gemmatimonadales | Gemmatimonadaceae | uncultured | -0.63 | -0.59 |
|  | Gemmatimonadales | Gemmatimonadaceae | uncultured |  | -0.52 |
|  | Gemmatimonadales | Gemmatimonadaceae | uncultured | -0.53 |  |
|  | Gemmatimonadales | Gemmatimonadaceae | uncultured | 0.51 |  |
|  | Longimicrobiales | Longimicrobiaceae | uncultured.bacterium | -0.53 |  |
|  | uncultured.bacterium | NA | NA |  | 0.52 |
| Latescibacterota | Latescibacterales | Latescibacteraceae | uncultured.bacterium | -0.52 | -0.57 |
|  | NA | NA | NA | -0.50 |  |
|  | NA | NA | NA |  | -0.50 |
|  | NA | NA | NA | -0.64 | -0.62 |
|  | NA | NA | NA | -0.54 |  |
| MBNT15 | NA | NA | NA | -0.50 |  |
|  | NA | NA | NA |  | -0.56 |
|  | NA | NA | NA | -0.54 | -0.51 |
|  | NA | NA | NA | -0.56 |  |
| Methylomirabilota | Rokubacteriales | uncultured.bacterium | NA | -0.51 | -0.52 |
|  | Rokubacteriales | uncultured.bacterium | NA | -0.72 | -0.72 |
|  | Rokubacteriales | uncultured.bacterium | NA | -0.66 | -0.58 |
|  | Rokubacteriales | uncultured.bacterium | NA | -0.63 | -0.56 |
|  | Rokubacteriales | uncultured.bacterium | NA | -0.67 | -0.53 |
|  | Rokubacteriales | uncultured.bacterium | NA | -0.62 | -0.53 |
|  | Rokubacteriales | uncultured.bacterium | NA | -0.52 |  |
|  | Rokubacteriales | uncultured.Firmicutes.bacterium | NA | -0.64 | -0.58 |
|  | Rokubacteriales | WX65 | bacterium.WX65 | -0.70 | -0.71 |
|  | Rokubacteriales | WX65 | uncultured.bacterium | -0.51 | -0.55 |
|  | Rokubacteriales | WX65 | uncultured.bacterium | -0.55 |  |
| Myxococcota | uncultured.bacterium | NA | NA | -0.66 | -0.69 |
|  | uncultured.bacterium | NA | NA | -0.51 |  |
|  | Blfdi19 | uncultured.delta.proteobacterium | NA | -0.53 |  |
|  | Haliangiales | Haliangiaceae | Haliangium | 0.50 | 0.55 |
|  | Haliangiales | Haliangiaceae | Haliangium |  | 0.51 |
|  | Haliangiales | Haliangiaceae | Haliangium | 0.59 | 0.64 |
|  | Haliangiales | Haliangiaceae | Haliangium |  | -0.51 |
|  | Haliangiales | Haliangiaceae | Haliangium | -0.60 |  |
|  | Haliangiales | Haliangiaceae | Haliangium |  | -0.53 |
|  | Haliangiales | Haliangiaceae | Haliangium |  | -0.51 |
|  | Haliangiales | Haliangiaceae | Haliangium |  | -0.54 |
|  | Haliangiales | Haliangiaceae | Haliangium |  | -0.56 |
|  | Haliangiales | Haliangiaceae | Haliangium | -0.51 | -0.51 |
|  | Nannocystales | Nannocystaceae | metagenome |  | -0.52 |
|  | Nannocystales | Nannocystaceae | Nannocystis | -0.52 | -0.65 |
|  | Nannocystales | Nannocystaceae | Nannocystis | -0.53 |  |
|  | Polyangiales | BIrii41 | uncultured.bacterium |  | -0.55 |
|  | Polyangiales | BIrii41 | uncultured.bacterium | -0.57 |  |
|  | Polyangiales | BIrii41 | uncultured.bacterium | -0.51 |  |
|  | Polyangiales | BIrii41 | uncultured.delta.proteobacterium |  | 0.51 |
|  | Polyangiales | BIrii41 | uncultured.proteobacterium | -0.57 | -0.55 |
|  | Polyangiales | Phaselicystidaceae | Phaselicystis | -0.54 | -0.54 |
|  | Polyangiales | Polyangiaceae | Pajaroellobacter | -0.62 | -0.59 |
|  | Polyangiales | Polyangiaceae | Pajaroellobacter | -0.50 |  |
|  | Polyangiales | Polyangiaceae | Pajaroellobacter | -0.51 |  |
|  | Polyangiales | Polyangiaceae | Pajaroellobacter |  | -0.59 |
|  | Polyangiales | Polyangiaceae | Pajaroellobacter | 0.51 | 0.57 |
|  | Polyangiales | Polyangiaceae | Pajaroellobacter | -0.59 | -0.57 |
|  | Polyangiales | Polyangiaceae | Pajaroellobacter | -0.60 | -0.55 |
|  | Polyangiales | Polyangiaceae | Pajaroellobacter |  | -0.51 |
|  | Polyangiales | Polyangiaceae | Polyangium |  | -0.54 |
|  | Polyangiales | Polyangiaceae | Sorangium | -0.71 | -0.72 |
|  | Polyangiales | Polyangiaceae | Sorangium | -0.58 |  |
|  | Polyangiales | Polyangiaceae | Sorangium | -0.52 |  |
|  | Polyangiales | Polyangiaceae | uncultured | -0.52 |  |
|  | Polyangiales | Polyangiaceae | uncultured |  | -0.55 |
|  | Polyangiales | Polyangiaceae | uncultured |  | -0.57 |
|  | Polyangiales | Sandaracinaceae | uncultured | -0.52 |  |
|  | Polyangiales | Sandaracinaceae | uncultured | 0.53 | 0.51 |
|  | Polyangiales | Sandaracinaceae | uncultured | -0.57 | -0.60 |
|  | Polyangiales | Sandaracinaceae | uncultured | -0.60 | -0.54 |
|  | Polyangiales | Sandaracinaceae | uncultured | -0.63 | -0.53 |
|  | Polyangiales | Sandaracinaceae | uncultured |  | 0.61 |
|  | Polyangiales | Sandaracinaceae | uncultured | -0.52 | -0.55 |
| NB1.j | NA | NA | NA | -0.60 | -0.57 |
| NB1.j | NA | NA | NA | -0.55 | -0.58 |
| Nitrospirota | Nitrospirales | Nitrospiraceae | Nitrospira | -0.66 | -0.63 |
|  | Nitrospirales | Nitrospiraceae | Nitrospira | -0.52 |  |
|  | Nitrospirales | Nitrospiraceae | Nitrospira |  | -0.52 |
|  | Nitrospirales | Nitrospiraceae | Nitrospira | -0.76 | -0.72 |
|  | Nitrospirales | Nitrospiraceae | Nitrospira | -0.58 | -0.51 |
|  | Nitrospirales | Nitrospiraceae | Nitrospira | -0.71 | -0.66 |
|  | Nitrospirales | Nitrospiraceae | Nitrospira | -0.64 | -0.63 |
|  | Nitrospirales | Nitrospiraceae | Nitrospira | -0.56 | -0.55 |
|  | Nitrospirales | Nitrospiraceae | Nitrospira | -0.53 |  |
|  | Nitrospirales | Nitrospiraceae | Nitrospira | -0.57 | -0.52 |
|  | Nitrospirales | Nitrospiraceae | Nitrospira | -0.50 | -0.52 |
| Patescibacteria | Candidatus.Lloydbacteria | Candidatus.Lloydbacteria.bacterium.RIFOXYC12.FULL.46.25 | NA | -0.51 | -0.51 |
|  | Saccharimonadales | LWQ8 | uncultured.bacterium |  | 0.52 |
|  | Saccharimonadales | Saccharimonadaceae | TM7a |  | 0.54 |
|  | Saccharimonadales | uncultured.bacterium | NA |  | 0.53 |
|  | Saccharimonadales | uncultured.bacterium | NA |  | 0.53 |
|  | Saccharimonadales | uncultured.bacterium | NA |  | 0.53 |
|  | Saccharimonadales | uncultured.bacterium | NA | 0.50 | 0.54 |
|  | Saccharimonadales | uncultured.bacterium | NA | -0.57 | -0.56 |
|  | Saccharimonadales | uncultured.bacterium | NA | -0.53 |  |
|  | Saccharimonadales | uncultured.Candidatus.Saccharibacteria.bacterium | NA |  | 0.53 |
|  | Saccharimonadales | uncultured.soil.bacterium | NA | 0.59 | 0.55 |
|  | Saccharimonadales | WWH38 | uncultured.bacterium | 0.60 | 0.60 |
|  | Saccharimonadales | WWH38 | uncultured.Candidatus.Saccharibacteria.bacterium | 0.50 |  |
| Planctomycetota | uncultured.bacterium | NA | NA | -0.56 | -0.53 |
|  | uncultured.bacterium | NA | NA | -0.57 | -0.58 |
|  | uncultured.bacterium | NA | NA | 0.55 | 0.62 |
|  | uncultured.bacterium | NA | NA | -0.65 | -0.62 |
|  | uncultured.bacterium | NA | NA | -0.52 |  |
|  | uncultured.bacterium | NA | NA |  | -0.54 |
|  | uncultured.bacterium | NA | NA | -0.61 | -0.50 |
|  | Phycisphaerales | Phycisphaeraceae | AKYG587 | -0.61 | -0.60 |
|  | Phycisphaerales | Phycisphaeraceae | AKYG587 | -0.57 | -0.58 |
|  | Tepidisphaerales | CPla.3.termite.group | metagenome | 0.67 | 0.62 |
|  | Tepidisphaerales | CPla.3.termite.group | uncultured.planctomycete | 0.55 | 0.67 |
|  | Tepidisphaerales | WD2101.soil.group | Planctomycetales.bacterium.Ellin6207 | 0.67 | 0.69 |
|  | Tepidisphaerales | WD2101.soil.group | planctomycete.WY108 | -0.51 |  |
|  | Tepidisphaerales | WD2101.soil.group | uncultured.bacterium | 0.52 | 0.50 |
|  | Tepidisphaerales | WD2101.soil.group | uncultured.bacterium | 0.54 | 0.61 |
|  | Tepidisphaerales | WD2101.soil.group | uncultured.bacterium | 0.66 | 0.62 |
|  | Tepidisphaerales | WD2101.soil.group | uncultured.bacterium | -0.55 | -0.58 |
|  | Tepidisphaerales | WD2101.soil.group | uncultured.bacterium | 0.66 | 0.59 |
|  | Tepidisphaerales | WD2101.soil.group | uncultured.bacterium | 0.53 |  |
|  | Tepidisphaerales | WD2101.soil.group | uncultured.bacterium | -0.53 |  |
|  | Tepidisphaerales | WD2101.soil.group | uncultured.bacterium |  | 0.57 |
|  | Tepidisphaerales | WD2101.soil.group | uncultured.bacterium | 0.61 | 0.50 |
|  | Tepidisphaerales | WD2101.soil.group | uncultured.bacterium |  | 0.51 |
|  | Tepidisphaerales | WD2101.soil.group | uncultured.bacterium | -0.55 |  |
|  | Tepidisphaerales | WD2101.soil.group | uncultured.bacterium | -0.50 | -0.50 |
|  | Tepidisphaerales | WD2101.soil.group | uncultured.bacterium | 0.58 | 0.57 |
|  | Tepidisphaerales | WD2101.soil.group | uncultured.bacterium | 0.53 |  |
|  | Tepidisphaerales | WD2101.soil.group | uncultured.bacterium |  | -0.57 |
|  | Tepidisphaerales | WD2101.soil.group | uncultured.bacterium |  | -0.52 |
|  | Tepidisphaerales | WD2101.soil.group | uncultured.Planctomycetales.bacterium | 0.53 | 0.55 |
|  | Tepidisphaerales | WD2101.soil.group | uncultured.soil.bacterium | 0.56 | 0.59 |
|  | Tepidisphaerales | WD2101.soil.group | uncultured.soil.bacterium | -0.52 |  |
|  | uncultured.bacterium | NA | NA |  | -0.62 |
|  | metagenome | NA | NA |  | -0.55 |
|  | uncultured.bacterium | NA | NA |  | -0.50 |
|  | Gemmatales | Gemmataceae | Fimbriiglobus |  | -0.56 |
|  | Gemmatales | Gemmataceae | Fimbriiglobus | -0.53 | -0.51 |
|  | Gemmatales | Gemmataceae | Fimbriiglobus |  | -0.51 |
|  | Gemmatales | Gemmataceae | Fimbriiglobus |  | -0.53 |
|  | Gemmatales | Gemmataceae | Fimbriiglobus | -0.52 |  |
|  | Gemmatales | Gemmataceae | Gemmata | 0.64 | 0.62 |
|  | Gemmatales | Gemmataceae | Gemmata | 0.60 | 0.55 |
|  | Gemmatales | Gemmataceae | Gemmata |  | -0.51 |
|  | Gemmatales | Gemmataceae | Gemmata | 0.56 | 0.51 |
|  | Gemmatales | Gemmataceae | Gemmata | -0.51 |  |
|  | Gemmatales | Gemmataceae | uncultured | -0.55 | -0.54 |
|  | Gemmatales | Gemmataceae | uncultured |  | 0.52 |
|  | Gemmatales | Gemmataceae | uncultured | 0.54 |  |
|  | Gemmatales | Gemmataceae | uncultured | 0.52 |  |
|  | Gemmatales | Gemmataceae | uncultured | 0.59 | 0.52 |
|  | Gemmatales | Gemmataceae | uncultured | 0.56 |  |
|  | Gemmatales | Gemmataceae | uncultured |  | 0.51 |
|  | Gemmatales | Gemmataceae | uncultured |  | 0.51 |
|  | Gemmatales | Gemmataceae | uncultured | -0.58 |  |
|  | Gemmatales | Gemmataceae | uncultured | -0.52 | -0.57 |
|  | Gemmatales | Gemmataceae | uncultured | -0.53 | -0.55 |
|  | Gemmatales | Gemmataceae | uncultured | -0.52 |  |
|  | Gemmatales | Gemmataceae | uncultured |  | -0.51 |
|  | Isosphaerales | Isosphaeraceae | Aquisphaera | 0.62 | 0.62 |
|  | Isosphaerales | Isosphaeraceae | Aquisphaera | 0.62 |  |
|  | Isosphaerales | Isosphaeraceae | Aquisphaera | 0.51 |  |
|  | Isosphaerales | Isosphaeraceae | Aquisphaera | 0.66 | 0.57 |
|  | Isosphaerales | Isosphaeraceae | Candidatus.Nostocoida | 0.56 | 0.55 |
|  | Isosphaerales | Isosphaeraceae | Singulisphaera |  | 0.51 |
|  | Isosphaerales | Isosphaeraceae | Singulisphaera | -0.58 | -0.63 |
|  | Isosphaerales | Isosphaeraceae | Singulisphaera | 0.51 | 0.50 |
|  | Isosphaerales | Isosphaeraceae | uncultured | 0.57 | 0.60 |
|  | Isosphaerales | Isosphaeraceae | uncultured |  | 0.55 |
|  | Isosphaerales | Isosphaeraceae | uncultured | -0.51 | -0.51 |
|  | Isosphaerales | Isosphaeraceae | uncultured | 0.57 | 0.58 |
|  | Isosphaerales | Isosphaeraceae | uncultured | -0.59 | -0.61 |
|  | Isosphaerales | Isosphaeraceae | uncultured | -0.64 | -0.67 |
|  | Isosphaerales | Isosphaeraceae | uncultured | 0.62 | 0.57 |
|  | Isosphaerales | Isosphaeraceae | uncultured | 0.59 | 0.54 |
|  | Isosphaerales | Isosphaeraceae | uncultured | 0.53 | 0.58 |
|  | Isosphaerales | Isosphaeraceae | uncultured | 0.58 | 0.57 |
|  | Isosphaerales | Isosphaeraceae | uncultured | -0.54 | -0.55 |
|  | Isosphaerales | Isosphaeraceae | uncultured | -0.51 |  |
|  | Pirellulales | Pirellulaceae | Blastopirellula | -0.54 |  |
|  | Pirellulales | Pirellulaceae | Pir4.lineage | 0.63 | 0.63 |
|  | Pirellulales | Pirellulaceae | Pir4.lineage | 0.63 | 0.60 |
|  | Pirellulales | Pirellulaceae | Pir4.lineage | -0.53 | -0.56 |
|  | Pirellulales | Pirellulaceae | Pir4.lineage | -0.58 |  |
|  | Pirellulales | Pirellulaceae | Pir4.lineage | -0.65 | -0.66 |
|  | Pirellulales | Pirellulaceae | Pir4.lineage | -0.52 | -0.62 |
|  | Pirellulales | Pirellulaceae | Pir4.lineage | -0.52 |  |
|  | Pirellulales | Pirellulaceae | Pir4.lineage | -0.50 |  |
|  | Pirellulales | Pirellulaceae | Pir4.lineage | -0.52 | -0.53 |
|  | Pirellulales | Pirellulaceae | Pirellula | 0.59 | 0.56 |
|  | Pirellulales | Pirellulaceae | Pirellula |  | -0.50 |
|  | Pirellulales | Pirellulaceae | Pirellula | -0.61 | -0.55 |
|  | Pirellulales | Pirellulaceae | Pirellula | -0.52 | -0.51 |
|  | Pirellulales | Pirellulaceae | Pirellula | 0.53 |  |
|  | Pirellulales | Pirellulaceae | Pirellula | -0.66 | -0.57 |
|  | Pirellulales | Pirellulaceae | Pirellula | -0.64 | -0.54 |
|  | Pirellulales | Pirellulaceae | Pirellula | -0.64 | -0.57 |
|  | Pirellulales | Pirellulaceae | Pirellula | -0.63 | -0.62 |
|  | Pirellulales | Pirellulaceae | Pirellula | -0.53 |  |
|  | Pirellulales | Pirellulaceae | uncultured | 0.65 | 0.67 |
|  | Pirellulales | Pirellulaceae | uncultured | 0.58 | 0.53 |
|  | Pirellulales | Pirellulaceae | uncultured | 0.57 | 0.50 |
|  | Pirellulales | Pirellulaceae | uncultured | 0.59 | 0.59 |
|  | Pirellulales | Pirellulaceae | uncultured | -0.53 |  |
|  | Pirellulales | Pirellulaceae | uncultured | 0.64 | 0.64 |
|  | Pirellulales | Pirellulaceae | uncultured | -0.57 | -0.57 |
|  | Pirellulales | Pirellulaceae | uncultured |  | -0.56 |
|  | Pirellulales | Pirellulaceae | uncultured | -0.51 | -0.51 |
|  | Pirellulales | Pirellulaceae | uncultured | 0.52 |  |
|  | Pirellulales | Pirellulaceae | uncultured | -0.66 | -0.59 |
|  | Pirellulales | Pirellulaceae | uncultured | -0.55 | -0.54 |
|  | Pirellulales | Pirellulaceae | uncultured |  | -0.53 |
|  | Pirellulales | Pirellulaceae | uncultured |  | 0.50 |
|  | Planctomycetales | Gimesiaceae | uncultured | 0.59 | 0.56 |
|  | Planctomycetales | Gimesiaceae | uncultured | 0.62 | 0.57 |
|  | Planctomycetales | Rubinisphaeraceae | SH.PL14 | -0.58 | -0.53 |
|  | Planctomycetales | Rubinisphaeraceae | SH.PL14 | -0.63 | -0.60 |
|  | Planctomycetales | Rubinisphaeraceae | SH.PL14 | -0.50 |  |
|  | Planctomycetales | uncultured | metagenome |  | -0.51 |
|  | Planctomycetales | uncultured | uncultured.bacterium | 0.52 | 0.52 |
|  | Planctomycetales | uncultured | uncultured.bacterium |  | -0.56 |
|  | Planctomycetales | uncultured | uncultured.bacterium | 0.57 | 0.54 |
|  | Planctomycetales | uncultured | uncultured.bacterium | -0.62 | -0.70 |
|  | Planctomycetales | uncultured | uncultured.Planctomyces.sp. | 0.50 |  |
|  | uncultured.bacterium | NA | NA | -0.52 |  |
| Pseudomonadota syn. Proteobacteria | Acetobacterales | Acetobacteraceae | Acidisphaera.sp. | 0.52 | 0.50 |
|  | Acetobacterales | Acetobacteraceae | uncultured | 0.55 | 0.56 |
|  | Acetobacterales | Acetobacteraceae | uncultured | 0.61 | 0.60 |
|  | Acetobacterales | Acetobacteraceae | uncultured | 0.72 | 0.71 |
|  | Acetobacterales | Acetobacteraceae | uncultured | 0.54 | 0.56 |
|  | Acetobacterales | Acetobacteraceae | uncultured | 0.64 | 0.57 |
|  | Acetobacterales | Acetobacteraceae | uncultured | 0.63 | 0.60 |
|  | Acetobacterales | Acetobacteraceae | uncultured | -0.67 | -0.69 |
|  | Acetobacterales | Acetobacteraceae | uncultured | 0.52 | 0.51 |
|  | Acetobacterales | Acetobacteraceae | uncultured | 0.60 | 0.64 |
|  | Acetobacterales | Acetobacteraceae | uncultured | 0.75 | 0.65 |
|  | Acetobacterales | Acetobacteraceae | uncultured | 0.59 | 0.59 |
|  | Azospirillales | Azospirillaceae | Skermanella | -0.61 | -0.55 |
|  | Azospirillales | Azospirillaceae | Skermanella | -0.78 | -0.79 |
|  | Caulobacterales | Caulobacteraceae | Phenylobacterium | 0.61 | 0.51 |
|  | Caulobacterales | Caulobacteraceae | Phenylobacterium | -0.57 |  |
|  | Caulobacterales | Caulobacteraceae | uncultured | 0.57 | 0.54 |
|  | Caulobacterales | Caulobacteraceae | uncultured |  | 0.54 |
|  | Caulobacterales | Caulobacteraceae | uncultured | -0.55 |  |
|  | Caulobacterales | Hyphomonadaceae | SWB02 |  | 0.50 |
|  | Dongiales | Dongiaceae | Dongia |  | -0.59 |
|  | Dongiales | Dongiaceae | Dongia | -0.53 | -0.55 |
|  | Dongiales | Dongiaceae | Dongia | -0.51 |  |
|  | Dongiales | Dongiaceae | Dongia | -0.59 | -0.64 |
|  | Elsterales | uncultured | uncultured.Alphaproteobacteria.bacterium | 0.60 |  |
|  | Elsterales | uncultured | uncultured.Alphaproteobacteria.bacterium | 0.62 | 0.62 |
|  | Elsterales | uncultured | uncultured.Alphaproteobacteria.bacterium |  | 0.56 |
|  | Elsterales | uncultured | uncultured.bacterium | 0.57 | 0.50 |
|  | Elsterales | uncultured | uncultured.bacterium | 0.79 | 0.73 |
|  | Elsterales | uncultured | uncultured.bacterium | 0.57 | 0.61 |
|  | Elsterales | uncultured | uncultured.bacterium | 0.65 | 0.64 |
|  | Elsterales | uncultured | uncultured.bacterium | 0.55 | 0.54 |
|  | Elsterales | uncultured | uncultured.bacterium | 0.57 | 0.55 |
|  | Elsterales | uncultured | uncultured.bacterium |  | 0.52 |
|  | Elsterales | uncultured | uncultured.bacterium | -0.52 |  |
|  | Elsterales | uncultured | uncultured.forest.soil.bacterium | 0.66 | 0.66 |
|  | Elsterales | uncultured | uncultured.forest.soil.bacterium | 0.66 | 0.56 |
|  | Elsterales | URHD0088 | uncultured.bacterium | 0.55 | 0.53 |
|  | Elsterales | URHD0088 | uncultured.bacterium | 0.52 | 0.54 |
|  | Elsterales | URHD0088 | uncultured.Rhodospirillaceae.bacterium |  | 0.51 |
|  | Micropepsales | Micropepsaceae | uncultured | 0.63 | 0.64 |
|  | Micropepsales | Micropepsaceae | uncultured | 0.64 | 0.63 |
|  | Micropepsales | Micropepsaceae | uncultured | 0.63 | 0.51 |
|  | Micropepsales | Micropepsaceae | uncultured | 0.71 | 0.66 |
|  | Micropepsales | Micropepsaceae | uncultured | 0.63 | 0.67 |
|  | Micropepsales | Micropepsaceae | uncultured | 0.52 | 0.50 |
|  | Micropepsales | Micropepsaceae | uncultured | 0.62 | 0.53 |
|  | Micropepsales | Micropepsaceae | uncultured |  | -0.61 |
|  | Micropepsales | Micropepsaceae | uncultured.proteobacterium | 0.63 | 0.58 |
|  | Reyranellales | Reyranellaceae | Reyranella | 0.65 | 0.55 |
|  | Reyranellales | Reyranellaceae | Reyranella | 0.53 | 0.53 |
|  | Reyranellales | Reyranellaceae | uncultured | -0.54 |  |
|  | Rhizobiales | A0839 | uncultured.bacterium | -0.66 | -0.67 |
|  | Rhizobiales | A0839 | uncultured.bacterium |  | -0.59 |
|  | Rhizobiales | Beijerinckiaceae | Bosea | -0.53 |  |
|  | Rhizobiales | Beijerinckiaceae | FFCH5858 | -0.52 |  |
|  | Rhizobiales | Beijerinckiaceae | Methylobacterium.Methylorubrum |  | -0.55 |
|  | Rhizobiales | Beijerinckiaceae | Microvirga | -0.71 | -0.70 |
|  | Rhizobiales | Beijerinckiaceae | Microvirga | -0.51 |  |
|  | Rhizobiales | Beijerinckiaceae | Microvirga | -0.50 | -0.68 |
|  | Rhizobiales | Beijerinckiaceae | Microvirga | -0.72 | -0.65 |
|  | Rhizobiales | Beijerinckiaceae | Psychroglaciecola |  | 0.60 |
|  | Rhizobiales | Beijerinckiaceae | Psychroglaciecola | -0.50 | -0.53 |
|  | Rhizobiales | Beijerinckiaceae | Roseiarcus | 0.63 | 0.64 |
|  | Rhizobiales | Beijerinckiaceae | Roseiarcus | 0.51 | 0.53 |
|  | Rhizobiales | Beijerinckiaceae | uncultured.Beijerinckiaceae.bacterium | -0.55 |  |
|  | Rhizobiales | Hyphomicrobiaceae | Filomicrobium |  | -0.53 |
|  | Rhizobiales | Hyphomicrobiaceae | Hyphomicrobium | -0.60 | -0.60 |
|  | Rhizobiales | Hyphomicrobiaceae | Hyphomicrobium | -0.63 | -0.73 |
|  | Rhizobiales | Hyphomicrobiaceae | Hyphomicrobium |  | -0.53 |
|  | Rhizobiales | Hyphomicrobiaceae | Hyphomicrobium | 0.65 | 0.59 |
|  | Rhizobiales | Hyphomicrobiaceae | Pedomicrobium | 0.70 | 0.72 |
|  | Rhizobiales | Hyphomicrobiaceae | Pedomicrobium | -0.52 | -0.51 |
|  | Rhizobiales | KF.JG30.B3 | metagenome | 0.63 | 0.65 |
|  | Rhizobiales | KF.JG30.B3 | uncultured.Alphaproteobacteria.bacterium | 0.57 | 0.51 |
|  | Rhizobiales | KF.JG30.B3 | uncultured.Alphaproteobacteria.bacterium | -0.50 | -0.53 |
|  | Rhizobiales | KF.JG30.B3 | uncultured.bacterium | 0.55 | 0.53 |
|  | Rhizobiales | Methyloligellaceae | uncultured | -0.67 | -0.62 |
|  | Rhizobiales | Methyloligellaceae | uncultured.Rhodobiaceae.bacterium | 0.66 | 0.64 |
|  | Rhizobiales | Rhizobiaceae | Allorhizobium.Neorhizobium.Pararhizobium.Rhizobium | -0.51 |  |
|  | Rhizobiales | Rhizobiaceae | Allorhizobium.Neorhizobium.Pararhizobium.Rhizobium | -0.54 |  |
|  | Rhizobiales | Rhizobiaceae | Aminobacter |  | -0.60 |
|  | Rhizobiales | Rhizobiaceae | Ensifer | -0.69 | -0.67 |
|  | Rhizobiales | Rhizobiaceae | Mesorhizobium | -0.67 | -0.62 |
|  | Rhizobiales | Rhizobiaceae | Mesorhizobium | -0.53 | -0.55 |
|  | Rhizobiales | Rhizobiaceae | Mesorhizobium | -0.53 |  |
|  | Rhizobiales | Rhizobiales.Incertae.Sedis | Bauldia |  | -0.51 |
|  | Rhizobiales | Rhizobiales.Incertae.Sedis | Bauldia |  | -0.52 |
|  | Rhizobiales | Rhizobiales.Incertae.Sedis | Nordella | -0.67 | -0.62 |
|  | Rhizobiales | Rhizobiales.Incertae.Sedis | Nordella | -0.66 | -0.66 |
|  | Rhizobiales | Rhizobiales.Incertae.Sedis | Nordella | -0.60 | -0.62 |
|  | Rhizobiales | Rhodomicrobiaceae | Rhodomicrobium |  | 0.53 |
|  | Rhizobiales | Rhodomicrobiaceae | Rhodomicrobium | -0.53 |  |
|  | Rhizobiales | uncultured | uncultured.Alphaproteobacteria.bacterium | -0.51 |  |
|  | Rhizobiales | uncultured | uncultured.forest.soil.bacterium | 0.64 | 0.59 |
|  | Rhizobiales | Xanthobacteraceae | Bradyrhizobium | 0.65 | 0.62 |
|  | Rhizobiales | Xanthobacteraceae | Bradyrhizobium | 0.55 | 0.59 |
|  | Rhizobiales | Xanthobacteraceae | Bradyrhizobium | 0.58 | 0.53 |
|  | Rhizobiales | Xanthobacteraceae | Pseudolabrys | 0.74 | 0.77 |
|  | Rhizobiales | Xanthobacteraceae | Pseudolabrys | 0.60 | 0.62 |
|  | Rhizobiales | Xanthobacteraceae | Pseudolabrys | 0.65 | 0.61 |
|  | Rhizobiales | Xanthobacteraceae | Pseudolabrys | 0.68 | 0.72 |
|  | Rhizobiales | Xanthobacteraceae | Pseudorhodoplanes | -0.61 | -0.69 |
|  | Rhizobiales | Xanthobacteraceae | Rhodoplanes | -0.54 |  |
|  | Rhizobiales | Xanthobacteraceae | Rhodoplanes | 0.55 | 0.65 |
|  | Rhizobiales | Xanthobacteraceae | Rhodoplanes | 0.54 | 0.53 |
|  | Rhizobiales | Xanthobacteraceae | uncultured | 0.55 | 0.55 |
|  | Rhizobiales | Xanthobacteraceae | uncultured | 0.68 | 0.66 |
|  | Rhizobiales | Xanthobacteraceae | uncultured | 0.59 | 0.62 |
|  | Rhizobiales | Xanthobacteraceae | uncultured | -0.65 | -0.68 |
|  | Rhizobiales | Xanthobacteraceae | uncultured | 0.56 | 0.62 |
|  | Rhizobiales | Xanthobacteraceae | uncultured | 0.63 | 0.61 |
|  | Rhizobiales | Xanthobacteraceae | uncultured | -0.54 |  |
|  | Rhizobiales | Xanthobacteraceae | uncultured |  | 0.52 |
|  | Rhizobiales | Xanthobacteraceae | uncultured.bacterium | 0.62 | 0.64 |
|  | Rhizobiales | Xanthobacteraceae | uncultured.bacterium | 0.55 |  |
|  | Rhodobacterales | Rhodobacteraceae | Amaricoccus | -0.66 | -0.69 |
|  | Rhodobacterales | Rhodobacteraceae | Amaricoccus | -0.63 | -0.69 |
|  | Rhodobacterales | Rhodobacteraceae | Amaricoccus | -0.54 |  |
|  | Rhodobacterales | Rhodobacteraceae | uncultured | -0.63 | -0.67 |
|  | Rhodobacterales | Rhodobacteraceae | uncultured | -0.52 | -0.55 |
|  | Rhodospirillales | uncultured | uncultured.proteobacterium | 0.53 | 0.53 |
|  | Rickettsiales | Anaplasmataceae | Candidatus.Neoehrlichia.sp..HT.IT.4 | 0.59 | 0.61 |
|  | Sphingomonadales | Sphingomonadaceae | Altererythrobacter | -0.53 |  |
|  | Sphingomonadales | Sphingomonadaceae | Altererythrobacter |  | 0.51 |
|  | Sphingomonadales | Sphingomonadaceae | Altererythrobacter | -0.50 |  |
|  | Sphingomonadales | Sphingomonadaceae | Ellin6055 |  | 0.57 |
|  | Sphingomonadales | Sphingomonadaceae | Novosphingobium |  | -0.58 |
|  | Sphingomonadales | Sphingomonadaceae | Sphingomonas | -0.57 |  |
|  | Sphingomonadales | Sphingomonadaceae | Sphingomonas | -0.69 | -0.59 |
|  | Sphingomonadales | Sphingomonadaceae | Sphingomonas | -0.65 | -0.59 |
|  | Sphingomonadales | Sphingomonadaceae | Sphingomonas | -0.51 |  |
|  | Sphingomonadales | Sphingomonadaceae | uncultured | -0.67 | -0.63 |
|  | Tistrellales | Geminicoccaceae | Candidatus.Alysiosphaera | -0.61 | -0.56 |
|  | Tistrellales | Geminicoccaceae | Candidatus.Alysiosphaera | -0.71 | -0.68 |
|  | Tistrellales | Geminicoccaceae | Candidatus.Alysiosphaera | -0.71 | -0.66 |
|  | Tistrellales | Geminicoccaceae | Candidatus.Alysiosphaera | -0.57 | -0.51 |
|  | Tistrellales | Geminicoccaceae | Candidatus.Alysiosphaera |  | -0.51 |
|  | Tistrellales | Geminicoccaceae | Geminicoccus | -0.58 | -0.62 |
|  | uncultured | uncultured.bacterium | NA | 0.70 | 0.63 |
|  | uncultured | uncultured.bacterium | NA | -0.75 | -0.71 |
|  | uncultured | uncultured.bacterium | NA | -0.63 | -0.58 |
|  | uncultured | uncultured.bacterium | NA | -0.51 | -0.59 |
|  | uncultured | uncultured.bacterium | NA |  | -0.52 |
|  | Burkholderiales | A21b | uncultured.bacterium |  | 0.50 |
|  | Burkholderiales | A21b | uncultured.bacterium | 0.69 | 0.64 |
|  | Burkholderiales | A21b | uncultured.beta.proteobacterium | 0.55 | 0.55 |
|  | Burkholderiales | A21b | uncultured.Burkholderiaceae.bacterium | 0.74 | 0.76 |
|  | Burkholderiales | Burkholderiaceae | Burkholderia.Caballeronia.Paraburkholderia | 0.62 | 0.64 |
|  | Burkholderiales | Burkholderiaceae | Burkholderia.Caballeronia.Paraburkholderia |  | 0.52 |
|  | Burkholderiales | Burkholderiaceae | Burkholderia.Caballeronia.Paraburkholderia | -0.56 | -0.57 |
|  | Burkholderiales | Burkholderiaceae | Cupriavidus |  | -0.51 |
|  | Burkholderiales | Burkholderiaceae | Lautropia | -0.54 | -0.53 |
|  | Burkholderiales | Comamonadaceae | Ideonella | -0.66 | -0.62 |
|  | Burkholderiales | Comamonadaceae | Leptothrix | -0.59 |  |
|  | Burkholderiales | Comamonadaceae | Polaromonas | -0.57 | -0.57 |
|  | Burkholderiales | Comamonadaceae | Rhizobacter | -0.66 | -0.63 |
|  | Burkholderiales | Comamonadaceae | uncultured |  | 0.59 |
|  | Burkholderiales | Comamonadaceae | uncultured | -0.56 | -0.51 |
|  | Burkholderiales | Methylophilaceae | Methylotenera |  | -0.60 |
|  | Burkholderiales | Methylophilaceae | Methylotenera |  | -0.59 |
|  | Burkholderiales | Methylophilaceae | uncultured |  | -0.54 |
|  | Burkholderiales | Nitrosomonadaceae | Ellin6067 | -0.55 | -0.53 |
|  | Burkholderiales | Nitrosomonadaceae | Ellin6067 | 0.63 | 0.63 |
|  | Burkholderiales | Nitrosomonadaceae | Ellin6067 | -0.75 | -0.66 |
|  | Burkholderiales | Nitrosomonadaceae | Ellin6067 |  | -0.51 |
|  | Burkholderiales | Nitrosomonadaceae | Ellin6067 |  | -0.52 |
|  | Burkholderiales | Nitrosomonadaceae | GOUTA6 | 0.51 | 0.51 |
|  | Burkholderiales | Nitrosomonadaceae | IS.44 | -0.67 | -0.70 |
|  | Burkholderiales | Nitrosomonadaceae | IS.44 | -0.54 | -0.58 |
|  | Burkholderiales | Nitrosomonadaceae | mle1.7 | -0.63 | -0.69 |
|  | Burkholderiales | Nitrosomonadaceae | mle1.7 | -0.65 | -0.65 |
|  | Burkholderiales | Nitrosomonadaceae | MND1 | 0.51 |  |
|  | Burkholderiales | Nitrosomonadaceae | MND1 | 0.61 | 0.57 |
|  | Burkholderiales | Nitrosomonadaceae | MND1 | -0.56 | -0.52 |
|  | Burkholderiales | Nitrosomonadaceae | MND1 | -0.57 | -0.52 |
|  | Burkholderiales | Nitrosomonadaceae | MND1 | -0.74 | -0.72 |
|  | Burkholderiales | Nitrosomonadaceae | MND1 | -0.55 |  |
|  | Burkholderiales | Nitrosomonadaceae | MND1 | -0.56 | -0.58 |
|  | Burkholderiales | Nitrosomonadaceae | MND1 | -0.59 | -0.55 |
|  | Burkholderiales | Oxalobacteraceae | Massilia | 0.52 |  |
|  | Burkholderiales | Oxalobacteraceae | Massilia | 0.57 | 0.53 |
|  | Burkholderiales | Oxalobacteraceae | Massilia | -0.53 | -0.60 |
|  | Burkholderiales | Oxalobacteraceae | Noviherbaspirillum | 0.66 | 0.68 |
|  | Burkholderiales | SC.I.84 | metagenome |  | -0.52 |
|  | Burkholderiales | SC.I.84 | uncultured.bacterium | -0.64 | -0.65 |
|  | Burkholderiales | SC.I.84 | uncultured.bacterium | -0.50 |  |
|  | Burkholderiales | SC.I.84 | uncultured.bacterium | -0.53 |  |
|  | Burkholderiales | SC.I.84 | uncultured.bacterium | -0.58 | -0.53 |
|  | Burkholderiales | SC.I.84 | uncultured.beta.proteobacterium | -0.57 | -0.63 |
|  | Burkholderiales | SC.I.84 | uncultured.beta.proteobacterium | 0.63 | 0.67 |
|  | Burkholderiales | SC.I.84 | uncultured.soil.bacterium | 0.57 | 0.56 |
|  | Burkholderiales | Sutterellaceae | uncultured | -0.51 | -0.56 |
|  | Burkholderiales | TRA3.20 | uncultured.bacterium | -0.66 | -0.60 |
|  | CCD24 | metagenome | NA | 0.51 |  |
|  | Enterobacterales | Morganellaceae | endosymbionts |  | 0.52 |
|  | Gammaproteobacteria.Incertae.Sedis | Unknown.Family | Acidibacter | 0.66 | 0.67 |
|  | Gammaproteobacteria.Incertae.Sedis | Unknown.Family | Acidibacter | 0.58 | 0.52 |
|  | Gammaproteobacteria.Incertae.Sedis | Unknown.Family | Acidibacter | 0.59 | 0.66 |
|  | Gammaproteobacteria.Incertae.Sedis | Unknown.Family | Acidibacter | 0.55 |  |
|  | Gammaproteobacteria.Incertae.Sedis | Unknown.Family | Acidibacter | 0.60 |  |
|  | JG36.TzT.191 | uncultured.bacterium | NA | 0.67 | 0.65 |
|  | JG36.TzT.191 | uncultured.bacterium | NA | 0.59 | 0.64 |
|  | JG36.TzT.191 | uncultured.bacterium | NA | 0.58 | 0.57 |
|  | JG36.TzT.191 | uncultured.bacterium | NA | 0.56 | 0.53 |
|  | PLTA13 | metagenome | NA | -0.53 | -0.51 |
|  | Pseudomonadales | Halieaceae | OM60.NOR5..clade | -0.53 | -0.53 |
|  | Pseudomonadales | Pseudohongiellaceae | BIyi10 |  | -0.50 |
|  | R7C24 | metagenome | NA |  | 0.58 |
|  | Salinisphaerales | Solimonadaceae | Fontimonas | -0.65 | -0.68 |
|  | Salinisphaerales | Solimonadaceae | Polycyclovorans | -0.71 | -0.73 |
|  | Salinisphaerales | Solimonadaceae | Polycyclovorans | -0.66 | -0.63 |
|  | Salinisphaerales | Solimonadaceae | Polycyclovorans | -0.66 | -0.57 |
|  | Salinisphaerales | Solimonadaceae | Polycyclovorans | -0.53 |  |
|  | Salinisphaerales | Solimonadaceae | Polycyclovorans | -0.67 | -0.62 |
|  | Salinisphaerales | Solimonadaceae | Polycyclovorans |  | -0.50 |
|  | Salinisphaerales | Solimonadaceae | uncultured | -0.53 |  |
|  | Steroidobacterales | Steroidobacteraceae | Steroidobacter |  | -0.53 |
|  | Steroidobacterales | Steroidobacteraceae | Steroidobacter | -0.56 | -0.60 |
|  | Steroidobacterales | Steroidobacteraceae | Steroidobacter | 0.55 |  |
|  | Steroidobacterales | Steroidobacteraceae | uncultured | -0.54 | -0.50 |
|  | Steroidobacterales | Steroidobacteraceae | uncultured | -0.53 | -0.51 |
|  | Steroidobacterales | Steroidobacteraceae | uncultured | -0.56 | -0.51 |
|  | Steroidobacterales | Steroidobacteraceae | uncultured | 0.55 |  |
|  | WD260 | uncultured.bacterium | NA | 0.77 | 0.77 |
|  | Xanthomonadales | Rhodanobacteraceae | Ahniella | -0.52 | -0.53 |
|  | Xanthomonadales | Rhodanobacteraceae | Ahniella | -0.59 | -0.59 |
|  | Xanthomonadales | Rhodanobacteraceae | Dokdonella | 0.62 | 0.55 |
|  | Xanthomonadales | Rhodanobacteraceae | Rhodanobacter | 0.55 |  |
|  | Xanthomonadales | Rhodanobacteraceae | Rhodanobacter | 0.55 | 0.62 |
|  | Xanthomonadales | Rhodanobacteraceae | Rhodanobacter |  | 0.50 |
|  | Xanthomonadales | Xanthomonadaceae | Lysobacter | 0.57 | 0.66 |
|  | Xanthomonadales | Xanthomonadaceae | Lysobacter | -0.67 | -0.54 |
|  | Xanthomonadales | Xanthomonadaceae | Lysobacter | -0.71 | -0.71 |
|  | Xanthomonadales | Xanthomonadaceae | Lysobacter |  | -0.56 |
|  | Xanthomonadales | Xanthomonadaceae | Lysobacter | -0.54 | -0.51 |
|  | Xanthomonadales | Xanthomonadaceae | uncultured | 0.54 | 0.51 |
|  | Xanthomonadales | Xanthomonadaceae | uncultured | 0.51 | 0.53 |
| RCP2.54 | NA | NA | NA | 0.66 | 0.66 |
|  | NA | NA | NA | 0.51 |  |
|  | NA | NA | NA | 0.70 | 0.72 |
|  | NA | NA | NA | 0.69 | 0.67 |
|  | NA | NA | NA | 0.67 | 0.66 |
|  | NA | NA | NA | -0.59 | -0.56 |
| Verrucomicrobiota | Chthoniobacterales | Chthoniobacteraceae | Candidatus.Udaeobacter | 0.54 |  |
|  | Chthoniobacterales | Chthoniobacteraceae | Candidatus.Udaeobacter | -0.52 |  |
|  | Chthoniobacterales | Chthoniobacteraceae | Candidatus.Udaeobacter | -0.53 |  |
|  | Chthoniobacterales | Chthoniobacteraceae | Chthoniobacter | -0.63 | -0.69 |
|  | Chthoniobacterales | Chthoniobacteraceae | Chthoniobacter | -0.50 |  |
|  | Chthoniobacterales | Chthoniobacteraceae | Chthoniobacter | -0.57 | -0.59 |
|  | Chthoniobacterales | Chthoniobacteraceae | Chthoniobacter | -0.51 | -0.50 |
|  | Chthoniobacterales | Chthoniobacteraceae | Chthoniobacter | -0.53 |  |
|  | Chthoniobacterales | Chthoniobacteraceae | Chthoniobacter | -0.59 | -0.57 |
|  | Chthoniobacterales | Chthoniobacteraceae | Chthoniobacter | -0.60 | -0.67 |
|  | Chthoniobacterales | Chthoniobacteraceae | Chthoniobacter |  | -0.56 |
|  | Chthoniobacterales | Xiphinematobacteraceae | Candidatus.Xiphinematobacter | 0.52 | 0.52 |
|  | Chthoniobacterales | Xiphinematobacteraceae | Candidatus.Xiphinematobacter | -0.57 | -0.64 |
|  | Pedosphaerales | Pedosphaeraceae | ADurb.Bin063.1 | 0.55 | 0.51 |
|  | Pedosphaerales | Pedosphaeraceae | ADurb.Bin063.1 | 0.60 | 0.54 |
|  | Pedosphaerales | Pedosphaeraceae | ADurb.Bin063.1 | 0.51 |  |
|  | Pedosphaerales | Pedosphaeraceae | Ellin516 | 0.60 | 0.56 |
|  | Pedosphaerales | Pedosphaeraceae | metagenome | -0.53 | -0.64 |
|  | Pedosphaerales | Pedosphaeraceae | uncultured | 0.51 | 0.55 |
|  | Pedosphaerales | Pedosphaeraceae | uncultured | 0.60 | 0.54 |
|  | Pedosphaerales | Pedosphaeraceae | uncultured |  | -0.52 |
|  | Pedosphaerales | Pedosphaeraceae | uncultured.bacterium | 0.50 |  |
|  | Pedosphaerales | Pedosphaeraceae | uncultured.bacterium |  | -0.54 |
|  | Pedosphaerales | Pedosphaeraceae | uncultured.bacterium | -0.56 | -0.55 |
|  | Pedosphaerales | Pedosphaeraceae | uncultured.bacterium | -0.51 | -0.51 |
|  | Verrucomicrobiales | Verrucomicrobiaceae | Roseimicrobium | -0.66 | -0.66 |
|  | Verrucomicrobiales | Verrucomicrobiaceae | uncultured | -0.54 | -0.61 |
|  | Verrucomicrobiales | Verrucomicrobiaceae | uncultured | -0.56 | -0.60 |
|  | Verrucomicrobiales | Verrucomicrobiaceae | uncultured |  | -0.53 |
| WPS.2 | NA | NA | NA | 0.55 |  |
|  | NA | NA | NA | -0.56 | -0.54 |
|  | NA | NA | NA | 0.68 | 0.68 |
|  | NA | NA | NA | 0.56 | 0.59 |
|  | NA | NA | NA | 0.53 |  |
|  | NA | NA | NA | 0.60 |  |
|  | NA | NA | NA | 0.59 |  |
|  | NA | NA | NA | 0.53 |  |
|  | NA | NA | NA | 0.59 | 0.60 |
| Zixibacteria | NA | NA | NA | -0.52 | -0.53 |

**Supplementary table 4** **Correlation analysis between soil OM% and fungal OTU abundance based on GMPR-normalized data. Spearman correlation coefficients calculated separately for rhizosphere and bulk soil. Table shows moderate and strong correlations |>0.5|, P<0.05.**

| phylum | family | genus | OTU | soil | Ritso |
| --- | --- | --- | --- | --- | --- |
| p__Aphelidiomycota | f__unidentified | g__unidentified | GS16_sp_SH1571569_Aphel_118 | -0.56051 |  |
| p__Ascomycota | f__Archaeorhizomycetaceae | g__Archaeorhizomyces | Archaeorhizomyces_sp_SH1571534_Ascom_332 | 0.53093 |  |
|  | f__Didymellaceae | g__Didymella | **Didymella_aurea_SH2232261_Ascom_60979** | -0.66557 | -0.68057 |
|  | f__Didymellaceae | g__Didymella | Didymella_protuberans_SH2232168_Ascom_502 | -0.61387 | -0.58161 |
|  | f__Phaeosphaeriaceae | g__Paraphoma | Paraphoma_chrysanthemicola_SH1576212_Ascom_8335 |  | -0.5603 |
|  | f__unidentified | g__unidentified | Pleosporales_sp_SH1527184_Ascom_4800 | -0.56475 | -0.60646 |
|  | f__unidentified | g__unidentified | Pleosporales_sp_SH1649808_Ascom_1609 | -0.52054 | -0.55525 |
|  | f__Herpotrichiellaceae | g__Exophiala | **Exophiala_radicis_SH2083560_Ascom_28089** | -0.72453 | -0.77938 |
|  | f__Aspergillaceae | g__Penicillium | Penicillium_adametzii_SH1535734_Ascom_2356 | 0.642881 | 0.591223 |
|  | f__Aspergillaceae | g__Penicillium | Penicillium_simplicissimum_SH1896005_Ascom_2419 | 0.630489 | 0.543146 |
|  | f__Dermateaceae | g__Mollisia | **Mollisia_sp_SH1524858_Ascom_3815** | 0.726588 | 0.645629 |
|  | f__Dermateaceae | g__Pyrenopeziza | Pyrenopeziza_dilutella_SH1545977_Ascom_2127 | -0.58225 | -0.56907 |
|  | f__Helotiaceae | g__Collophora | Collophora_sp_SH1647638_Ascom_483 | -0.59046 | -0.57524 |
|  | f__Helotiaceae | g__Glarea | Glarea_lozoyensis_SH1543039_Ascom_5905 | 0.528027 |  |
|  | f__Helotiaceae | g__Mycosymbioces | **Mycosymbioces_sp_SH1509588_Ascom_10487** | 0.720487 | 0.675647 |
|  | f__Helotiaceae | g__Tetracladium | Tetracladium_sp_SH1648798_Ascom_19405 | -0.5246 | -0.51254 |
|  | f__Helotiaceae | g__unidentified | **Helotiaceae_sp_SH1523512_Ascom_23486** | 0.682152 | 0.648337 |
|  | f__Helotiales_fam_Incertae_sedis | g__Malotium | Malotium_paludosum_SH2713370_Ascom_344 | -0.51546 |  |
|  | f__Hyaloscyphaceae | g__Clathrosphaerina | Clathrosphaerina_zalewskii_SH1566287_Ascom_1246 | 0.586533 |  |
|  | f__Hyaloscyphaceae | g__Hyaloscypha | Hyaloscypha_sp_SH1506161_Ascom_7982 |  | -0.51606 |
|  | f__Myxotrichaceae | g__Oidiodendron | Oidiodendron_echinulatum_SH1564457_Ascom_921 | 0.512262 |  |
|  | f__Sclerotiniaceae | g__Botrytis | Botrytis_californica_SH2311583_Ascom_179320 |  | 0.610059 |
|  | f__unidentified | g__unidentified | Helotiales_sp_SH1522489_Ascom_2643 | 0.574392 | 0.640558 |
|  | f__unidentified | g__unidentified | **Helotiales_sp_SH1545902_Ascom_5079** | 0.759143 | 0.704014 |
|  | f__unidentified | g__unidentified | Helotiales_sp_SH1648839_Ascom_4804 | -0.51217 | -0.5181 |
|  | f__unidentified | g__unidentified | Helotiales_sp_SH1648870_Ascom_19849 | -0.5419 |  |
|  | f__unidentified | g__unidentified | Helotiales_sp_SH2723084_Ascom_1040 | 0.633009 | 0.534093 |
|  | f__unidentified | g__unidentified | Leotiomycetes_sp_SH2721119_Ascom_488 | 0.501736 |  |
|  | f__unidentified | g__unidentified | Leotiomycetes_sp_SH2721720_Ascom_2583 | 0.590738 | 0.585707 |
|  | f__Orbiliaceae | g__Arthrobotrys | Arthrobotrys_xiangyunensis_SH1570340_Ascom_341 |  | -0.54652 |
|  | f__Coniochaetaceae | g__Coniochaeta | **Coniochaeta_sp_SH1645136_Ascom_2134** | 0.700124 | 0.637734 |
|  | f__Coniochaetaceae | g__Lecythophora | Lecythophora_sp_SH1645147_Ascom_2663 | -0.52323 | -0.53261 |
|  | f__unidentified | g__unidentified | Coniochaetales_sp_SH1645163_Ascom_351 | -0.5872 |  |
|  | f__Clavicipitaceae | g__Metarhizium | **Metarhizium_marquandii_SH1561418_Ascom_9672** | -0.72112 | -0.68348 |
|  | f__Hypocreales_fam_Incertae_sedis | g__Emericellopsis | Emericellopsis_minima_SH2008692_Ascom_1413 | -0.5784 | -0.58188 |
|  | f__Nectriaceae | g__Fusicolla | **Fusicolla_aquaeductuum_SH1546339_Ascom_6765** | 0.730078 | 0.695967 |
|  | f__Nectriaceae | g__unidentified | Nectriaceae_sp_SH1546526_Ascom_16240 | 0.577067 | 0.648071 |
|  | f__Nectriaceae | g__unidentified | **Nectriaceae_sp_SH1653658_Ascom_1232** | -0.6725 | -0.68172 |
|  | f__Ophiocordycipitaceae | g__Haptocillium | Haptocillium_sp_SH2724229_Ascom_261 | 0.527148 |  |
|  | f__unidentified | g__unidentified | Hypocreales_sp_SH1651746_Ascom_914 | -0.56545 | -0.56369 |
|  | f__Microascaceae | g__Pseudallescheria | Pseudallescheria_boydii_SH2328456_Ascom_576 | -0.56554 |  |
|  | f__Microascales_fam_Incertae_sedis | g__Wardomyces | Wardomyces_inflatus_SH1575618_Ascom_641 | -0.51257 |  |
|  | f__Chaetomiaceae | g__Arxotrichum | Arxotrichum_wyomingense_SH1800557_Ascom_1971 | 0.511865 | 0.573811 |
|  | f__Lasiosphaeriaceae | g__Podospora | Podospora_sp_SH1567848_Ascom_9312 | 0.621137 | 0.575716 |
|  | f__Lasiosphaeriaceae | g__unidentified | Lasiosphaeriaceae_sp_SH1506805_Ascom_45537 |  | 0.510904 |
|  | f__Lasiosphaeriaceae | g__unidentified | Lasiosphaeriaceae_sp_SH1567857_Ascom_5923 | 0.633351 | 0.59828 |
|  | f__Sordariales_fam_Incertae_sedis | g__Ramophialophora | Ramophialophora_humicola_SH1615709_Ascom_571 | -0.51328 | -0.50844 |
|  | f__unidentified | g__unidentified | Sordariales_sp_SH1555263_Ascom_2078 | 0.613918 | 0.594 |
|  | f__unidentified | g__unidentified | Sordariomycetes_sp_SH1615621_Ascom_873 | -0.52682 | -0.54947 |
|  | f__unidentified | g__unidentified | Ascomycota_sp_SH1648788_Ascom_54217 | -0.6303 | -0.59099 |
| p__Basidiomycota | f__Typhulaceae | g__Typhula | Typhula_incarnata_SH1525687_Basid_990 |  | -0.54865 |
|  | f__unidentified | g__unidentified | Agaricales_sp_SH1574424_Basid_17540 |  | 0.506252 |
|  | f__unidentified | g__unidentified | Agaricales_sp_SH2720470_Basid_504 | 0.544535 |  |
|  | f__unidentified | g__unidentified | Agaricales_sp_SH2748586_Basid_891 | 0.526304 |  |
|  | f__Ceratobasidiaceae | g__Thanatephorus | Thanatephorus_cucumeris_SH1522029_Basid_3241 |  | -0.59656 |
|  | f__Schizoporaceae | g__Hyphodontia | Hyphodontia_sp_SH1522035_Basid_2354 | -0.55786 | -0.5852 |
|  | f__Erythrobasidiales_fam_Incertae_sedis | g__Sakaguchia | Sakaguchia_meli_SH1797925_Basid_832 |  | -0.56228 |
|  | f__Chrysozymaceae | g__Sampaiozyma | Sampaiozyma_ingeniosa_SH2108419_Basid_12555 |  | -0.53637 |
|  | f__Chrysozymaceae | g__Slooffia | Slooffia_cresolica_SH1515292_Basid_4228 | 0.617797 | 0.560947 |
|  | f__Piskurozymaceae | g__Solicoccozyma | Solicoccozyma_terricola_SH1649268_Basid_340797 | 0.579824 | 0.601801 |
|  | f__Bulleribasidiaceae | g__Dioszegia | Dioszegia_butyracea_SH1609792_Basid_898 |  | -0.5202 |
|  | f__Trimorphomycetaceae | g__Saitozyma | **Saitozyma_podzolica_SH1565595_Basid_118490** | 0.758444 | 0.751584 |
|  | f__unidentified | g__unidentified | Tremellales_sp_SH2737670_Basid_1465 | 0.568878 |  |
|  | f__Tetragoniomycetaceae | g__unidentified | Tetragoniomycetaceae_sp_SH2720338_Basid_11266 | 0.553217 | 0.51925 |
|  | f__unidentified | g__unidentified | Tremellomycetes_sp_SH1557488_Basid_46725 | 0.512363 | 0.557623 |
|  | f__unidentified | g__unidentified | Basidiomycota_sp_SH2748115_Basid_4518 | 0.547753 |  |
| p__Chytridiomycota | f__Rhizophlyctidaceae | g__Rhizophlyctis | Rhizophlyctis_rosea_SH1653265_Chytr_3570 | -0.53116 | -0.53526 |
|  | f__unidentified | g__unidentified | Rhizophydiales_sp_SH3331458_Chytr_1066 |  | -0.56966 |
|  | f__Powellomycetaceae | g__Fimicolochytrium | Fimicolochytrium_jonesii_SH1570166_Chytr_9402 | -0.50078 |  |
|  | f__Powellomycetaceae | g__unidentified | Powellomycetaceae_sp_SH2720249_Chytr_1214 | -0.53315 |  |
|  | f__Powellomycetaceae | g__unidentified | **Powellomycetaceae_sp_SH2720457_Chytr_4635** | 0.778834 | 0.766387 |
|  | f__Spizellomycetaceae | g__Spizellomyces | Spizellomyces_sp_SH2715388_Chytr_3314 |  | 0.506816 |
|  | f__Spizellomycetaceae | g__Spizellomyces | Spizellomyces_sp_SH2720877_Chytr_2113 | 0.533527 |  |
|  | f__Spizellomycetaceae | g__unidentified | Spizellomycetaceae_sp_SH2722637_Chytr_3727 | 0.544421 | 0.507495 |
|  | f__unidentified | g__unidentified | **Spizellomycetales_sp_SH3322078_Chytr_14758** | 0.692854 | 0.661589 |
|  | f__unidentified | g__unidentified | Chytridiomycota_sp_SH2716664_Chytr_6149 | 0.590574 | 0.551042 |
| p__Glomeromycota | f__Claroideoglomeraceae | g__Claroideoglomus | Claroideoglomus_sp_SH1570676_Glome_953 | -0.50971 |  |
|  | f__Glomeraceae | g__unidentified | Glomeraceae_sp_SH1547694_Glome_4891 | 0.55084 | 0.584327 |
|  | f__Paraglomeraceae | g__Paraglomus | Paraglomus_laccatum_SH1513315_Glome_3880 | 0.529766 | 0.594293 |
| p__Monoblepharomycota | f__Harpochytriaceae | g__unidentified | Harpochytriaceae_sp_SH3339798_Monob_3854 | 0.528873 | 0.515929 |
| p__Mortierellomycota | f__Mortierellaceae | g__Mortierella | Mortierella_basiparvispora_SH1629830_Morti_6886 | 0.561192 | 0.533928 |
|  | f__Mortierellaceae | g__Mortierella | Mortierella_elongata_SH1938497_Morti_208813 | 0.638189 | 0.636042 |
|  | f__Mortierellaceae | g__Mortierella | **Mortierella_horticola_SH2444331_Morti_12455** | 0.653578 | 0.636768 |
|  | f__Mortierellaceae | g__Mortierella | Mortierella_humilis_SH1607997_Morti_82102 | 0.5792 | 0.544689 |
|  | f__Mortierellaceae | g__Mortierella | Mortierella_sp_SH1557076_Morti_2780 | -0.58567 | -0.59463 |
|  | f__Mortierellaceae | g__Mortierella | Mortierella_sp_SH1608167_Morti_6104 | 0.59332 | 0.559217 |
|  | f__Mortierellaceae | g__Mortierella | Mortierella_sp_SH1650292_Morti_5940 |  | -0.50855 |
|  | f__Mortierellaceae | g__unidentified | **Mortierellaceae_sp_SH1650296_Morti_37997** | -0.79067 | -0.80921 |
| p__Mucoromycota | f__Umbelopsidaceae | g__Umbelopsis | **Umbelopsis_vinacea_SH1522250_Mucor_15916** | 0.733077 | 0.721739 |
|  | f__Umbelopsidaceae | g__Umbelopsis | **Umbelopsis_vinacea_SH1522267_Mucor_14701** | 0.678562 | 0.682514 |
| p__unidentified | f__unidentified | g__unidentified | Fungi_sp_SH3315124_unide_90158 |  | 0.562199 |
|  | f__unidentified | g__unidentified | Fungi_sp_SH3337234_unide_1729 | 0.566047 |  |
